# Postsynaptic frequency filters shaped by the interplay of synaptic short-term plasticity and cellular time scales

**DOI:** 10.1101/2023.07.02.547439

**Authors:** Yugarshi Mondal, Guillermo Villanueva Benito, Rodrigo F. O. Pena, Horacio G. Rotstein

## Abstract

Neuronal frequency filters can be thought of as constituent building blocks underlying the ability of neuronal systems to process information, generate rhythms and perform computations. How neuronal filters are generated by the concerted activity of a multiplicity of processes (e.g., electric circuit, history-dependent) and interacting time scales within and across levels of neuronal network organization is poorly understood. In this paper, we use mathematical modeling, numerical simulations and analytical calculations of the postsynaptic response to presynaptic spike trains to address this issue in a basic feedforward network motif in the presence of synaptic short-term plasticity (STP, depression and facilitation). The network motif consists of a presynaptic spike-train, a postsynaptic passive cell, and an excitatory (AMPA) chemical synapse. The dynamics of each network component are controlled by one or more time scales. We explain the mechanisms by which the participating time scales shape the neuronal filters at the (i) synaptic update level (the target of the synaptic variable in response to presynaptic spikes), which is shaped by STP, (ii) the synaptic level, and (iii) the postsynaptic membrane potential (PSP) level. We focus on three metrics that gives rise to three types of profiles (curves of the corresponding metrics as a function of the spike-train input frequency or firing rate): (i) peak profiles, (ii) peak-to-trough amplitude profiles, and (iii) phase profiles. The effects of STP are present at the synaptic update level and are communicated to the synaptic level where they interact with the synaptic time scales. The PSP filters result from the interaction between these variables and time scales and the biophysical properties and time scales of the postsynaptic cell. Band-pass filters (BPFs) result from a combination of low-pass filters (LPFs) and high-pass filters (HPFs) operating at the same or different levels of organization. PSP BPFs can be inherited from the synaptic level (STP-mediated BPFs) or they can be generated across levels of organization due to the interaction between (i) a synaptic LPF and the PSP summation-mediated HPF (PSP peaks), and (ii) a synaptic HPF and the PSP summation-mediated LPF (PSP amplitude). These types of BPFs persist in response to more realistic presynaptic spike trains: jittered (randomly perturbed) periodic spike trains and Poisson-distributed spike trains. The response variability is frequency-dependent and is controlled by STP in a non-monotonic frequency manner. The results and and lessons learned from the investigation of this basic network motif are a necessary step for the construction of a framework to analyze the mechanisms of generation of neuronal filters in networks with more complex architectures and a variety of interacting cellular, synaptic and plasticity time scales.

## 1 Introduction

Neuronal filters allow neuronal systems to select certain information or enhance the communication of specific information components over others [1–6]. Owing to this, neuronal filters play important roles in neuronal information processing, rhythm generation and brain computations [3, 4, 7–18]. Band-pass frequency-filters are associated to the notion of neuronal resonance [1, 4, 19], which has been observed at various levels of neuronal organization, from the subthreshold membrane potential to the network levels [4] (Fig. 1). Neuronal resonance refers to the ability of a neuronal system to exhibit a maximal response (e.g., subthreshold membrane potential, postsynaptic potential, firing rate) to periodic inputs at a preferred (resonant), non-zero frequency band (Fig. 2-A). At the cellular level, in response to oscillatory (sinusoidal) inputs, frequency-filters reflect the time scales of the participating currents [1, 20, 21]. At the synaptic level, in response to presynaptic spike inputs, frequency-filters [22] reflect the synaptic rise and decay times. The observed postsynaptic responses reflect the combination of these and the time scales of the postsynaptic cell’s participating currents [23], which may give rise to additional filtering components resulting from the summation phenomenon [24]. In synaptic pairs with more complex synaptic dynamics, frequency-filters also reflect the time scales associated with synaptic short-term plasticity (STP; depression and facilitation) and may give rise to band-pass filters (BPFs) at the synaptic and postsynaptic levels [2–4, 23, 25–27]. How the concerted activity of this multiplicity of time scales shape neuronal filters is poorly understood.

**Figure 1.**
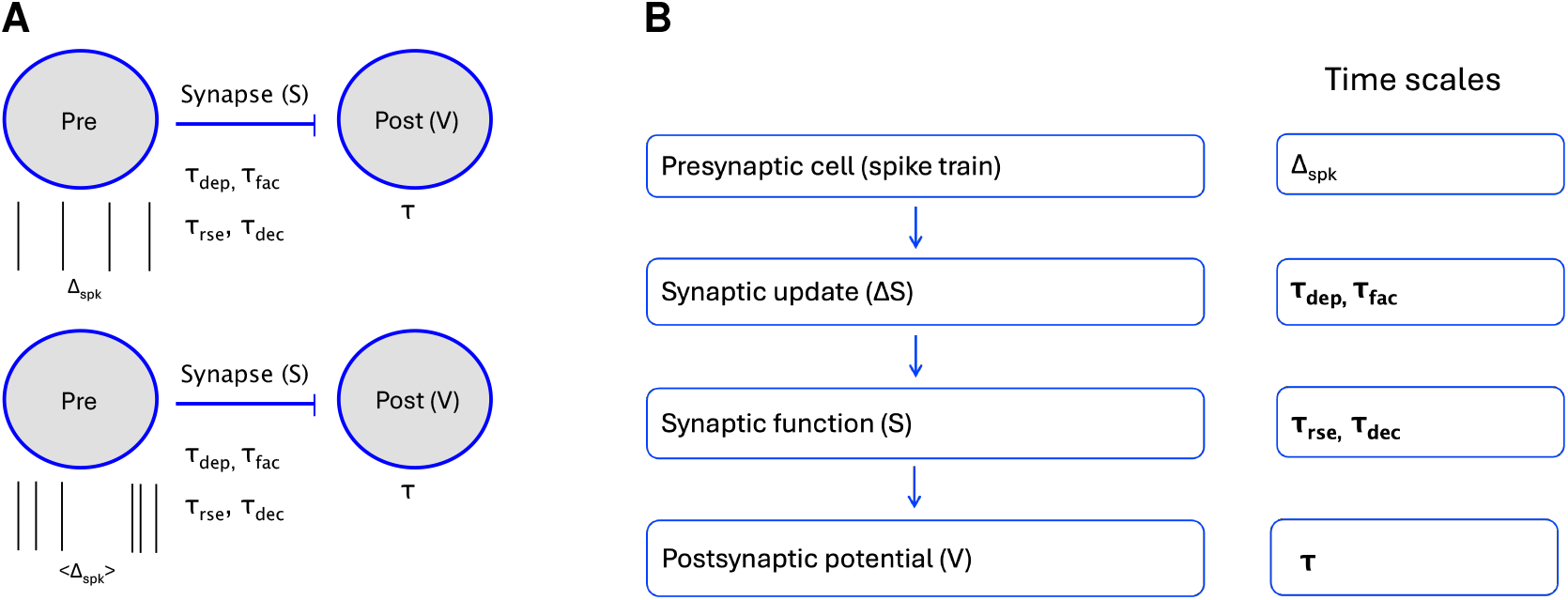
Network motif: Feedforward excitation in the presence of short-term synaptic plasticity. **A**. The presynaptic cell (Pre) is modeled as a spike train either periodic (period Δ_*spk*_, top), jittered-periodic or Poisson distributed (mean interspike interval *<* Δ_*spk*_ *>*, bottom). The postsynaptic cell (Post, membrane potential *V*) is modeled as a passive cell (capacitive and leak currents) with a membrane time constant *τ*. The excitatory synaptic variable (*S*) raises and decays with time constants *τ*_*rse*_ and *τ*_*dec*_, respectively. The synaptic depression and facilitation time constants are *τ*_*dep*_ and *τ*_*fac*_, respectively. **B**. We divide the study of the generation of postsynaptic membrane potential (PSP) filters in response presynaptic inputs within a range of (input) spiking frequencies (periodic or randomly-distributed) in three steps: (i) The Δ*S*-filters generated in response to the presynaptic spike trains are shaped by the synaptic depression and facilitation time constants *τ*_*dep*_ and *τ*_*fac*_, respectively (e.g., Fig. 3, left columns), (ii) The synaptic *S*-filters generated as the result of the Δ*S*-filters are further shaped by the synaptic rise and decay times *τ*_*rse*_ and *τ*_*dec*_, respectively (e.g., Fig. 3, middle columns), and (iii) the PSP filters generated in response to the *S*-filters are further shaped by the postsynaptic membrane potential time constant *τ* (e.g., Fig. 3, right columns).

**Figure 2.**
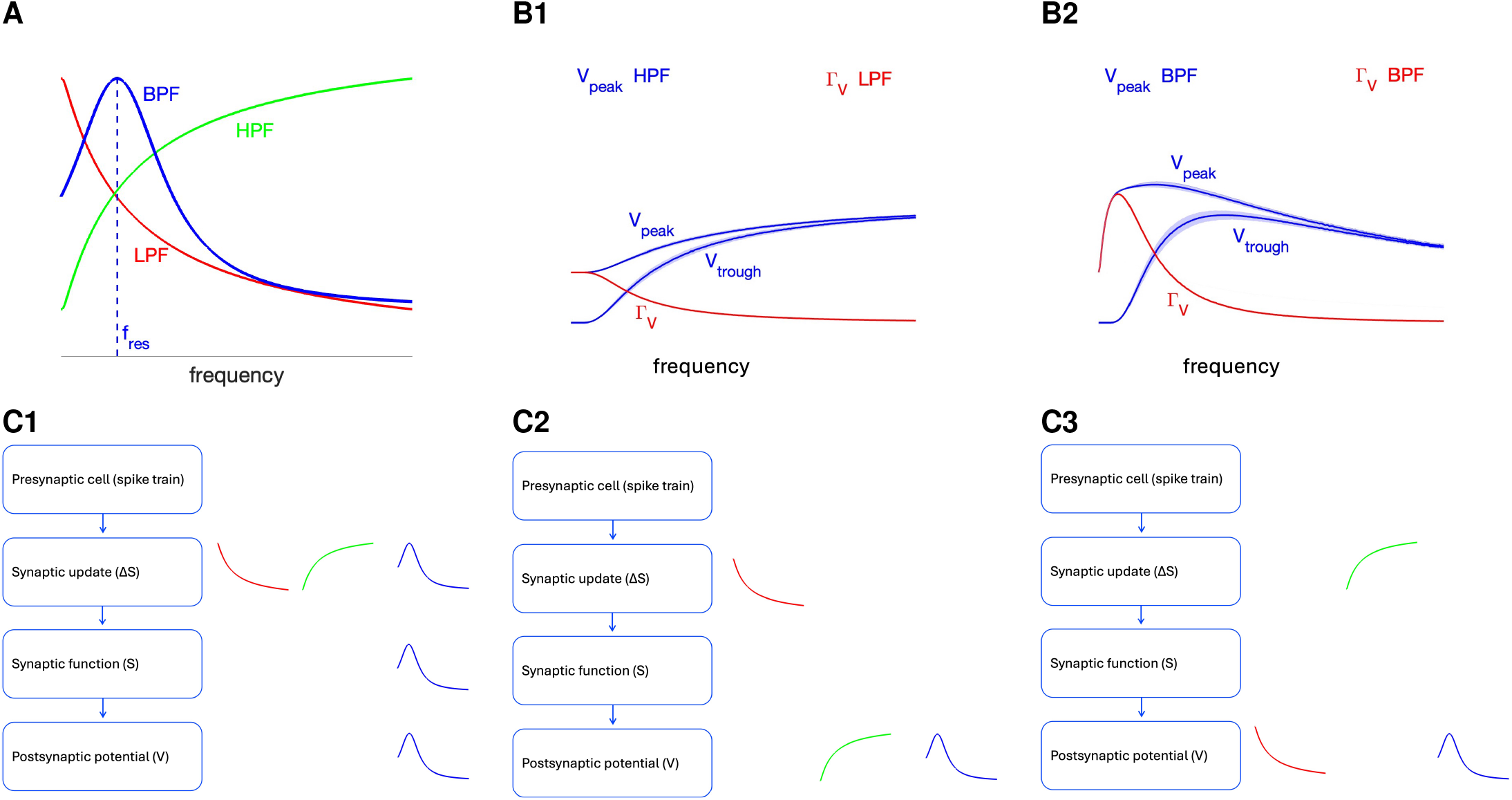
Postsynaptic potential (PSP) Peak (*V*_*peak*_) and amplitude (Γ_*V*_) filters can be generated within and across levels of organization and communicated across levels of organization. **A**. Band-pass filters (BPFs, blue) can be generated by the interplay of low-pass filters (LPFs, red) and high-pass filters (HPFs, green) within and across levels of neuronal organization and are shaped by the participating time constants (Fig. 1). The PSP resonant frequency *f*_*res*_ is the BPF’s peak frequency. **B**. The PSP (*V*_*peak*_) and PSP amplitude (Γ_*V*_ = *V*_*peak*_ − *V*_*trough*_) profiles (curves of these metrics as a function of the spiking input frequency) can have the different (HPF and LPF, left) or the same (both BPFs, right) filtering properties. **C**. Schematic diagrams of PSP BPFs (*V*_*peak*_ or Γ_*V*_) generated within and across levels of organization. **C1**. A PSP BPF can be generated by the interplay of a LPF and a HPF at the synaptic update level. The Δ*S* BPF is inherited to the PSP level. **C2**. A PSP BPF can generated by the interplay of a synaptic update LPF and a PSP HPF. **C3**. A PSP BPF can generated by the interplay of a synaptic update HPF and a PSP LPF. **C1, C2, C3**. The synaptic update BFPs, LPFs and HPFs can experience modulations as they are communicated across levels.

STP refers to the changes of the efficacy of synaptic transmission (synaptic conductance strength) in response to presynaptic spike trains (as the number spikes increases) with time scales ranging in the order of hundreds of milliseconds to seconds [25–27]. STP consists of the combination of two opposing processes with characteristic time scales: synaptic short-term depression (efficacy decrease; STD) and facilitation (efficacy increase; STF). STP has been investigated in both vertebrates and invertebrates. It has been shown to be involved in a number of brain functions, including information filtering (temporal and frequency-dependent) [3, 9–11, 17, 24, 26, 28–45], adaptive filtering [10] and related phenomena (e.g., burst detection) [3,35,46–49], temporal coding and information processing [35, 36, 50–53], information flow [42, 54, 55] (given the presynaptic history-dependent nature of STP), gain control [56–58], the modulation of network responses to external inputs [59, 60], the prolongation of neural responses to transient inputs [16, 61, 62], direction selectivity [63], vision (e.g., microsacades) [64], sound localization and hearing [65, 66], the generation of cortical up and down states [67], attractor dynamics [57, 68], navigation (e.g., place field sensing) [10, 39], working memory [62, 69], decision making [70] and neuronal computation [7, 55, 58, 71–73].

Neuronal resonance has been investigated both experimentally and theoretically at various levels of organization ranging from the cellular to the synaptic to the network levels [3, 4, 12, 18–21, 23, 48, 74–84]. In recent work [4], we demonstrated that BPFs can be inherited from lower (e.g., cellular) to higher (e.g., network) levels of organization or can be created independently at various levels of organization by the interplay of low-pass filters (LPFs) and high-pass filters (HPFs) belonging to the same or different levels (e.g., Fig. 2). However, much remains to be understood about the biophysical and dynamic mechanisms of generation of BPFs beyond the single cell level. It is unclear how neuronal filters are shaped by the time scales of the participating building blocks at each level of organization (Fig. 1). It is also unclear how the neuronal filtering properties are communicated across levels of organization, how they are modulated by the time scales and other biophysical properties as they transition from one level to another, and how they interact within and across levels of organization.

In this paper we systematically analyze and develop these ideas in a feedforward network motif (Fig. 1-A), which is arguably the elementary neuronal network processing unit and involves a multiplicity of interacting time scales.

The building blocks consist of a presynaptic spike-train (characteristic period Δ_*spk*_), a postsynaptic passive cell (time constant *τ*), an excitatory (AMPA) chemical synapse (rise and decay time constants *τ*_*rse*_ and *τ*_*dec*_, respectively), and STP (depression and facilitation constants *τ*_*dep*_ and *τ*_*fac*_, respectively). We use mathematical modeling, numerical simulations and analytical calculations on simplified models to analyze the PSP response to presynaptic spike trains across three levels of organization: (i) the so-called synaptic update level, which is the target of the synaptic variable and is affected by STP, (ii) the synaptic variable level, and (iii) the PSP level.

We first use periodic spike-trains with frequencies *f*_*spk*_ (period Δ_*spk*_) within some range to characterize the different types of filters (BPFs, LPFs and HPFs; Fig. 2-A) and to understand the mechanisms by which they are generated. We then use Poisson-distributed spike trains with mean rates within the same range (as the periodic spike trains) to extend our investigation and findings to more realistic scenarios. To link between the purely deterministic and purely stochastic approaches, we use the so-called jitter-periodic inputs [46]. The use of randomly distributed spikes also allow us to explore how the variability of the filtering properties is controlled by STP and other biophysical properties and time scales of the participating building blocks.

We compute the stationary, frequency-dependent PSP response to presynaptic spike trains. The temporal (transient) responses and the associated temporal filters in the same feedforward network were thoroughly investigated in [46]. We primarily focus on two metrics (Fig. 2-B) that give rise to two types of profiles (curves of the corresponding metrics as a function of the spike-train input frequency or firing rate): (i) peak profiles (*V*_*peak*_), (ii) peak-to-trough amplitude profiles (Γ_*V*_). For PSP filters, Γ_*V*_ is analogous to the impedance amplitude profile for the computation of the standard subthreshold filters (in response to direct activation of sinusoidal inputs), while *V*_*peak*_ is relevant for the communication of the PSP filtering effects to the spiking regime and therefore to other building blocks in larger networks. To a lesser extent, we consider a third metrics, the phase profiles, which extend the notion of impedance phase to the PSP responses to presynaptic inputs.

The effects of STP are present at the synaptic update level (Fig. 1-B, box 2) and are communicated to the synaptic level where they interact with the synaptic time scales (Fig. 1-B, box 3). The PSP filters result from the interaction between these variables and time scales and the biophysical properties and time scales of the postsynaptic cell (Fig. 1-B, box 4). Band-pass filters (BPFs) result from a combination of low-pass filters (LPFs) and high-pass filters (HPFs) (Fig. 2-A) operating at the same (Fig. 2-C1) or different (Fig. 2-C2 and -C3) levels of organization. PSP BPFs can be inherited from the synaptic update level (STP-mediated BPFs; Fig. 2-C1) or they can be generated across levels of organization due to the interaction between (i) a synaptic LPF and the PSP summation-mediated HPF filter (PSP peaks; Fig. 2-C2), and (ii) a synaptic HPF and the PSP summation-mediated LPF (PSP amplitude; Fig. 2-C3).

In spite of its simplicity, the feedforward network motif we use is dynamically rich. First, passive cells produce subthreshold (membrane potential) LPFs in response to sinusoidal inputs currents (the voltage amplitude response as a function of the input frequency is monotonically decreasing), but they may produce subthreshold HPFs in response to presynaptic periodic spike-train inputs due to the effects of summation. Second, synaptic depression and facilitation produce LPFs and HPFs, respectively in response to presynaptic spike inputs [3]. Third, the cellular filtering properties are primarily the result of electric circuit effects, while STP filtering properties are primarily the result of history-dependent processes. Finally, filters may be generated and interact within and across levels of organization [4, 46] as the result of the interplay of the biophysical building blocks that give rise to them. The lessons learned from this study will serve to construct a framework to analyze the mechanisms of generation of neuronal filters in networks with more complex cells and architecture.

The outline of the paper is as follows. In Section 2 we present the models and tools we use in this paper. We first describe the biophysically-plausible model for the postsynaptic response of passive cells to presynaptic spikes in the presence of STP. Then, we develop several reduced models for the synaptic dynamics, PSP dynamics and the dynamics of the membrane potential of passive cells in response to presynaptic spikes. These simplified models are amenable to analytical calculations, which allow us to understand how the interplay of the participating STP, synaptic and membrane time constants shape the PSP filters.

In Section 3.1 we analyze the generation of the synaptic update filters, which feed into the synaptic filters (and are the target of the synaptic variable *S*), in the presence of STP and the dependence of their characteristic frequencies and attributes (e.g., resonance frequency) on the single event STP time constants *τ*_*dep*_ and *τ*_*fac*_. In Section 3.2 we analyze the inherited and cross-level mechanisms of generation of synaptic) STP-mediated BPFs (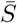 in a reduced model for synaptic dynamics. In Section 3.3 we analyze the inherited and cross-level mechanisms of generation of PSP BPFs in a reduced models of PSP dynamics in the presence of STP that incorporates the effects of summation (SUM) and SUM-mediated HPFs.

Armed with the insight and intuition gained from the previous sections, in Sections 3.4 and 3.5 we begin our analysis of the interplay of STP- and SUM-mediated filters in the postsynaptic cell, explain the mechanisms of generation of SUM-mediated HPFs in passive cells and compare the mechanisms by which they are generated with the mechanisms of generation of LPFs in response to direct sinusoidal inputs. In Section 3.6 we focus on the inherited and cross-level mechanisms of generation of PSP peak and amplitude BPFs (PSP resonance). These tools are used in the following section to understand the mechanisms underlying the generation of the different types of BPFs. In Section 3.7 we show that the interplay of STD-mediated LPFs and PSP SUM-mediated HPFs produce *V*_*peak*_ BPFs and Γ_*V*_ LPFs. In Section 3.8 we show that the Interplay of STF-mediated HPFs and PSP SUM-mediated filters produce *V*_*peak*_ LPFs and Γ_*V*_ BPFs. In Section 3.9 we show that interplay of STD-mediated LPFs, STF-mediated HPFs and PSP SUM-mediated filters produces *V*_*peak*_ and Γ_*V*_ BPFs. In all cases, the participating filters are modulated by the time constants not directly involved in their generation. The filters investigated in the previous sections are deterministic (in response to periodic presynaptic spike trains). In Sections 3.10 and 3.11 we extend our investigation to include randomly-distributed presynaptic spike trains. In Section 3.10 we show that the STP-mediated PSP *V*_*peak*_ and Γ_*V*_ BPFs persist with modulations in response to randomly-distributed spike trains both jittered-periodic and Poisson-distributed. In Section 3.11 we demonstrate that STP controls the variability of PSP *V*_*peak*_ and Γ_*V*_ BPFs in response to jittered-periodic and Poisson distributed spike trains in a frequency-dependent manner. We discuss our results and their implications in Section 4.

The development of the approximate solutions to simplified models is presented in the Appendix A. Tables of acronyms (Table 1), metrics (Table 2), model variables (Table 3, top) and model parameters (Table 3, bottom) are provided in the Appendix B.

**Table 1:** Acronyms.

|  |  |
| --- | --- |
| STP | Short-term plasticity |
| STD | Short-term depression |
| STF | Short-term facilitation |
| PSP | Post-synaptic potentials |
| LPF | Low-pass filter |
| HPF | High-pass filter |
| BPF | Band-pass filter |
| ISI | Interspike interval |
| SUM | Summation |
| DA | Dayan-Abbott (model) |
| MT | Markram-Tsodyks (model) |

**Table 2:**
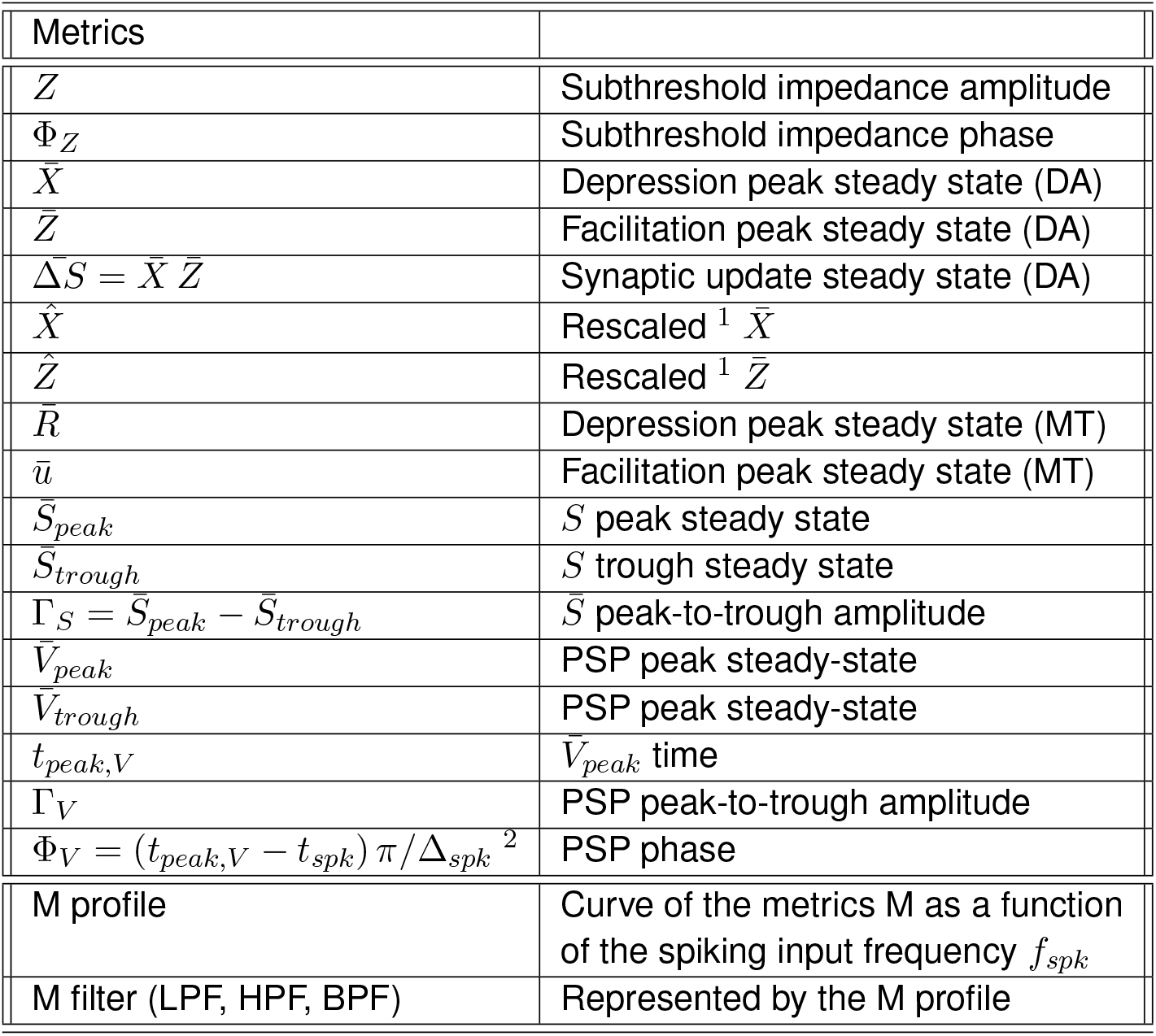
Metrics. DA refers to the DA model (Section 2.1.5). MT refers to the MT model (Section 2.1.7). ^1^ See eq. (44). ^2^ *t*_*spk*_ is the presynaptic spike time immediately preceding the occurrence of the *V* peak (*V*_*peak*_) at *t*_*peak,V*_

**Table 3:** Variables and parameters. DA refers to the DA model (Section 2.1.5). MT refers to the MT model (Section 2.1.7).

|  |  |  |  |
| --- | --- | --- | --- |
| $V$ | membrane potential | $t$ | time |
| $S$ | synaptic variable | $\Delta S$ | synaptic update (target value for $S$) |
| $x$ | depression variable (DA) | $z$ | facilitation variable (DA) |
| $R$ | depression variable (MT) | $u$ | facilitation variable (MT) |
| $\{X_n\}_{n=1}^{\infty}$ | depression peak sequence (DA) | $\{Z_n\}_{n=1}^{\infty}$ | facilitation peak sequence (DA) |
| $\{\Delta S_n\}_{n=1}^{\infty}$ | synaptic update sequence ($= \{X_n Z_n\}$) | | |
| $\{V_{peak,n}\}_{n=1}^{\infty}$ | PSP peak sequence | $\{V_{trough,n}\}_{n=1}^{\infty}$ | PSP trough sequence |
| $\{S_{peak,n}\}_{n=1}^{\infty}$ | $\bar{S}$ peak sequence | $\{S_{trough,n}\}_{n=1}^{\infty}$ | $\bar{S}$ trough sequence |
| $\{R_n\}_{n=1}^{\infty}$ | depression peak sequence (MT) | $\{u_n\}_{n=1}^{\infty}$ | facilitation peak sequence (MT) |
| $C$ | membrane capacitance | $I_{app}$ | applied current |
| $G_L$ | maximal leak conductance | $E_L$ | leak reversal potential |
| $G_{syn}$ | maximal synaptic conductance ($G_{ex}$) | $E_{syn}$ | synaptic reversal potential ($E_{ex}$) |
| $\tau_{dep}$ | synaptic depression time constant | $\tau_{fac}$ | synaptic facilitation time constant |
| $\tau_{rse}$ | synaptic rise time | $\tau_{dec}$ | synaptic decay time |
| $\tau = C/G_L$ | Membrane potential time constant | $\gamma_{syn}$ | $= G_{syn}/C$ |
| $\{t_{spk,n}\}_{n=1}^{\infty}$ | spike times sequence ($\{t_n\}_{n=1}^{\infty}$), | $\Delta_{spk,n}$ | presynaptic ISI sequence |
| $r_{spk}$ | presynaptic mean rates | $T_{sw}$ | presynaptic spike duration |
| $f_{spk}$ | presynaptic spiking frequency | $\Delta_{spk}$ | $= 1000/f_{spk}$ presynaptic ISI |
| $a_d$ | depression update (DA) | $a_f$ | facilitation update (DA) |
| $x_{\infty} (= 1)$ | depression saturation (DA) | $z_{\infty} (= 0)$ | facilitation saturation (DA) |
| $\sigma_{dep}$ | $\bar{X}$-filter characteristic time | $\sigma_{fac}$ | $\bar{Z}$-filter characteristic time |
| $\Delta S_{max}$ | $\Delta S$-BPF peak | $f_{\Delta S, res}$ | $\Delta S$-BPF resonant frequency |
| $\kappa_{rse}$ | $\Delta S$-BPF rise frequency | $\kappa_{dec}$ | $\Delta S$-BPF decay frequency |
| $U (= 1)$ | facilitation update (MT) | $\hat{U} (= 0)$ | facilitation saturation (MT) |
| $\alpha$ | see eq. (26) [22] | $\alpha_n$ | see eq. (26) [22] |
| $\sigma_s$ | see eq. (27) [23] | $\delta_{spk,n}$ | jitter perturbation sequence |
| $S_a$ | synaptic variable in eq. (27) [23] | $\eta_{stp}$ | see eq. (43) [38] |
| $f$ | sinusoidal input frequency | $\omega$ | $= 2\pi f/1000$ |
| $A_{in}$ | sinusoidal input amplitude | $\hat{\Delta}_{spk}$ | see eq. (43) [38] |
| $V_{eq}$ | $= E_L + I_{app}/G_L$ in eq. (26) | | |

## 2 Methods

### 2.1 Models

#### 2.1.1 Postsynaptic cell

The current-balance equation for the post-synaptic cell is given by

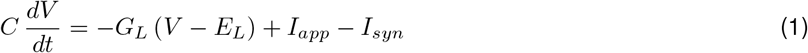

where *t* is time, *V* represents the voltage, *C* is the specific capacitance, *G*_*L*_ is the leak conductance, *I*_*app*_ is the tonic (DC) current, and *I*_*syn*_ is an excitatory synaptic current of the form

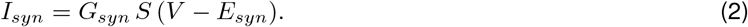

In eq. (2), *G*_*syn*_ is the maximal synaptic conductance, *E*_*syn*_ is the reversal potential and *S* is the synaptic variable.

We used the following units: ms for time, mV for membrane potential (voltage), *µ*F/cm^2^ for capacitance, mS/cm^2^ for conductance, *µ*A/cm^2^ for current and Hz for frequency.

#### 2.1.2 Synaptic dynamics

The synaptic variables *S* obey a kinetic equation of the form

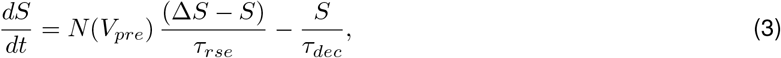

where *V*_*pre*_ is the membrane potential of the presynaptic spike, *N* (*V*) denotes the sigmoid function

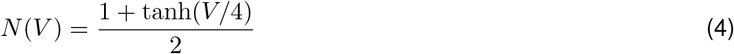

*τ*_*rse*_ and *τ*_*dec*_ are the rise and decay time constants respectively, and Δ*S* is a target value for *S*. For AMPA excitation (E-cells, *G*_*syn*_ = *G*_*ex*_), we used *E*_*syn*_ = *E*_*ex*_ = 0, *τ*_*rse*_ = 0.1 and *τ*_*dec*_ = 3.0 [85]. In the absence of synaptic short-term dynamics (depression and facilitation), Δ*S* = 1. Otherwise, Δ*S*, interpreted as the magnitude Δ*S* of the synaptic release per presynaptic spike, is determined as described below (Sections 2.1.4 and 2.1.6). We refer the reader to [86–88] for additional details on biophysical (conductance-based) models.

#### 2.1.3 Presynaptic spike-trains

We model the spiking activity of the presynaptic cell as a spike train with presynaptic spike times *t*_1_, *t*_2_, …, *t*_*N*_. We consider three types of input spike-trains. Periodic inputs are characterized by the interspike interval (ISI) of length Δ_*spk*_ or, alternatively, by the spiking frequency (Hz)

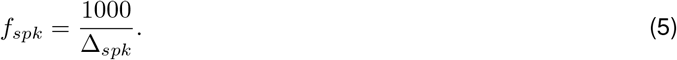

Jittered-periodic presynaptic spike trains consist of perturbations of periodic presynaptic spiking patterns of the form

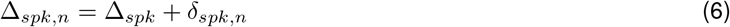

where Δ_*spk*_ is constant (*n*-independent) and 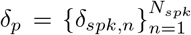 is a sequence of real numbers. We take *δ*_*p*_ to be normally distributed with zero mean and variance *σ*^2^. Poisson distributed (homogeneous) presynaptic spike trains are characterized by the mean spiking rate (and the associated exponential distribution of ISIs).

#### 2.1.4 The DA (Dayan-Abbott) model for short-term dynamics: synaptic depression and facilitation

This phenomenological model is presented in [87] and attributed to Dayan and Abbott (and collaborators). The magnitude Δ*S* of the synaptic release per presynaptic spike is assumed to be the product of the depression (*x*) and facilitation (*z*) variables

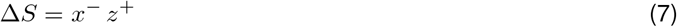

where

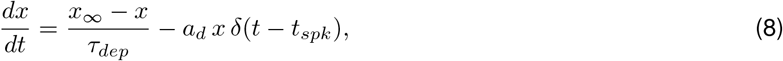

and

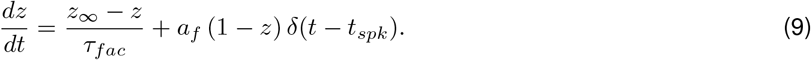

Each time a presynaptic spike arrives (*t* = *t*_*spk*_), the depressing variable *x* is decreased by an amount *a*_*d*_ *x* (the release probability is reduced) and the facilitating variable *z* is increased by an amount *a*_*f*_ (1 − *z*) (the release probability is augmented). During the presynaptic ISIs both *x* and *z* decay exponentially to their saturation values *x*_∞_ and *z*_∞_ respectively. The rate at which this occurs is controlled by the parameters *τ*_*dep*_ and *τ*_*fac*_. Following others we use *x*_∞_ = 1 and *z*_∞_ = 0. The superscripts “±” in the variables *x* and *z* indicate that the update is carried out by taking the values of these variables prior (^−^) or after (^+^) the arrival of the presynaptic spike.

Figs. 3-A1 and -B1 illustrates the *x*-, *z*-traces (curves of *x* and *z* as a function of time) in response to a periodic presynaptic input train for representative parameter values. The updated of the variable *S* at the arrival of presynaptic spikes is given by Δ*S* = *x*^−^*z*^+^. The sequence of peaks for Δ*S* is defined by Δ*S*_*n*_ = *X*_*n*_*Z*_*n*_ where *X*_*n*_ and *Z*_*n*_ are the sequence of peaks for the variables *x* and *z*, respectively (see also [46]).

**Figure 3.**
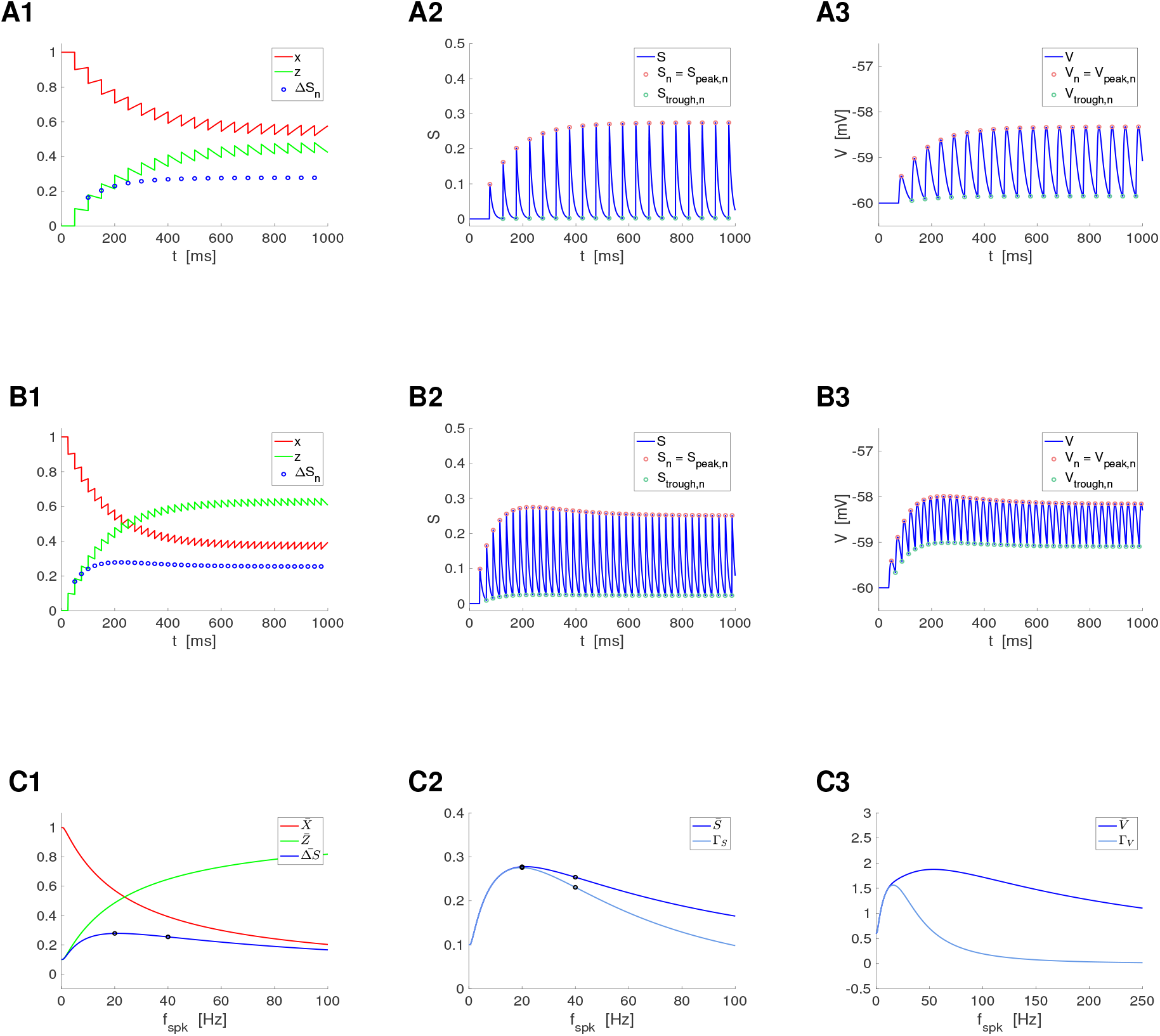
Representative temporal patterns for the synaptic update Δ*S*, the synaptic variable *S*, and the postsynaptic membrane potential *V* in the presence of short-term dynamics (STD). We used the model for the postsynaptic cell described by eqs. (1)-(4) with STD described by the DA model (7)-(9), and periodic presynaptic spike trains with frequency *f*_*spk*_ (see schematic Fig. 1, left). **Left column**. Short-term dynamics. The peak sequence Δ*S*_*n*_ (*n* = 1, …, *N*_*spk*_) is the synaptic update to the synaptic variable *S* upon the arrival of each presynaptic spike, and results from the combined effect of the depression (x) and facilitation (z) variables. The stationary value of the Δ*S*_*n*_ sequences is referred to as 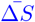. **Middle column**. Synaptic dynamics. The amplitude 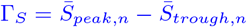, where 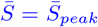 and 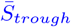 are the stationary values of the sequenes *S*_*peak,n*_ and *S*_*trough,n*_, respectively. **Right column**. Membrane potential dynamics. The amplitude 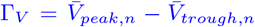, where 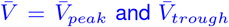 are the stationary values of the sequenes *V*_*peak,n*_ and *V*_*trough,n*_, respectively. **A**. *f*_*spk*_ = 20*Hz*. **B**. *f*_*spk*_ = 40*Hz*. **C**. Frequency profiles of the stationary peaks 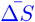 (left, blue), 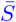 (middle, blue) and 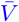 (right, blue) for the peak sequences Δ*S*_*n*_, *S*_*n*_ and *V*_*n*_, respectively, and stationary peak-to-trough amplitude profiles Γ_*S*_ (middle, light blue) and Γ_*V*_ (right, light blue) for *S* and *V*, respectively. The black dots correspond to the presynaptic input frequencies in *A* and *B*. We used the following additional parameter values: *a*_*d*_ = 0.1, *a*_*f*_ = 0.1, *x*_∞_ = 1, *z*_∞_ = 0, *τ*_*dep*_ = 400, *τ*_*fac*_ = 400, *τ*_*rse*_ = 0.1, *τ*_*dec*_ = 10, *C* = 1, *E*_*L*_ = −60, *G*_*L*_ = 0.1, *I*_*app*_ = 0, *G*_*syn*_ = 0.025, *E*_*syn*_ = 0.

#### 2.1.6 Response of the DA model to presynaptic inputs Peak dynamics and temporal filters

By solving the differential equations (8)-(9) during the presynaptic ISIs and appropriately updating the solutions at *t* = *t*_*n*_ (occurrence of each presynaptic spike), one arrives at the following recurrent formula for the peak sequences in terms of the model parameters

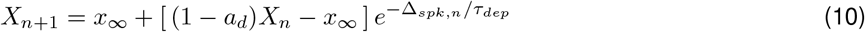

and

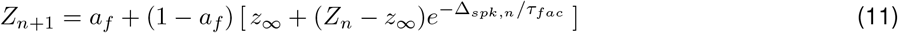

where 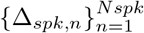 represents the lengths of the presynaptic ISIs. The temporal filtering properties of the DA model in response to periodic and Poisson-distributed presynaptic inputs was studied in [46].

##### Steady-state frequency-dependent filters

For periodic inputs, Δ_*spk,n*_ = Δ_*spk*_, independent of *n*, eqs. (10)-(11) are linear 1D difference equations. Therefore both the sequences *X* and *Z* obey linear discrete dynamics (e.g., see [89]), decaying to their steady state values

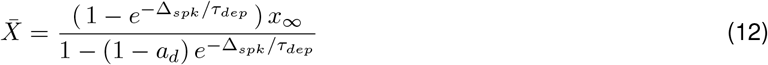

and

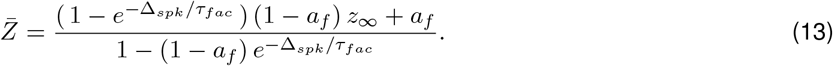

For the reminder of this paper we use *x*_∞_ = 1 and *z*_∞_ = 0.

#### 2.1.6 The MT (Markram-Tsodkys) model for short-term dynamics: synaptic depression and facilitation

This phenomenological model was introduced in [48] as a simplification of earlier models [25, 51, 90]. It is slightly more complex and widely used than the DA model described above [3, 91]. As for the DA model, the magnitude Δ*S* of the synaptic release per presynaptic spike is assumed to be the product of the depressing and facilitating variables

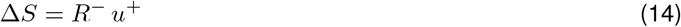

where, in its more general formulation,

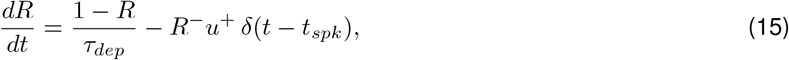

and

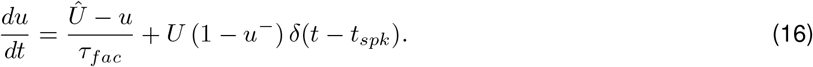

Each time a presynaptic spike arrives (*t* = *t*_*spk*_), the depressing variable *R* is decreased by *R*^−^*u*^+^ and the facilitating variable *u* is increased by *U* (1 −*u*^−^). As before, the superscripts “±” in the variables *R* and *u* indicate that the update is carried out by taking the values of these variables prior (^−^) or after (^+^) the arrival of the presynaptic spike. In contrast to the DA model, the update of the depression variable *R* is affected by the value of the facilitation variable *u*^+^. Simplified versions of this model include making *Û* = 0 [2, 30, 48, 49, 92, 93] and *Û* = *U* [3].

#### 2.1.7 Response of the MT model to presynaptic inputs Peak dynamics and temporal filters

By solving the differential equations (15)-(16) during the presynaptic ISIs and appropriately updating the solutions at *t* = *t*_*n*_ (occurrence of each presynaptic spike), one arrives at the following recurrent formula for the peak sequences in terms of the model parameters

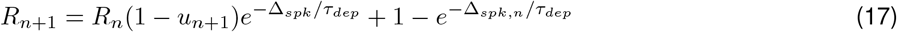

and

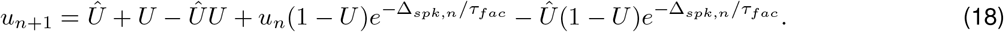

The temporal filtering properties of the DA model in response to periodic presynaptic inputs was studied in [46].

##### Steady-state frequency-dependent filters

As before, for periodic presynaptic inputs Δ_*spk,n*_ = Δ_*spk*_, independent of *n*, these equations represent a system of two 1D difference equations. The steady-state values are given by

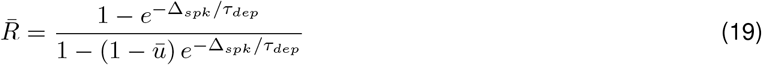

and

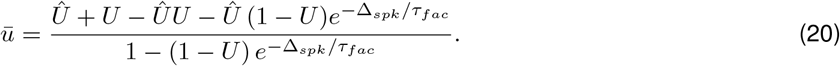

For the reminder of this paper we will use *Û* = 0 and *U* = 0.1.

### 2.2 Reduced models

#### 2.2.1 A reduced model for synaptic dynamics in response to presynaptic spikes: The to-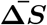 synaptic update model

Here and in the next section we present two sets of reduced models under certain simplifying assumptions. These simplified models are amenable to analytical calculations, which allow us to understand how the interplay of the participating STP time constants (*τ*_*dep*_ and *τ*_*fac*_) and synaptic time constants (*τ*_*dec*_ and *τ*_*rse*_) shape the synaptic and postsynaptic responses to presynaptic inputs.

For the parameter values consistent with AMPA excitation, the synaptic rise time *τ*_*rse*_ is very fast as compared to the synaptic decay time *τ*_*dec*_ and other times scales present in the model. Therefore, as a first level of approximation one can reduce the dynamics of the synaptic variable *S* in eq. (3) by

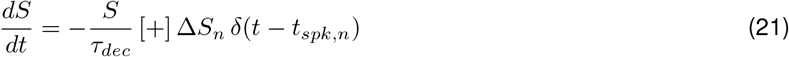

where the sign [+] indicates that each presynaptic spike (e.g., at time *t*_*n*_) instantaneously raises *S* “to” Δ*S*_*n*_, which depend on the combined dynamics of STD and STF. We refer to this model as the “to-Δ*S*” model in contrast to the scenario where each presynaptic spike instantaneously raises *S* “by” some value Δ*S*_*n*_, which is discussed in the next section.

For generality and to make the model more realistic, it is instructive to explore how the results obtained for the above simplified model are affected by the presence of non-zero synaptic rise times, while keeping the model simplified. To this end, we substitute the factor *N* (*V*_*pre*_) in eq. (3) by a square pulse for the duration of the spike and separate the rise and decay process. The extended to-Δ*S* model reads

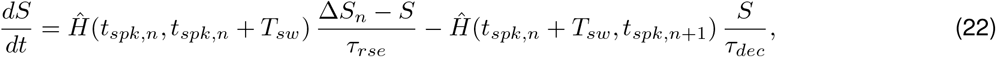

where *T*_*sw*_ is the spike width, *t*_*spk,n*+1_ = *t*_*spk,n*_ + Δ_*spk,n*_ and *Ĥ* (*t*_1_, *t*_2_) is a square pulse defined as the product of two Heaviside functions *H*(*t*), *Ĥ* (*t*_1_, *t*_2_) = *H*(*t* − *t*_1_) *H*(*t*_2_ −*t*). From the arrival of each spike and for the duration of this spike, *S* evolves according the first term in eq. (22), while for the remaining of the presynaptic period, *S* evolves according to the second term in eq. (22).

#### 2.2.2 A reduced model for PSP dynamics in response to presynaptic spikes: The by-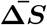 synaptic update model and the summation phenomenon

The reduced formulation we present here combines the synaptic and postsynaptic potential (PSP) responses to presynaptic input into a single equation. In contrast to the reduced models presented in Section 2.2.1, each presynaptic spike raises the variable *S* “by” some value Δ*S*_*n*_, which depends on the combined dynamics of STD and STF. This allow for the effect of summation to unfold.

In the models presented in here, the variable *S* is interpreted as the PSP response. Previous work by other authors have considered the PSP response frequency profiles to presynaptic inputs in the presence of STP to be proportional to the 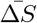 profiles (e.g., [30, 48, 93], but see [23]), without separating the dynamics of the synaptic and postsynaptic process. While we show that this is not always the case (as for STP-mediated temporal filters [46]), the use of the simplified models we present here are amenable to analytical calculations and allow to gain insight and intuition into the interplay of the STP and membrane time constants and the interaction between the STP-mediated 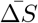 and summation filters in shaping the PSP filtering properties.

We first assume instantaneous rate. The “by-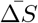” model reads

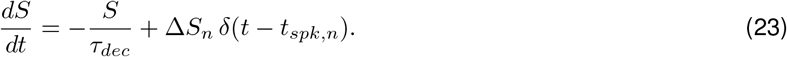

The “+” sign indicates that each presynaptic spike instantaneously raises *S* “by” Δ*S*_*n*_.

The extended by-Δ*S* model, which includes the dynamics of the rise process and rise constant *τ*_*rse*_ reads

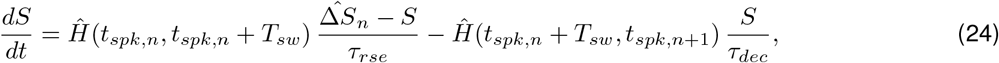

where 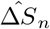 is the sum of Δ*S*_*n*_ and the value of *S* preceding the arrival of each presynaptic spike. The other components are as for the to-Δ*S* model (22).

#### 2.2.3 A reduced model for the membrane potential response of passive cells to presynaptic spikes

In order to analyze the PSP peak and amplitude responses of passive cells to presynaptic spikes we derive an analytical approximation of the solution to the model (1)-(4). The process consists of creating a reduced, hybrid model by substituting the conductance-based synaptic current in *I*_*syn*_ (2) by a presynaptic ISI-dependent current-based input where the synaptic input coefficient is updated every cycle to account for the changes in the driving force *V* −*E*_*syn*_ across cycles.

The postsynaptic passive cell (1) is linear, but the conductance-based synaptic input in *I*_*syn*_ (2) is multiplicative. Previous work showed that the subthreshold rhythmic properties of cells in response to conductance- and current-based synaptic inputs, where the driving force is substituted by a constant, may qualitatively differ [94, 95]. Therefore, a simply substitution of the product of the synaptic input and the driving force by the product of the synaptic input and a constant (a current-based synaptic input) will not produce good enough approximations to the responses of conductance-based synaptic inputs (e.g., Figs. S3 and S4). In the hybrid approach we develop here, we approximate *V* − *E*_*syn*_ by a constant, which we update at the end of each presynaptic ISI to account for the changing value of *V* − *E*_*syn*_ across cycles. This hybrid model is amenable to analytical calculations and therefore is an analytical tool to analyze how the participating time constants shape the PSP peak and amplitude filters.

We first approximate eq. (1) as follows

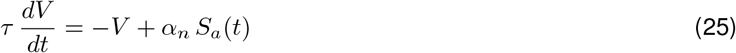

for *n* = 1, …, *N* where

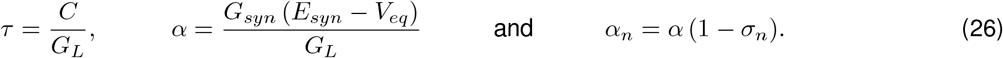

The variable *V* in eq. (25) represents *V* − *V*_*eq*_ in eq. (1) and *V*_*eq*_ = *E*_*L*_ + *I*_*app*_*/G*_*L*_. The right hand second term in eq. (25) is a current input approximation to the conductance input in the synaptic current eq. (1). To account for this, we introduce a correction factor 1 − *σ*_*n*_ where *σ*_*n*_ is updated at the beginning of each ISI proportionally to the distance between the PSP response and *E*_*syn*_ and *σ*_1_ = 0. The function *S*_*a*_(*t*) is an approximation to the variable *S* whose dynamics are described by eq. (3) with *S*(0) = 0, under the assumption of instantaneous rise to a value Δ*S*_*n*_ (*n* = 1, …, *N*) at the arrival of each presynaptic spike. The assumption *S*(0) = 0 implies that *S*(*t*) = 0 for 0 ≤ *t* ≤ *t*_1_.

For the duration of each presynaptic spike (*t*_*n*_ *< t < t*_*n*_ + *T*_*sw*_), we approximate *S*(*t*) by the synaptic update value Δ*S*_*n*_ (constant). For the remainder of the presynaptic interspike interval (*t*_*n*_ + *T*_*sw*_ ≤ *t < t*_*n*+1_), *S*(*t*) decreases according to the second term in (3). The approximate description of the synaptic term is given by

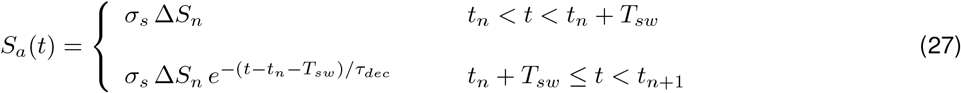

for *n* = 1, …, *N*. In our computations we take *T*_*sw*_ = 1 and *σ*_*s*_ = *τ*_*dec*_*/*(*τ*_*dec*_+*τ*_*rse*_) (1 − *exp*(−(*τ*_*dec*_+*τ*_*rse*_)*/*(*τ*_*dec*_*τ*_*rse*_))), which is the solution of eq. (3) computed at *t* = *T*_*sw*_ = 1. For *τ*_*rse*_ = 0.1 and *τ*_*dec*_ = 10, *σ*_*s*_ = 0.9901, while for *τ*_*rse*_ = 0.1 and *τ*_*dec*_ = 3, *σ*_*s*_ = 0.9677. The approximation error is *τ*_*dec*_*/*(*τ*_*dec*_ + *τ*_*rse*_) Δ*S τ*_*rse*_*τ*_*dec*_*/*(*τ*_*rse*_ + *τ*_*dec*_) − (1 + *τ*_*rse*_*τ*_*dec*_*/*(*τ*_*rse*_ + *τ*_*dec*_)) *exp*(−(*τ*_*dec*_ + *τ*_*rse*_)*/*(*τ*_*dec*_*τ*_*rse*_)). For *τ*_*dec*_ = 10, the error is equal to 0.0098 (0.098 Δ*S*) and for *τ*_*dec*_ = 3, the error is equal to 0.0094 (0.094 Δ*S*). However, we note that we consider the hybrid model to be a simplified model that captures the dynamics of the model (1)-(4) rather than a model that produces an accurate approximation to the solution of the model (1)-(4)

#### 2.2.4 Postsynaptic potential (PSP) peak sequences for the reduced model

We present here the main ideas and results. The detailed calculations are provided in the Appendix A.

For each input frequency *f*_*spk*_, we compute the PSP peak sequence *V*_*peak,n*_ (*n* = 1, 2, …) until |*V*_*peak,n*_ − *V*_*peak,n*−1_ |*< δ*_*tol*_ = 0.0001. This tolerance *δ*_*tol*_ is a conservative number that allows for the transient responses (temporal filters, see [46]) to wear off. We approximate the stationary value of 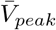 by *V*_*peak,n*_ (the last value in the resulting vector), *V*_*trough*_ by *V*_*trough,n*_, *t*_*V,peak*_ by the time at which *V*_*peak,n*_ occurs, and we use the corresponding value of *t*_*spk*_, *t*_*spk,n*_ (immediately preceding *V*_*peak,n*_) for the computation of the PSP phase response in eq. (42).

The PSP sequences are given by

##### PSP peak sequences for *τ*_*dec*_ */*= *τ*

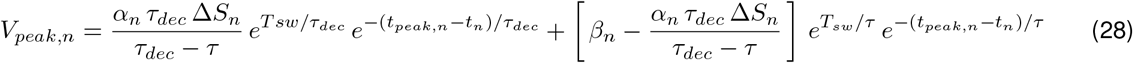

where

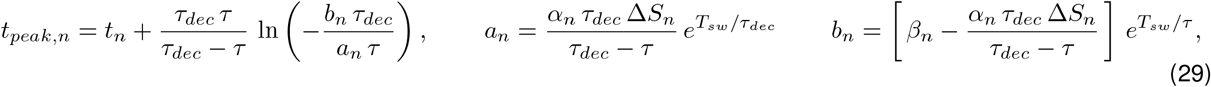

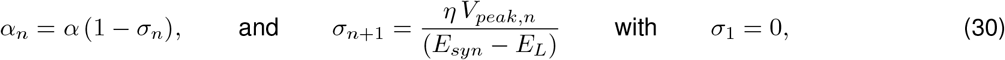

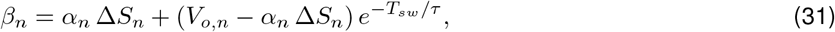

and

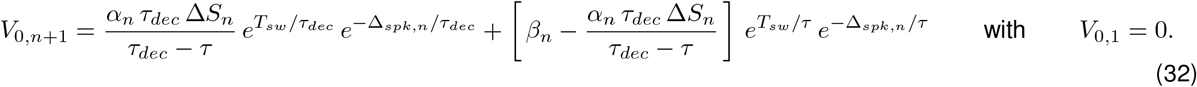

Eq. (28) is obtained from eq. (81) in the Appendix A. Note that *V*_*trough,n*_ = *V*_0,*n*+1_.

##### PSP peak sequences for *τ*_*dec*_ = *τ*

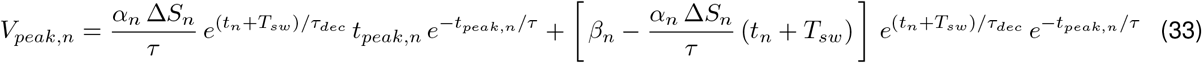

where

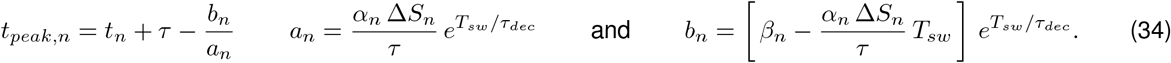

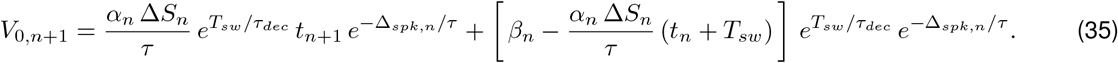

and *α*_*n*_, *σ*_*n*_ and *β*_*n*_ are given by eqs. (30) and (31). Eq. (88) is obtained from eq. (81) in the Appendix A. In both cases we use *η* = 1. Note that *V*_*trough,n*_ = *V*_0,*n*+1_.

Figs. S1 and S2 compare the numerical solutions to the model (1)-(4) and the analytical approximation using eqs. (25)-(27) together with eqs. (78) and (81) in the Appendix A for representative parameter values. The analytical approximation tracks the numerical solution with relatively high accuracy. Figs. S3 and S4 show the error between the numerical and analytical approximations to the stationary peaks (*V*_*peak*_), troughs (*V*_*trough*_) and peak times (*t*_*peak*_) of the membrane potential responses of passive cells to presynaptic spikes for representative parameter values. For *V*_*peak*_ and *V*_*trough*_ (left and middle columns), the error is significantly higher for *η* = 0 (green curves) than for values of *η* = 1 (blue curves) or around this value (red and light blue curves). Setting *η* = 0 is equivalent to the current-based synaptic input approximation, while setting *η* = 1 (or around this value), corrects for the driving force, which varies as *V* varies. In contrast, the error for *t*_*peak*_ (right columns) is largely independent of *η*.

### 2.3 Membrane potential response to presynaptic inputs and to direct activation of sinusoidal inputs

The filtering properties of passive cells (and of neurons in general) depend on whether they are directly activated by sinusoidal inputs (*I*_*in*_) or they are indirectly activated by periodic presynaptic inputs (*I*_*syn*_) within the same frequency range. We distinguish between the cell’s membrane potential (peak-to-trough) amplitude and peak response profiles (curves of these quantities as a function of the input frequency *f*; Fig. 2-B). For cells with linear subthreshold dynamics receiving direct sinusoidal current activation, these two metrics coincide. (For certain types of cells with nonlinear subthreshold dynamics, they are good approximations of each other [79, 80, 96, 97].) However, for presynaptic activation of postsynaptic cells, the PSP (peak-to-trough) amplitude and peak profiles do not generally coincide [95] (Fig. 2-B).

#### 2.3.1 Impedance and amplitude profiles: response to direct activation of sinusoidal input currents

The impedance of a neuronal system receiving an input current *I*_*in*_(*t*) is defined as the ratio of the output (voltage) *V*_*out*_(*t*) and input (current) Fourier transforms

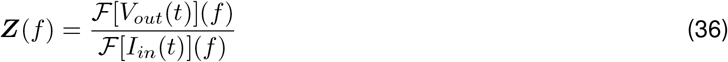

where *f* is frequency. The impedance ***Z***(*f*) is a complex quantity with amplitude *Z*(*f*) and phase Φ_*Z*_. For nonlinear systems, we use the Fast Fourier Transform algorithm (FFT) to compute these quantities.

#### 2.3.2 Impedance and phase profiles for the passive cell

The steady-state voltage response of a linear system receiving sinusoidal input currents of the form

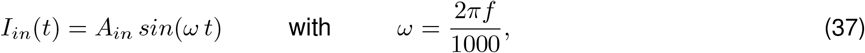

where *f* has units of Hz, is given by

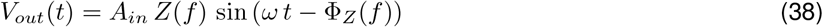

where *Z*(*f*) is the impedance amplitude and Φ_*Z*_(*f*) is the phase offset (time difference between the peaks of the input current and the output voltage normalized by 2 *π*).

For a passive cell of the form (1), standard calculations show that

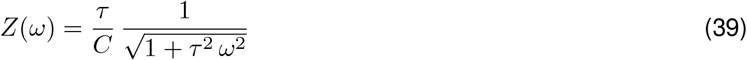

and

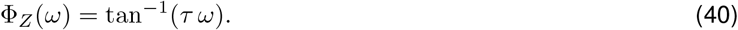

The passive cell is a low-pass filter since the *Z* (*f*) is monotonically decreasing. The response of the passive cell is delayed for all input frequencies and this delay increases with the input frequency since Φ_*Z*_ (*ω*) is monotonically increasing.

#### 2.3.3 PSP Peak, amplitude and phase profiles: response to presynaptic inputs

We characterize the PSP response to presynaptic inputs by using three metrics: the peak profiles, the peak-to-trough amplitude profiles and the phase profiles.

The PSP peak profiles *V*_*peak*_(*f*_*spk*_) are defined as the curves of the steady-state peak values of *V* as a function of the presynaptic spike input frequency *f*_*spk*_ (Fig. 2-B, blue). The PSP peak-to-trough amplitude profiles Γ_*V*_ (*f*_*spk*_) are defined as

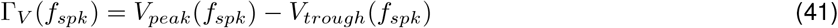

where *V*_*trough*_ are the PSP trough profiles (curves of the steady-state trough values of *V* as a function of *f*_*spk*_) (Fig. 2-B, red).

The PSP phase profiles Φ_*V*_ are defined as

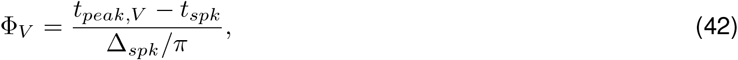

expressed in radians, where *t*_*peak,V*_ is the *V*_*peak*_ time and *t*_*spk*_ is the presynaptic spike time immediately preceding the occurrence of this peak (Fig. 9-A2, blue).

The Γ_*V*_ and Φ_*V*_ profiles are analogous metrics to the *Z* and Φ_*Z*_ profiles. The *V*_*peak*_ profiles capture the post-synaptic cell’s ability to preferentially produce spikes within certain presynaptic frequency ranges, and is therefore relevant for the frequency-dependent communication of information to the postsynaptic spiking regime.

### 2.4 Numerical simulations

The numerical solutions were computed using the modified Euler method (Runge-Kutta, order 2) [98] with a time step Δ*t* = 0.01 ms (or smaller values of Δ*t* when necessary) in MATLAB (The Mathworks, Natick, MA). The codes are available at \https://github.com/BioDatanamics-Lab/Frequency_Filters_STP_21_06.

## 3 Results

The question we ask in this paper is how the postsynaptic cell’s membrane potential frequency-filters (or -profiles) (curves of the appropriate metrics for each level of organization as a function of the input frequency *f*_*spk*_; e.g., Fig. 3-C), depend on the properties of the participating building blocks: the presynaptic spike trains, the synaptic rise and decay dynamics, synaptic short-term plasticity (STP) and the intrinsic properties of the postsynaptic cells (Fig. 1-A).

Specifically, we conduct a systematic study of the steady-state postsynaptic membrane potential (PSP) response to periodic presynaptic inputs over a range of frequencies *f*_*spk*_ = 1000*/*Δ_*spk*_ (Fig. 1-A, top) that capture the PSP filtering properties. We then extend our study to include jittered periodic inputs over a range of mean frequency *f*_*spk*_ = 1000*/*Δ_*spk*_ and Poisson-distributed presynaptic inputs over a range of mean rates *r*_*spk*_ = 1000*/ <* Δ_*spk*_ *>* (Fig. 1-A, bottom).

We divide our study in three steps (Fig. 1-B): (i) the response profiles of the synaptic update Δ*S* to the presynaptic spike trains, (ii) the response profiles of the synaptic variable *S* to Δ*S*, and (iii) the response profiles of the postsynaptic membrane potential *V* to *S*. Synaptic short-term plasticity (STP) operates at the Δ*S* level. The interaction between depression and facilitation (time constants *τ*_*dep*_ and *τ*_*fac*_, respectively) creates the synaptic update sequences Δ*S*_*n*_ = *X*_*n*_*Z*_*n*_ (Fig. 3, left column, blue dots) where *X*_*n*_ and *Z*_*n*_ are the sequence of peaks for the depression and facilitation variables *x* and *z*, respectively. These sequences are the target for the synaptic variables *S* during the rise phase after the arrival of each presynaptic spike. In the absence of STP, Δ*S*_*n*_ is constant (typically set up to one). The interplay of Δ*S*_*n*_ and the synaptic dynamics (rise and decay time constants *τ*_*rse*_ and *τ*_*dec*_, respectively) creates the response synaptic (*S*) patterns (Fig. 3, middle column). The synaptic variable *S* is the input to the current-balance equation (1) where the synaptic patterns interact with the postsynaptic biophysical membrane time constant *τ* to generate the postsynaptic (*V*) response patterns (Fig. 3, right column).

Here we focus on the frequency filtering properties of the steady-state responses for Δ*S*_*n*_, *S* and *V*. We characterize them by using the 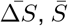 and 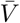 peak profiles (Fig. 3-C, blue), defined as the curves of the stationary peaks for the corresponding quantities as a function of the input frequency *f*_*spk*_, and the stationary peak-to-trough amplitude profiles (Fig. 3-C, light blue), consisting of the peak-to-trough amplitude curves as a function of *f*_*spk*_, for the latter two quantities.

The temporal filtering properties (transient responses to spike-spike trains) of these feedforward networks were systematically investigated in [46].

### 3.1 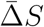 band-pass filters: interplay of low-pass (depression) and high-pass (facilitation) filters

From eqs. (12), (13) and (5), 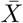 is monotonically decreasing (low-pass filter; LPF), transitioning from 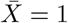 (*f*_*spk*_ = 0) to 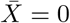 (*f*_*spk*_ → ∞), and 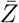 is monotonically increasing (high-pass filter; HPF), transitioning from 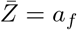 (*f*_*spk*_ = 0) to 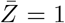 (*f*_*spk*_ → ∞). This is illustrated in Fig. 4 (red and green) for representative parameter values. The interplay of depression and facilitation produces 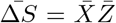 LPFs, BPFs or more complex patterns depending on the relative values of *τ*_*dep*_ and *τ*_*fac*_ (Fig. 4, blue), for fixed realistic values of the remaining parameters.

**Figure 4:**
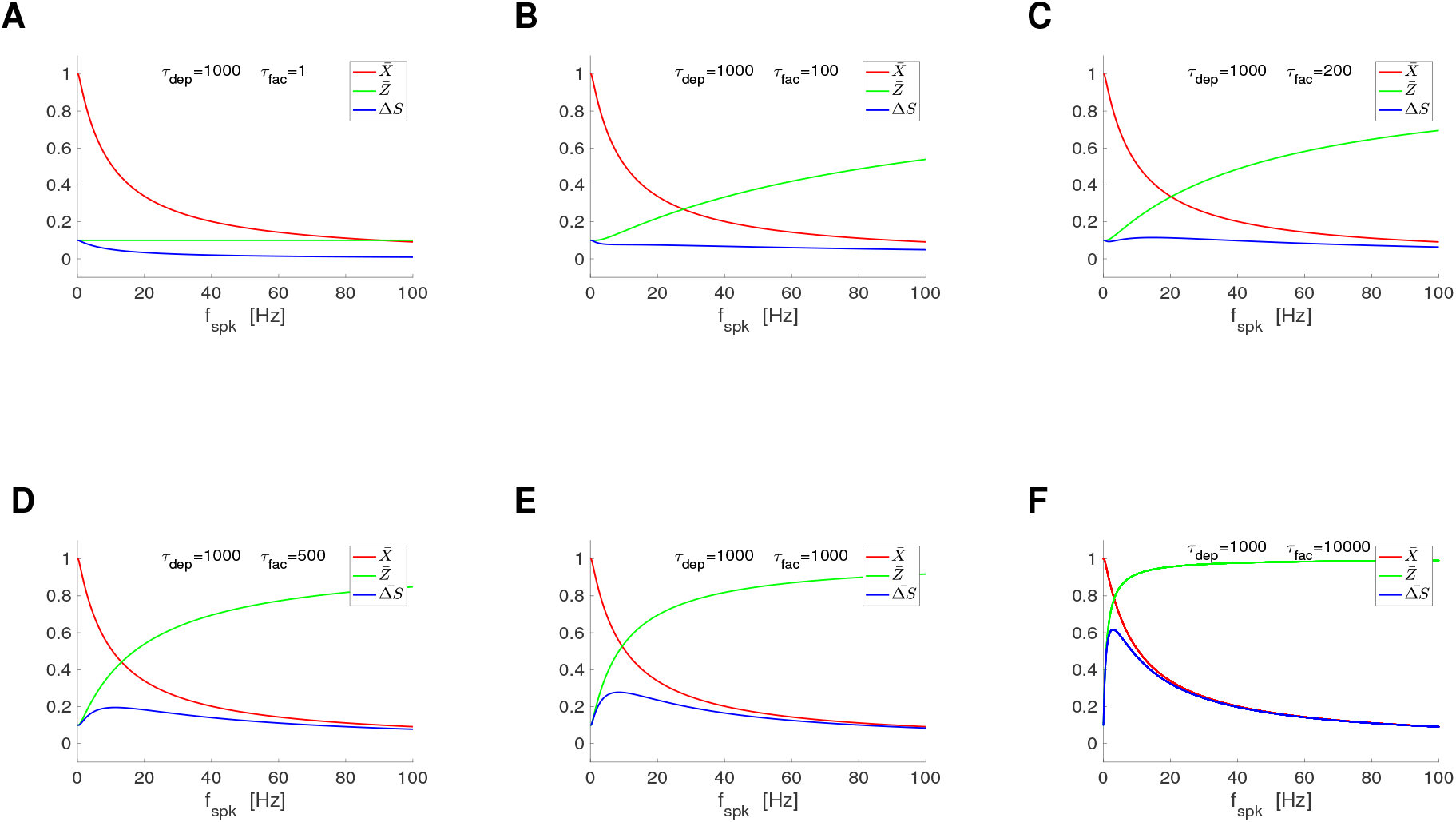
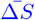 filters in response to periodic presynaptic spike inputs (frequency *f*_*spk*_) for the DA model: representative examples. We used eqs. (12) and (13). **A**. *τ*_*fac*_ = 1. **B**. *τ*_*fac*_ = 100. **C**. *τ*_*fac*_ = 200. **D**. *τ*_*fac*_ = 500. **E**. *τ*_*fac*_ = 1000. **F**. *τ*_*fac*_ = 10000. We used the following additional parameter values: *a*_*d*_ = 0.1, *a*_*f*_ = 0.1, *x*_∞_ = 1, *z*_∞_ = 0 and *τ*_*dep*_ = 1000.

#### 3.1.1 BPFs: A trade-off between depression- and facilitation-dominated regimes

To simplify the mechanistic analysis, we define

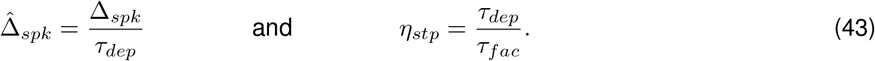

Substitution into eqs. (12) and (13) (with *x*_∞_ = 1 and *z*_∞_ = 0) yields

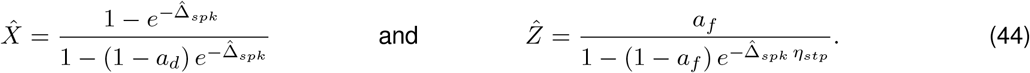

This rescaling allows us to investigate the mechanisms of generation of 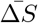 band-pass filters (BPFs) as a function of a single parameter (*η*_*stp*_) describing the relative magnitudes of the single event time constants *τ*_*dep*_ and *τ*_*fac*_. Specifically, 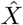 decreases with increasing values of 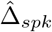 in a *η*_*stp*_-independent manner, while the rate of increase of 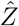 with 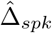 depends on the ratio *η*_*stp*_ of *τ*_*dep*_ and *τ*_*fac*_. The shapes of the 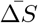 filters for all values of *τ*_*dep*_ and *τ*_*fac*_ unfold from the shapes of the corresponding 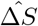 filters by reversing the rescaling.

For large enough values of *η*_*stp*_ (depression-dominated regime), the increase of 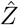 with increasing values of *f*_*spk*_ is much slower than the decrease of 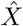 (in the limiting case *η* → ∞, 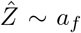, a constant). Therefore, 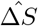 is a LPF (e.g., Fig. 4-A and -B, blue). For small enough values of *η*_*stp*_ (facilitation dominated regime), 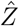 increases very fast with increasing values of *f*_*spk*_ as compared to 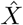 (in the limiting case *η*_*spk*_ → 0, 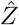 increases instantaneously and is approximately a constant, 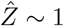). Therefore, 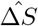 is also a LPF.

The transition between these two LPFs as *η*_*stp*_ changes occurs via the development BPFs (e.g., Fig. 4-D to -F, blue). Within some range of values of *η*_*stp*_ in between the LPFs and BPFs, the 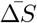 patterns develop a local minimum preceding the local maximum (e.g., Fig. 4-C, blue).

#### 3.1.2 From single-event time constants to frequency filters: Control of the filters’ shape by *τ*_*dep*_ and *τ*_*fac*_

The shapes of the 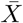 LPFs and 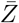 HPFs, and therefore the 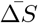 BPFs, are controlled by the time constants *τ*_*dep*_ and *τ*_*fac*_ operating at the “single-event” level, which govern the depression and facilitation dynamics, respectively, in response to each presynaptic spike. This is done by the communication of the single event time constants to the global time constants characterizing the temporal filters in response to repeated presynaptic inputs [46] in a presynaptic frequency-dependent manner. The 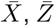 and 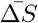 frequency filters are the steady-states of said temporal filters across presynaptic input frequencies.

We characterize the properties of the 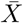 and 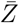 filters in terms of the characteristic frequencies *σ*_*dep*_ and *σ*_*fac*_, respectively (Fig. 5-A2). These attributes are defined as the frequencies for which the filters reached 63% of the gap between their values at *f*_*spk*_ = 0 and *f*_*spk*_ → ∞ (black dots in Figs. 5-A1 and -A2). For the characterization of the 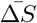 BPFs we use four attributes (Fig. 5-A3): the characteristic frequencies *κ*_*rse*_ and *κ*_*dec*_, the 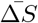 resonant frequency 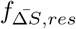 and the peak frequency 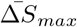. The characteristic frequencies were computed as the frequency difference between the peak and the frequency value at which 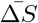 reached 63% of the gap between the peak and the value at *f*_*spk*_ = 0 (*κ*_*rse*_) and *f*_*spk*_ → ∞ (*κ*_*dec*_). The difference Δ*κ* = *κ*_*dec*_ − *κ*_*rse*_ is a measure of the spread of the BPFs.

**Figure 5.**
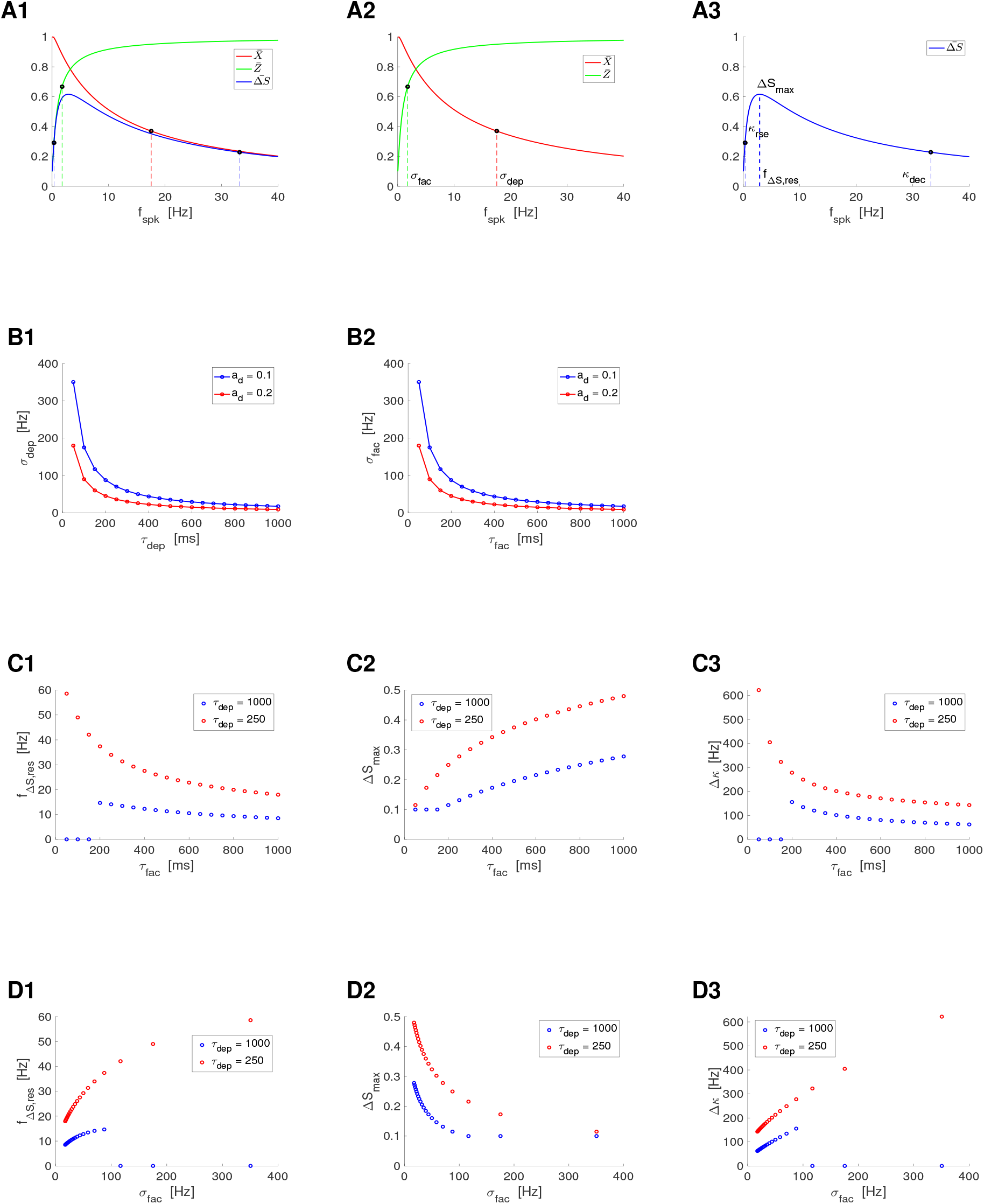
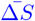 filters in response to periodic presynaptic spike inputs (frequency *f*_*spk*_) for the DA model: frequency attributes. **A**. The black dots on the 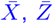 and 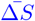 filters indicate the characteristic frequencies (projections on the *f*_*spk*_ axis) defined as the change in the corresponding quantities by 63 % of the gap between their final and initial values (*σ*_*dep*_ and *σ*_*fac*_) and between their maximum and minimum values (*κ*_*rse*_ and *κ*_*dec*_). The 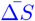 resonant frequency *f*_Δ*S,res*_ is the peak frequency and Δ*S*_*max*_ is the peak value. **B**. Dependence of the 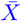 and 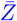 attributes (characteristic frequencies *σ*_*dep*_ and *σ*_*fac*_) with the depression and facilitation time constants *τ*_*dep*_ and *τ*_*fac*_, respectively. **C**. Dependence of the 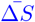 attributes with *τ*_*fac*_ for representative values of *τ*_*dep*_. **D**. Dependence of the 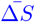 attributes with *σ*_*fac*_ for representative values of *τ*_*dep*_. For *τ*_*dep*_ = 1000, *σ*_*dep*_ ∼ 17.6, and for *τ*_*dep*_ = 250, *σ*_*dep*_1∼8 70.1.

Fig. 5-B shows the dependence of the characteristic frequencies for the 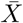 and 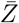 filters *τ*_*dep*_ and *τ*_*fac*_ (for representative values of *a*_*d*_ and *a*_*f*_). Specifically, *σ*_*dep*_ and *σ*_*fac*_ are decreasing functions of *τ*_*dep*_ and *τ*_*fac*_, respectively, and decreasing functions of *a*_*d*_ and *a*_*f*_, respectively. In other words, the larger the time constants, the more pronounced the decrease and increase of the corresponding filters with *f*_*spk*_.

Fig. 5-C shows the dependence of the attributes for the 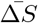 BPFs with *τ*_*dep*_ and *τ*_*fac*_. We fixed the value of *τ*_*dep*_ = 1000 (Fig. 5-C, blue) so the range of resonant frequencies 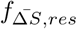 is relatively low. Using this information one can obtain the dependences for other values of *τ*_*dep*_ by reversing the rescaling (43). For comparison, we also present the results for *τ*_*dep*_ = 250 (Fig. 5-C, red). Specifically, the resonant frequency (*f*_Δ*S,res*_) decreases with increasing values of *τ*_*fac*_ and *τ*_*dep*_, the 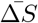 peak increases with increasing values of *τ*_*fac*_ and decreases with increasing values of *τ*_*dep*_ and the peak becomes sharper (Δ*κ* decreases) as *τ*_*fac*_ or *τ*_*dep*_ increase. Fig. 5-D shows the same results as a function of the characteristic frequency *σ*_*fac*_.

### 3.2 Interplay of 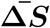 and 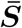 filters: inherited and cross-level mechanisms of generation of 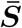 BPFs

We characterize the steady state response profiles of *S* to periodic presynaptic inputs by considering two attributes: the steady-state value 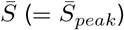 of the peak sequence *S*_*n*_ (= *S*_*peak,n*_) (Fig. 3, middle column, coral dots) and the peak-to-trough steady-state amplitude

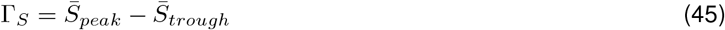

where 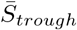 is the steady-state value of the trough sequence *S*_*trough,n*_ (Fig. 3, middle column, acquamarine dots). Fig. 3 (middle column) illustrates that both 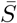 and Γ_*S*_ vary with the input frequency *f*_*spk*_. The temporal filtering properties of *S* using these two attributes were investigated in [46].

#### 3.2.1 The to-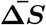 synaptic update model with instantaneous *S* rise

Here we use the approximate to- 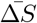 model (21) for the dynamics of the synaptic variable *S* under the assumption of instantaneous rise time. By construction, the steady state value 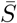 is given by

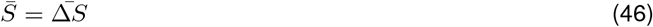

for all input frequencies *f*_*spk*_ (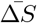 is the steady-state profile of the sequence Δ*S*_*n*_).

##### The 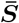 filtering properties are inherited from the synaptic update level

The peak envelope profiles 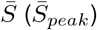 are identical to the 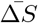 profiles for all input frequencies *f*_*spk*_. In the absence of STP, 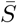 is a constant (= 1), while in the presence of STP, 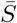 inherits the filtering properties from the synaptic update level discussed above.

In order to calculate Γ_*S*_, one needs to solve the differential equation (21) during the presynaptic ISIs (Δ*S*_*spk,n*_ = Δ*S*_*spk*_) and update the solution at the occurrence of each presynaptic spike at *t* = *t*_*spk,n*_ (*n* = 1, …, *N*_*spk*_), thus arriving to the following discrete linear difference equation for the trough sequences

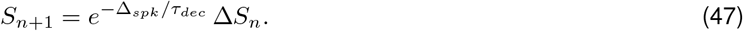

Therefore, the peak-to-trough envelope amplitude profile Γ_*S*_ is given by

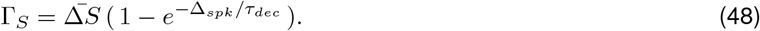

This expression is the product of two frequency-dependent processes: the 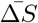 profile and

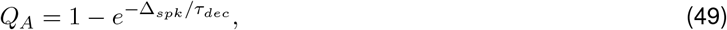

which is a LPF. The shape of the 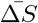 profile depends on the presence and properties of STP. In the absence of STP, the 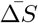 profile is independent of *f*_*spk*_ and Γ_*S*_ is a LPF.

As *f*_*spk*_ increases, the Γ_*S*_ profiles transition from Γ_*S*_ = 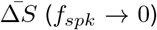 (*f*_*spk*_ → 0) to Γ_*S*_ = 0 (*f*_*spk*_ →∞) (Fig. 6). For fixed values of *f*_*spk*_, the Γ_*S*_ profiles transition from Γ_*S*_ = 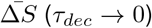 (*τ*_*dec*_ →0) to Γ_*S*_ = 0 (*τ*_*dec*_→ ∞) as *τ*_*dec*_ increases. In other words, for small enough values of *τ*_*dec*_, the Γ_*S*_ profiles reproduce the 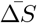 profiles (Fig. 6-A1 to A3), but for larger values of *τ*_*dec*_, the Γ_*S*_ and 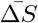 profiles are different. These differences increase as *f*_*spk*_ and *τ*_*dec*_ increase (Fig. 6). For generality, in Fig. 6 we included values of *τ*_*dec*_ beyond the biophysically plausible regime for AMPA excitation.

**Figure 6.**
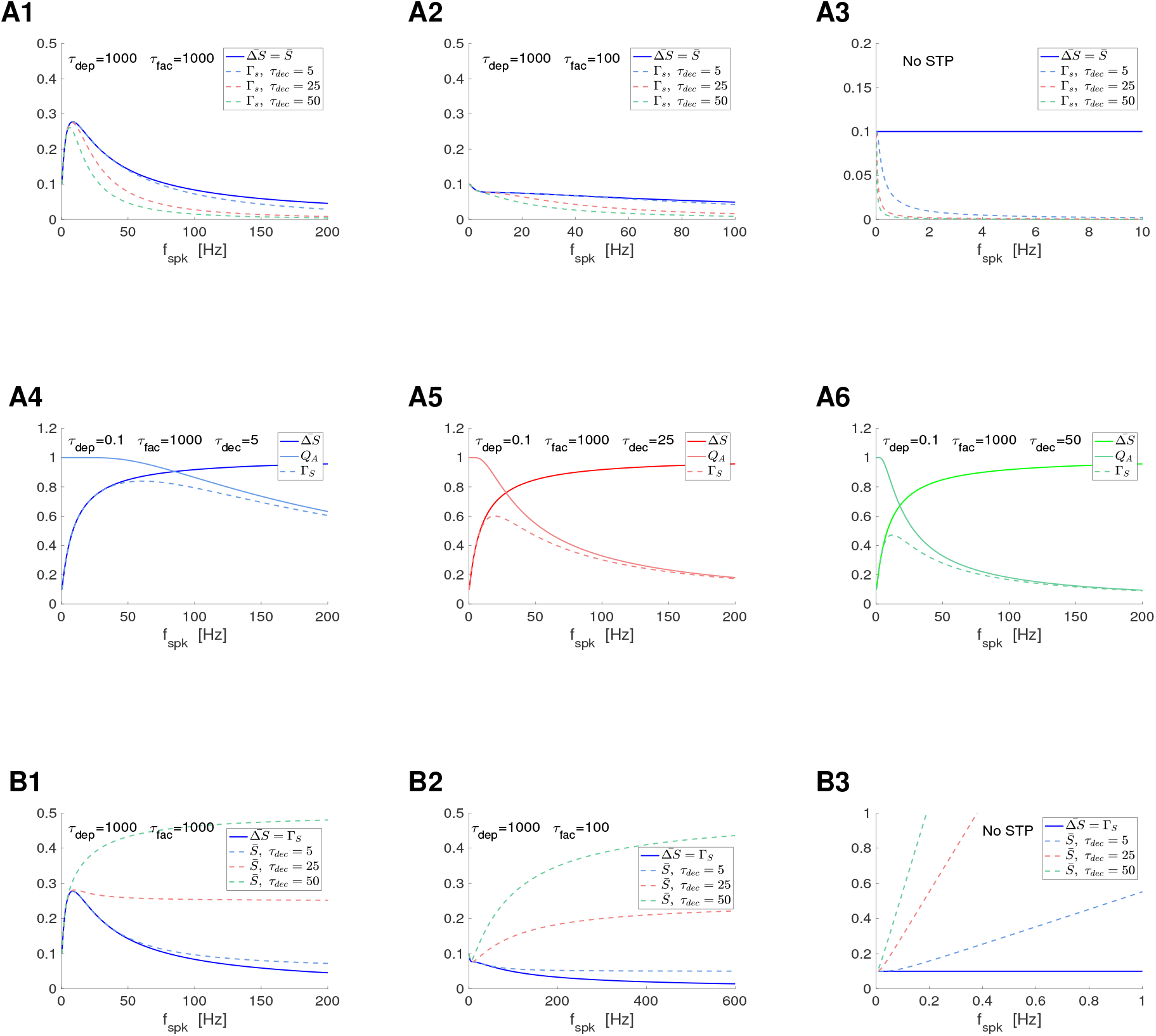
*S* and Γ_*S*_ filters in response to periodic presynaptic spike inputs (frequency *f*_*spk*_) for the to- and by-Δ*S* update models with instantaneous rise: representative examples. We used eqs. (12) and (13) (DA model) for 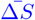. **A**. To-Δ*S* model (synaptic update to Δ*S*). We used eq. (48) for Γ_*S*_ and eq. (49) for *Q*_*A*_. **A1**. 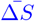 and Γ_*S*_ are band-pass filters. **A2**. 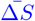 and Γ_*S*_ are low-pass filters. **A3**. No STP. 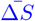 is constant and Γ_*S*_ are low-pass filters. **A4 to A6**. Γ_*S*_ band-pass filters generated from 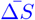 high-pass filters and *Q*_*A*_ low-pass filters. **B**. Synaptic update by Δ*S*. We used eq. (58) for 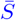. **B1**. 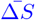 is a band-pass filter, while 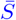 transitions form band-to high-pass filters as *τ*_*dec*_ increases. **B2**. 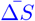 is primarily a low-pass filter, while 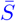 transitions form low-to high-pass filters as *τ*_*dec*_ increases. **B3**. No STP.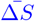 is constant, while 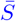 are high-pass filters. We used the following additional parameter values: *a*_*d*_ = 0.1, *a*_*f*_ = 0.1, *x*_∞_ = 1 and *z*_∞_ = 0.

##### Inherited mechanisms of generation of Γ_*S*_ BPFs

When 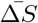 is constant (frequency-independent, no STP) or is an LPF, Γ_*S*_ is a LPF (Fig. 6-A2 and A3). These LPFs become more pronounced as *τ*_*dec*_ increases. When 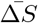 is a BPF, the Γ_*S*_ BPFs evoked by 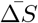 become sharper as *τ*_*dec*_ increases (Fig. 6-A1). These Γ_*S*_ filters are inherited from the 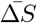 ones and modulated by *τ*_*dec*_.

##### Cross-level mechanisms of generation of Γ_*S*_ BPFs

In contrast, when 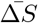 is a HPF, Γ_*S*_ BPFs emerge as the product of a HPF and a LPF (Fig. 6-A4). As *τ*_*dec*_ increases, the *Q*_*A*_ LPF is more pronounced as a function of *f*_*spk*_ and therefore the Γ_*S*_ BPF is sharper and peaks at a smaller value (compare Fig. 6-A4, -A5 and -A6).

#### 3.2.2 The to-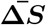 synaptic update model with non instantaneous *S* rise

Here we focus on the effects of the synaptic rise time on the generation and modulation of synaptic filters, particularly synaptic BPFs. We use the approximate model (22) for the dynamics of the synaptic variable *S* when the assumption of instantaneous rise is relaxed. The solution to the first and second terms in (22) are given by

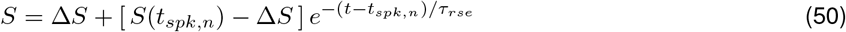

and

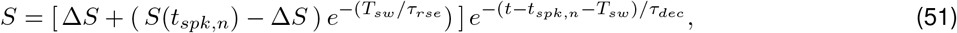

respectively. Using this, one can compute the difference equation governing the evolution of the sequence of peaks

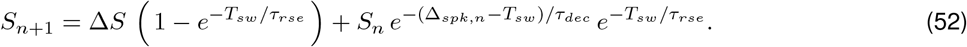

By assuming a constant Δ_*spk,n*_ = Δ_*spk*_, one obtains

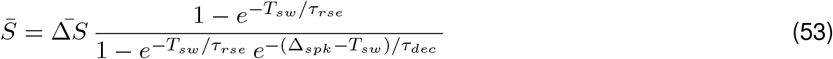

and

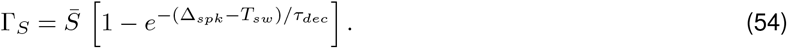

Both expressions are the product of frequency-dependent filters and reduce to eqs. (46) and (48) for *τ*_*rse*_ → 0 and *T*_*sw*_ → 0 with *τ*_*rse*_*/T*_*sw*_ *≪* 1.

We first focus on the 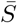 profiles. For *τ*_*rse*_ → 0, the second factor in eq. (53)

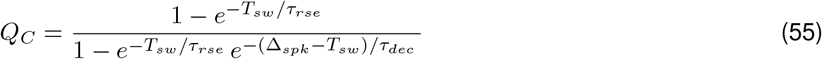

approaches *Q*_*C*_ = 1 for all *f*_*spk*_ and therefore the 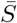 profiles are approximately equal to the 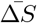 profiles. For *τ*_*rse*_ *>* 0, *Q*_*C*_ is a HPF and therefore the 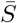 profiles depart from the 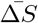 profiles Specifically, for *τ*_*rse*_ *>* 0, *Q*_*C*_ changes from 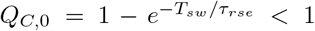 (for *f*_*spk*_ = 0) to 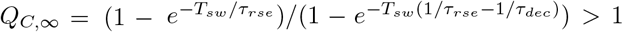 (as *f*_*spk*_→ ∞). *Q*_*C*_ = 1 for *f*_*spk*_ = 1000*/T*_*sw*_, independently of *τ*_*dec*_ and *τ*_*rse*_. As *τ*_*rse*_ increases (all other parameters fixed), within some bounds, *Q*_*C*,0_ decreases and *Q*_*C*,∞_ increases, causing an increase in the HPF amplitude of *Q*_*C*_. As *τ*_*dec*_ increases (all other parameters fixed), also within some bounds, *Q*_*C*_ increases for 0 *< f*_*spk*_ *<* 1000*/T*_*sw*_ and decreases for *f*_*spk*_ *>* 1000*/T*_*sw*_.

##### Attenuation of the [neq] filters (BPFs, LPFs and HPFs) inherited from the synaptic update level

Therefore, for *f*_*spk*_ *<* 1000*/T*_*sw*_, increasing values of *τ*_*rse*_ cause an attenuation of the 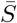 profiles (Fig. 7-A1 to -A3), and this attenuation is less pronounced the larger *τ*_*dec*_ (not shown). For *f*_*spk*_ *>* 1000*/T*_*sw*_, increasing values of *τ*_*rse*_ cause an amplification of the 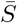 profiles (Fig. 7-A3), which is less pronounced the larger *τ*_*dec*_ (not shown). However, for *T*_*sw*_ = 1, the latter range is well beyond the frequencies we are interested in this paper.

**Figure 7.**
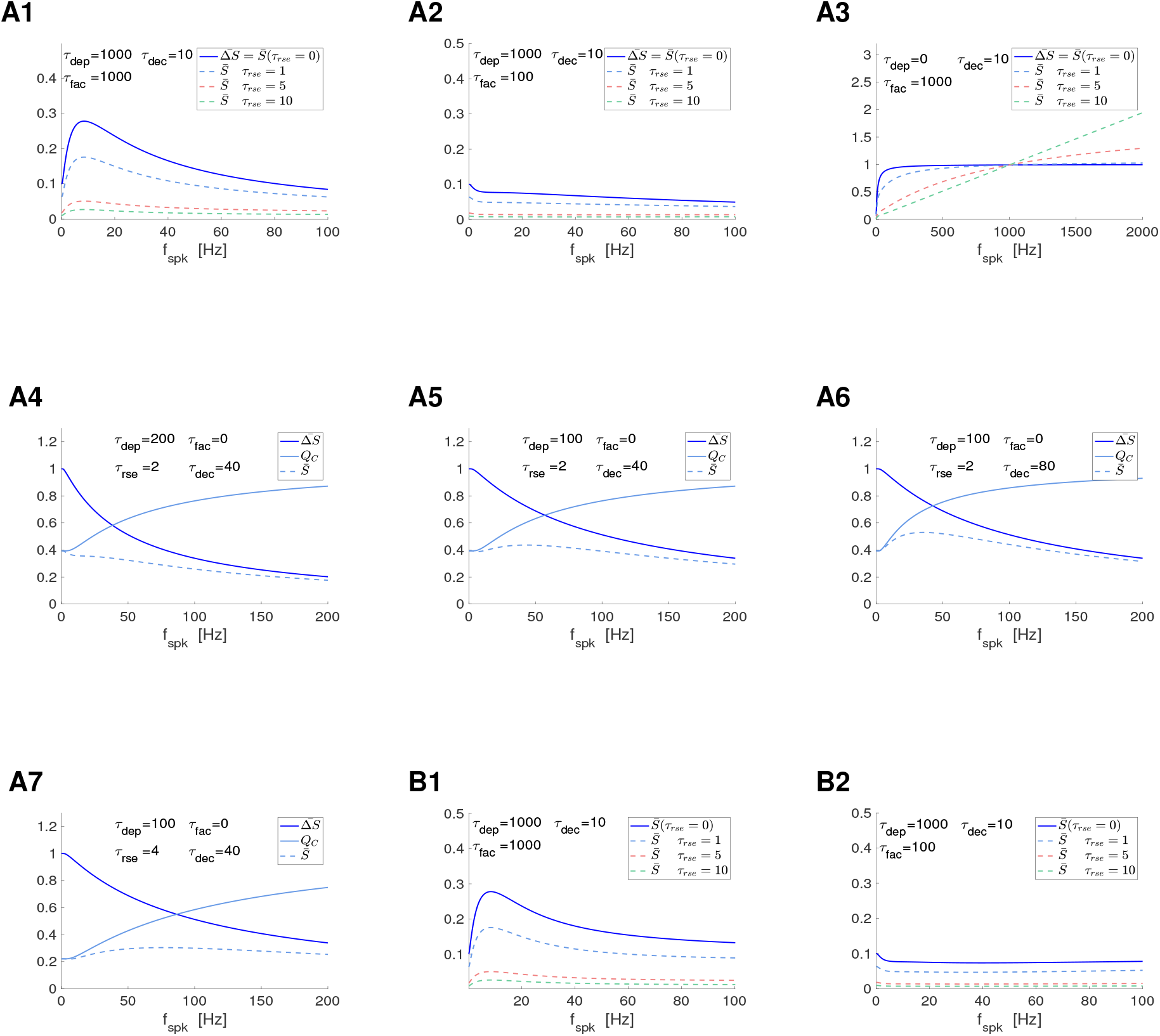
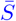 and Γ_*S*_ filters in response to periodic presynaptic spike inputs (frequency *f*_*spk*_) for the to- and by-Δ*S* update models with non-instantaneous rise: representative examples. We used eqs. (12) and (13) (DA model) for 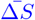 **A**. To-Δ*S* model (synaptic update to Δ*S*). The *S* and Γ_*S*_ filters were computed using eqs. (53) and (54), respectively, with *Q*_*C*_ and *Q*_*D*_ given by (55) and (56), respectively. **A1 to A3**. Effects of *τ*_*rse*_. **A1**. 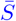 BPFs attenuated by increasing values of *τ*_*rse*_. **A2**. 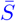 LPFs attenuated by increasing values of *τ*_*rse*_. **A3**. 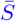 HPFs filters attenuated (amplified) by increasing values of *τ*_*rse*_ for lower (higher) values of *f*_*spk*_. **A4**. 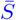 low-pass filters created by the interplay of a 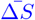 low-pass filter and a *Q*_*C*_ high-pass filter. **A5, A6, A7**. 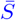 band-pass filters created by the interplay of 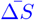 low-pass filters and *Q*_*C*_ high-pass filters. **B**. By-Δ*S* model (synaptic update by Δ*S*). The *S* and Γ_*S*_ filters were computed using eqs. (62) and (63), respectively, with *Q*_*E*_ given by (64). We used the following additional parameter values: *a*_*d*_ = 0.1, *a*_*f*_ = 0.1, *x*_∞_ = 1, *z*_∞_ = 0 and *T*_*sw*_ = 1.

The bounds mentioned above are set by the requirement that the denominator of *Q*_*C*_ is positive, which in turn requires that Δ_*spk*_ *>* −*T*_*sw*_(*τ*_*rse*_ *τ*_*dec*_)*/τ*_*rse*_. This is satisfied for all values of Δ_*spk*_ if *τ*_*rse*_ *< τ*_*dec*_. (For larger values of *τ*_*rse*_, this imposes a bound on Δ_*spk*_ for which *Q*_*C*_ *>* 0.) The realistic values of *τ*_*rse*_ and *τ*_*dec*_ we use here satisfy this condition. Moreover, for these values of *τ*_*rse*_ and *τ*_*dec*_, *Q*_*C*_ is a HPF, converging asymptotically to *Q*_*C*,∞_.

##### Cross-level mechanisms of generation of 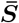 BPFs and attenuation of 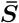 filters

Because *Q*_*C*_ has HPF properties, the question arises whether a 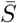 BPF can be created by the interplay of a 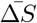 LPF and *Q*_*C*_ for nonzero values of *τ*_*rse*_. (For instantaneous rise times, 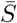 BPFs can only be inherited since 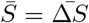; see Section 3.2.1). Figs. 7-A4 to -A7 illustrates that this is indeed possible *τ*_*rse*_ *>* 0. The generation of a 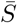 band-pass filter requires that the *Q*_*C*_(0) is low enough, which is achieved by increasing *τ*_*rse*_ above some threshold value (Fig. 7-A5). This BPF can be amplified by making the increase of *Q*_*C*_ sharper for low values of *f*_*spk*_, which can be achieved by increasing *τ*_*dec*_ (Fig. 7-A6). The BPF in Fig. 7-A5 is attenuated by further increasing *τ*_*rse*_ since this causes the intersection between the constituents low- and high-pass filters to move down (Fig. 7-A7). An increase in the values of *τ*_*dep*_ (Fig. 7-A7) causes the 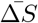 LPF to decrease sharper as compared to Fig. 7-A5, decreasing the intersection between the 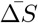 LPF and the *Q*_*C*_ HPF. The resulting attenuation produces a 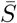 LPF (Fig. 7-A7).

##### Cross-level mechanisms of generation of Γ_*S*_ BFPs and attenuation of Γ_*S*_ filters

The second factor in eq. (54),

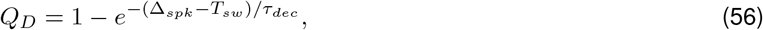

is a LPF, provided Δ_*spk*_ is large enough as compared to *T*_*sw*_, and is independent of *τ*_*rse*_. Therefore, the effect of *τ*_*rse*_ on the Γ_*S*_ filters is inherited from the effect of *τ*_*rse*_ on 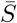 filters. The Γ_*S*_ filters are further attenuated by the *Q*_*D*_ filter. The Γ_*S*_ BPFs, in addition, become wider and the Γ_*S*_ resonant frequency is displaced (Fig. S14-A1, -A3). The attenuation is more pronounced for the larger frequencies as *τ*_*dec*_ increases, therefore the BPFs become sharper and the Γ_*S*_ resonant frequency is displaced as *τ*_*dec*_ increases (not shown).

### 3.3 Interplay of 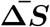 and summation PSP filters

The synaptic variable *S* determined by eq. (3) is the input to the current balance equation (1) for the postsynaptic voltage response *V*. In the absence of STP (Δ*S* = 1), summation effects give rise to postsynaptic (PSP) HPFs whose properties depend on the membrane potential properties, particularly the membrane time constant (*τ*). In the presence of STP, the PSP filters reflect the interaction of the 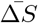 filters and the summation filters. For small enough values of *τ*, the PSP filters are well approximated by the 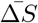 filters. Previous work by other authors has considered PSP and 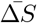 filters to be proportional (e.g., [30, 48, 93], but see [23]). However, for larger values of *τ*, the PSP filters are expected to depart from the weak modulation of the 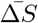 filters.

Here we us the “by-Δ*S*” model (23) described in Section 2.2.2 as an intermediate step for the investigation of the mechanisms that govern the interaction between the 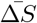 and PSP summation filters. We investigate the response of conductance-based models to periodic presynaptic inputs in the presence of STP in the next Section. The simplified model we study here has the advantage of being amenable to analytical calculations that provide a better insight into the mechanistic aspects of the interaction between the 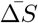 and PSP summation filters as compared to conductance-based models, and therefore pave the way for the investigation of said models.

#### 3.3.1 The by-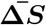 update model with instantaneous *S* rise

As a first step, we focus on instantaneous rise for *S*. In contrast to the to-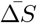 model, instantaneous rise is not a natural assumption when *S* is interpreted as the PSP because of the effects of the membrane time constant.

By solving the differential equation (23) for a constant value of Δ*S*_*spk,n*_ = Δ*S*_*spk*_ during the presynaptic ISIs and updating the solution at each occurrence of the presynaptic spikes at *t* = *t*_*n*_, *n* = 1, …, one arrives to the following discrete linear differential equation for the peak sequences in terms of the model parameters

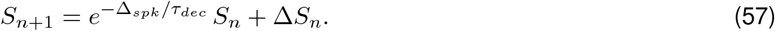

The 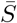 profile is given by the steady state values of (57)

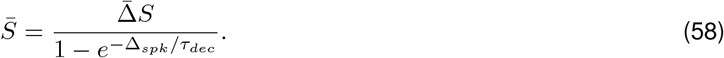

By construction, the Γ_*S*_ profiles are

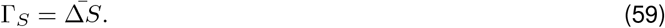

In other words, the Γ_*S*_ filtering properties are inherited from the 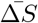 profiles.

##### The 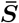 and 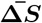 profiles depart from each other for relatively large values of *f*_*spk*_

Eq. (58) is the product of two frequency-dependent processes. The factor

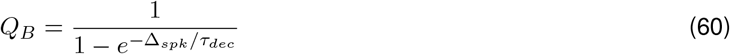

is a HPF. It transitions from *Q*_*B*_ = 1 (for *f*_*spk*_ = 0) to *Q*_*B*_ → ∞ (for *f*_*spk*_ → ∞), and it increases faster the larger *τ*_*dec*_. As already discussed, the shape of the 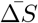 profile depends on the presence and properties of STP.

From eq. (58), for small enough values of *f*_*spk*_, 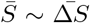. In the limit *f*_*spk*_ →0, 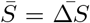. As *f*_*spk*_ increases, *Q*_*B*_ increases and therefore the difference between 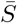 and 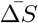 also increases.

##### 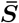 BPFs and LPFs transition to 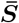 HPFs as *τ*_*dec*_ increases in the presence of STP

In the absence of STP, 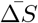 is constant and therefore 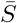 increases unboundedly as *f*_*spk*_ → ∞ (Fig. 6-B3). Similarly unbounded 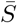 profiles are also obtained for 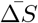 HPFs. Under certain circumstances, the presence of STP puts a bound on the increase of 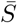, particularly for large values of *τ*_*dec*_ and the resulting filters remain bounded. Specifically, 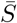 BPFs (Fig. 6-B1) and LPFs (Fig. 6-B2) may remain so for low enough values of *τ*_*dec*_ and transition to (bounded) HPFs for larger values of *τ*_*dec*_. However, these HPFs may reach saturation values that are too high to be realistic. This together with the presence of unbounded 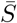 profiles (e.g., Fig. 6-B3) suggests that more complex biophysical models we investigate below include mechanism that cause the summation effects to be realistically saturated. Note that in some cases (e.g., Fig. 6-B2), the transition to a HPF involves the generation of a trough in the 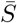 profiles for large enough values of *τ*_*dec*_.

#### 3.3.2 The by-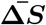 update model with non-instantaneous *S* rise

Here we extend our results to include the effects of noninstantaneous rise time. We use the approximate model (24) for the dynamics of the variable S when the assumption of instantaneous rise is relaxed.

The solution to the first and second terms in (24) are given by eqs. (50) and (51), respectively, with Δ*S* substituted by 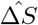. Using this, one can compute the difference equation governing the evolution of the sequence of peaks

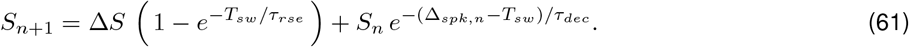

By assuming a constant Δ_*spk,n*_ = Δ_*spk*_, one obtains the 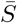 and Γ_*S*_ profiles

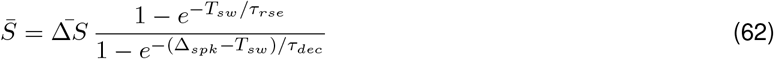

and

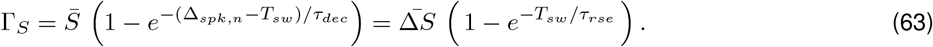

The 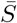 profile is the product of frequency-dependent processes: 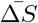 and

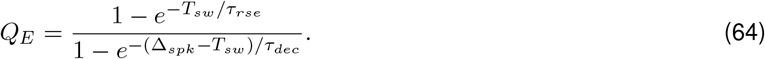

*Q*_*E*_ reduces to *Q*_*B*_ in eq. (49) for *τ*_*rse*_→ 0 and *T*_*sw*_ →0 with *τ*_*rse*_*/T*_*sw*_ *≪*1. This case was discussed in Section 3.3.1 and serves as a reference here.

##### 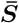 BFPs and LPFs are attenuated by increasing values of *τ*_*rse*_

The presence of *T*_*sw*_ in *Q*_*E*_ (for *τ*_*rse*_ *>* 0) causes a decrease in the initial values of *Q*_*E*_(*f*_*spk*_ = 0) and shrinks the range of values of *f*_*spk*_ for which the denominator of *Q*_*E*_ is positive to finite values: *f*_*spk*_ *<* 1000*/T*_*sw*_. As *f*_*spk*_ → 1000*/T*_*sw*_, *Q*_*E*_ increases unboundedly. This in turn causes 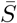 to increase unboundedly. However, this behavior is not alway monotonic. For low enough frequencies, but large enough to be within the range of realistic values we consider in this paper, the LPFs and BPFs (Fig. 7-B1 and -B2) inherited from the 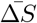 profiles (at the synaptic update level) and modulated by *τ*_*dec*_ are attenuated by increasing values of *τ*_*rse*_ (as discussed above). Away from this range of frequencies, the 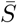 profiles for *τ*_*rse*_ *>* 0 increase above the 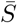 profile of *τ*_*rse*_ → 0 as they grow unboundedly. Increasing values of *τ*_*dec*_ amplify the grow of the 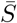 profiles consistent with the results of *τ*_*rse*_ → 0.

##### Γ_*S*_ profiles are uniformly attenuated by increasing values of *τ*_*rse*_

The second factor in eq. (63) is independent of *f*_*spk*_ and therefore increasing values of *τ*_*rse*_ attenuate the Γ_*S*_ filters without affecting their types (Fig. S15).

### 3.4 Digression: A heuristic explanation of summation and summation-mediated filters

Here and in the next sections we focus on the stationary membrane potential fluctuations of passive postsynaptic cells in response to periodic presynaptic inputs in the presence of STP (Fig. 1-A) for relatively fast synaptic rise and decay times (see Methods), consistent with AMPA excitation.

The passive postsynaptic cell is described by

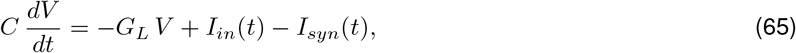

where the variable *V* in eq. (65) represents *V* − *V*_*eq*_ in eq. (1) with *V*_*eq*_ = *E*_*L*_ + *I*_*app*_*/G*_*L*_ and *I*_*syn*_ is described by eqs. (2)-(4) appropriately adapted (to account for the interpretation of *V* as membrane potential fluctuations around the equilibrium). In this section, we use the analytical approximations described in Section 2.2.4 (see also Appendix A).

#### 3.4.1 PSP response of passive cells to presynaptic spikes for individual input frequencies

Here we analyze the properties of the steady-state PSP response of passive cells to presynaptic spikes within one ISI, from the arrival of a presynaptic spike (*t* = *t*_*n*_) to the arrival of the next presynaptic spike (*t* = *t*_*n*+1_ = Δ_*spk,n*_). We then compare among the results for representative presynaptic ISIs (spiking input frequencies) to understand how the complex interaction among the participating time scales (*τ*_*dep*_, *τ*_*fac*_, *τ*_*dec*_, *τ*, Δ_*spk*_) and maximal synaptic conductance *G*_*syn*_ shape the PSP response profiles. Our results are presented in Fig. 8 for *f*_*spk*_ = 10 (left panels) and *f*_*spk*_ = 40 (right panels). We use the knowledge we gain from this analysis in the following sections.

**Figure 8.**
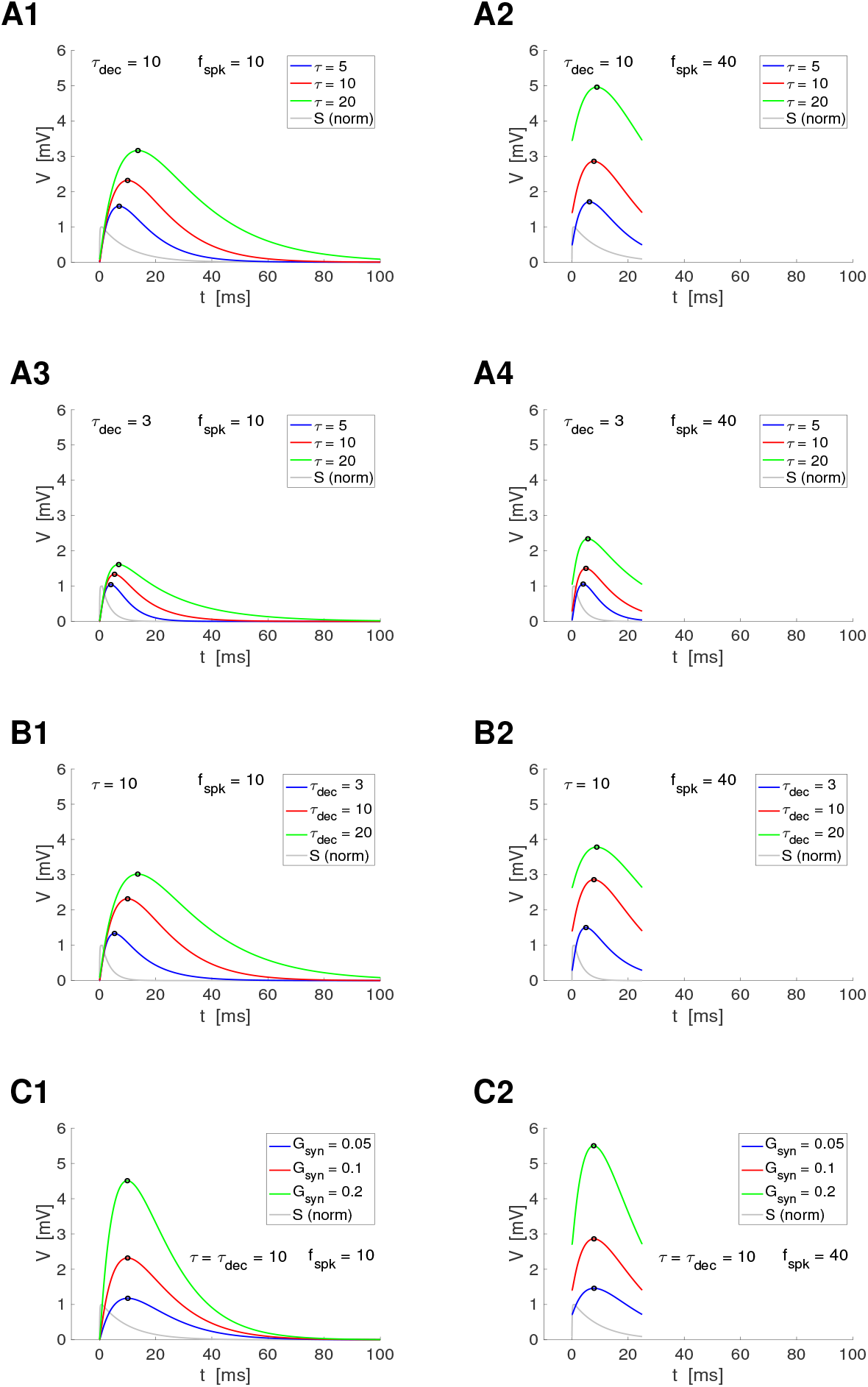
Properties of the membrane potential response of passive cells to periodic presynaptic spikes for representative parameter values. For the numerical approximations we used the model for a passive cell receiving presynaptic spike-train input (1)-(4). For STP we used the DA model (7)-(9). The graphs correspond to the steady-state solutions (translated to *t* = 0). The summation effect is observed in panels B. The synaptic function *S* was normalized by its maximum in the presynaptic interspike interval. **A**. *G*_*syn*_ = 0.1 **Top row:** *τ*_*dec*_ = 10. **Bottom row:** *τ*_*dec*_ = 3. **Left column**. *f*_*spk*_ = 10. **Right column**. *f*_*spk*_ = 40. *G*_*syn*_ = 0.1. **B**. *G*_*syn*_ = 0.1 and *τ* = 10 **B1**. *f*_*spk*_ = 10. **B2**. *f*_*spk*_ = 40. **C**. *τ* = 10, *τ*_*dec*_ = 10 **C1**. *f*_*spk*_ = 10. **C2**. *f*_*spk*_ = 40. We used the following additional parameter values: *a*_*d*_ = 0.1, *a*_*f*_ = 0.1, *x*_∞_ = 1, *z*_∞_ = 0, *τ*_*rse*_ = 0.1, *C* = 1, *E*_*L*_ = −60, *I*_*app*_ = 0, *E*_*syn*_ = −60, *τ*_*dep*_ = *τ*_*fac*_ = 0.1.

By construction (see Section 2.2.4 and Appendix A.2), the analytical approximation to *V* (*t*) consisting of *V*_*I*_ (*t*) (78) for the duration of the presynaptic spike (*t*_*n*_ *< t < t*_*n*_ + *T*_*sw*_) followed by *V*_*II*_ (*t*) (81 or 88) for the reminder of the presynaptic ISI (*t*_*n*_ + *T*_*sw*_ *< t < t*_*n*_ + Δ_*spk,n*_), depends on the model parameters both explicitly and implicitly through the initial condition *V*_0,*n*_ for each presynaptic ISI and the update parameter *α*_*n*_. The implicit dependence is inherited from the previous presynaptic ISI. We assume here the PSP response is in the steady-state regime and therefore we focus our analysis on the explicit dependence of *V* (*t*) on the model parameters.

From eq. (78) for *V*_*I*_ (*t*) (*t*_*n*_ *< t < t*_*n*_ + *T*_*sw*_), the larger *τ*, the larger *β*_*n*_ = *V*_*I*_ (*t*_*n*_ + *T*_*sw*_) = *V*_*I*_*I*(*t*_*n*_ + *T*_*sw*_) (79). This dependence is affected by the presynaptic ISI Δ_*spk,n*_ through *V*_0,*n*_ = *V*_*II*_ (*t*_*n*−1_) (from the previous presynaptic ISI). For the remainder of the presynaptic ISI (*t*_*n*_ + *T*_*sw*_ *< t < t*_*n*+1_), from eqs. (81) and (88), respectively,

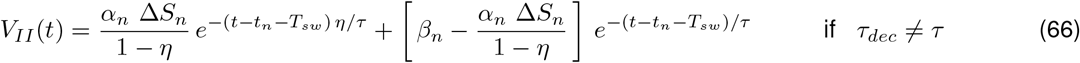

with

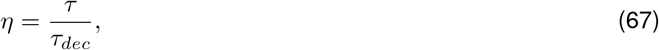

and

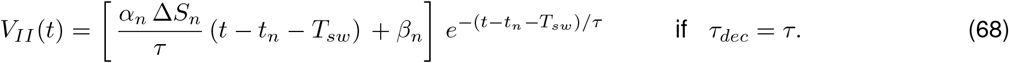

The PSP response *V* (*t*) to presynaptic inputs is shaped by a complex balance among *τ*_*dec*_, *τ* and Δ_*spk,n*_ (Fig. 8), and is modulated by *G*_*syn*_ (constant) and the STP time constants (*τ*_*dep*_ and *τ*_*fac*_) through Δ*S*_*n*_ (in a frequency-dependent manner), which determines the target for the peak of the synaptic function *S*. In Fig. 8, Δ*S*_*n*_ is independent of *f*_*spk*_. The properties of the PSP response for frequency-dependent Δ*S*_*n*_ profiles are investigated in the next Sections.

If *τ*_*dec*_ *≪τ, η≫* 1 in eq. (66) and *V*_*II*_ (*t*) is dominated by the second term. In the limiting cases *η* → ∞ (*τ*_*dec*_ →0 or *τ* → ∞), *V*_*II*_ (*t*) begins to decrease at *t*_*n*_ + *T*_*sw*_. As *η* decreases, *V*_*II*_ (*t*) continues to increase passed *t*_*n*_ + *T*_*sw*_ in response to *S*_*a*_(*t*) *>* 0. The larger *τ*_*dec*_ for fixed values of *τ*, the larger *V*_*II*_ (*t*) over the presynaptic ISI (Fig. 8-B) and the larger the peak time *t*_*peak,n*_. Similarly, the larger *τ* for fixed values of *τ*_*dec*_, the larger *V*_*II*_ (*t*) over the presynaptic

ISI (Fig. 8-A) and the larger the peak time *t*_*peak,n*_. If *τ*_*dec*_ *≫ τ, η ≪* 1 in eq. (66) and *V*_*II*_ (*t*) is dominated by the first term. In the limit *η* →0 (*τ*_*dec*_ → ∞ or *τ* → 0), *V*_*II*_ (*t*) only increases over the presynaptic ISI. If *τ*_*dec*_ = *τ*_*fac*_, *V*_*II*_ (*t*) has the form of an alpha function. The larger *τ*, the larger *V*_*II*_ (*t*) over the presynaptic ISI. Increasing values of *G*_*syn*_ also cause an increase in *V*_*II*_ (*t*) over the presynaptic ISI (Fig. 8-C) with at most a mild effect on the peak times. This suggests that while increasing values of *τ*_*dec*_ and *G*_*syn*_ increase the total input current to the postsynaptic cell, their effect on the properties of the PSP filters may differ.

Comparison between the left (*f*_*spk*_ = 10) and right (*f*_*spk*_ = 40) columns in Fig. 8 illustrates the effects of summation. One of them is the increase in the PSP peak response as *f*_*spk*_ increases and the other one is the decrease in the peak-to-trough amplitude as *f*_*spk*_ increases.

#### 3.4.2 PSP Summation-mediated filters

By approximating *S* by Δ*S*_*n*_ for the duration of the presynaptic spike (*t*_*n*_ ≤ *t* ≤ *t*_*n*_ + *T*_*sw*_), the passive membrane equation receiving presynaptic inputs equation is approximated by

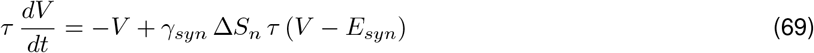

where the variable *V* has the same interpretation as in eq. (25), *V* (0) = 0 and

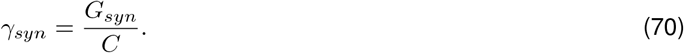

The solution to eq. (69) is given by

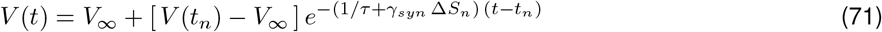

where

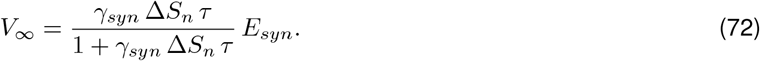

*V* (*t*) increases and approaches its steady-state value *V*_∞_, which increases from *V*_∞_ = 0 (*τ* = 0) to *V*_∞_ = *E*_*syn*_ (*τ* → ∞). *V* (*t*) reaches its peak value

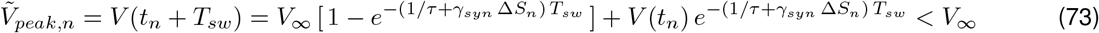

during the presynaptic ISI. For the remaining of the presynaptic ISI (*t*_*n*_ + *T*_*sw*_ *< t < t*_*n*+1_), *V* (*t*) decays exponentially to some value 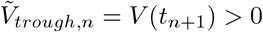, which depends on the presynaptic ISI Δ_*spk,n*_.

The peak value 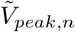 increases from 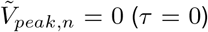 to 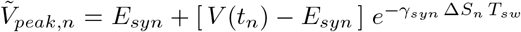 (*τ*→ ∞), and is an increasing function of *V* (*t*_*n*_) whose value is inherited from the previous presynaptic ISI.

For periodic presynaptic inputs and synaptic update values Δ*S*_*n*_ independent of *n*, the *V*_*peak*_ and Γ_*V*_ summation are originated in the temporal domain (as *n* increases) and are frequency-dependent. For each value of *f*_*spk*_, *V* (*t*_2_) = *V*_*trough*,1_ *> V* (*t*_1_) = 0 and therefore, *V*_*peak*,2_ *> V*_*peak*,1_. This causes *V*_*trough*,2_ = *V*_*trough*,1_. Following this process for increasing values of *n*, leads to two monotonically non-decreasing sequences converging to *V*_*peak*_ and *V*_*trough*_. As *f*_*spk*_ increases, *V*_*trough,n*_ also increase for fixed values of *n*, since there is less time for *V* to decay, and therefore *V*_*peak,n*_ also increase.

Therefore, *V*_*peak*_ and *V*_*trough*_ are increasing functions of *f*_*spk*_. If the *V*_*peak*_ profile increases slower than *V*_*trough*_ profile, then the Γ_*V*_ profile is a LPF (Fig. 9, column 1). For *τ* → 0, the *V*_*peak*_ profile is proportional to *S*_*peak*_ and *V*_*peak*_ → 0. As *τ* increases, both the *V*_*peak*_ and Γ_*V*_ profiles are amplified.

**Figure 9.**
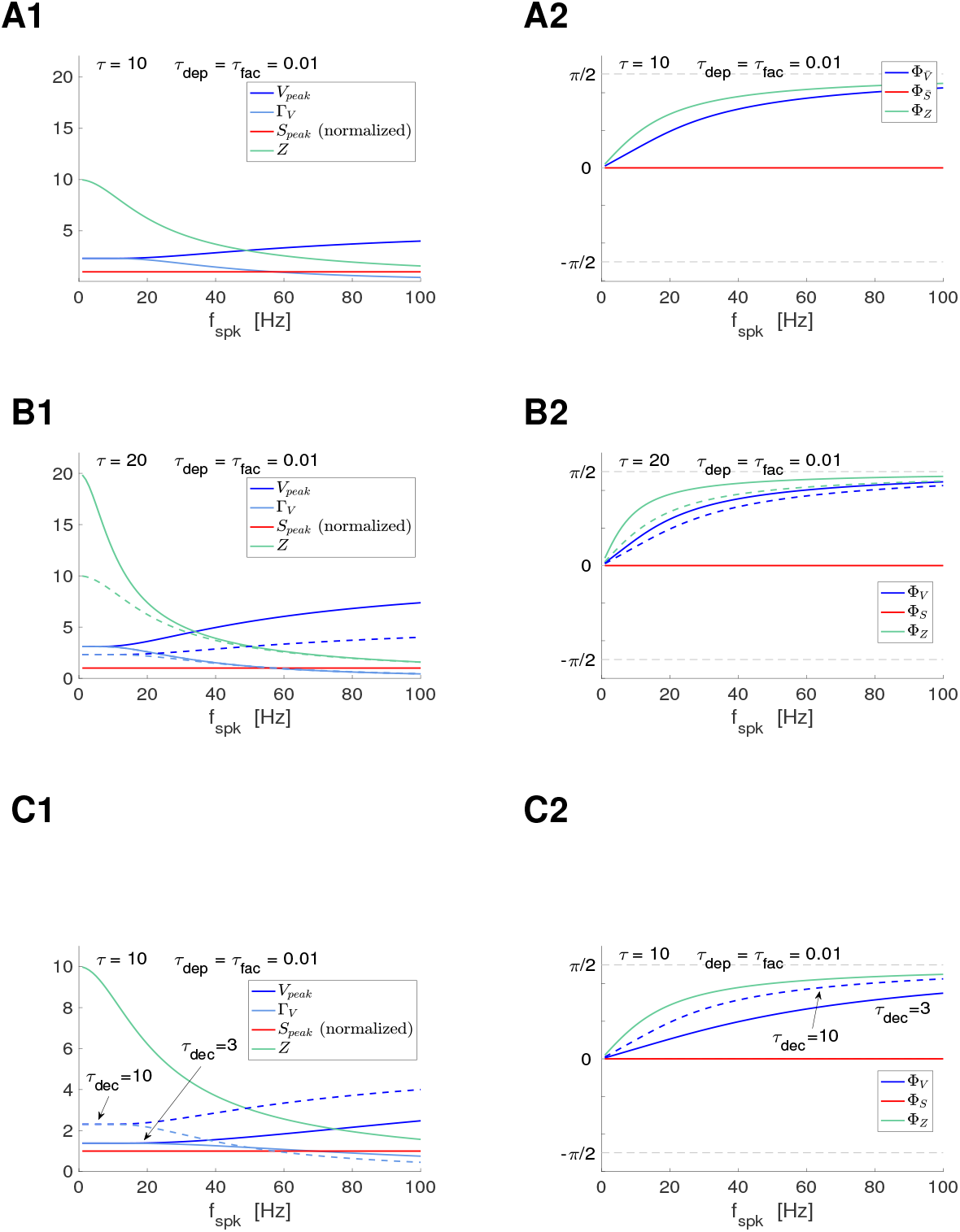
Postsynaptic filters in response to periodic presynaptic spike inputs for the passive (postsynaptic) cell in the absence of STP. We used eq. (65) for the PSP *V* with *I*_*syn*_ described by eqs. (2)-(4) appropriately adapted to account for the translation of *V* to the equilibrium point. We used eqs. (12) and (13) (DA model) with very values of *τ*_*dep*_ and *τ*_*fac*_ in the no STP regime. The impedance amplitude (*Z*) and phase (Φ_*Z*_) were computed using eqs. (39) and (40). The analytical approximations for the PSP peak sequence response of passive cells to presynaptic inputs are described in Section 2.2.4 (see also Appendix A). The approximation of *V*_*peak,n*_, *V*_*trough,n*_ and *t*_*V,peak*_ were computed as described in Section 2.2.4. The PSP amplitude Γ_*V*_ was computed by using eq. (41) and the PSP phase Φ_*V*_ was computed using eq. (42). The synaptic (*S*) peak (*S*_*peak*_) and phase (Φ_*S*_) profiles were computed similarly to these for *V*. **A**. *τ* = 10 (*G*_*L*_ = 0.1), *τ*_*dec*_ = 10. **B**. *τ* = 20 (*G*_*L*_ = 0.05), *τ*_*dec*_ = 10. The dashed curves correspond to panels B (*τ* = 10) and are presented for comparison purposes. **C**. *τ* = 10 (*G*_*L*_ = 0.1), *τ*_*dec*_ = 3. The dashed curves correspond to panels B (*τ*_*dec*_ = 10) and are presented for comparison purposes. We used the following additional parameter values: *C* = 1, *E*_*L*_ = −60, *I*_*app*_ = 0, *G*_*syn*_ = 0.1, *E*_*syn*_ = 0, *a*_*d*_ = 0.1, *a*_*f*_ = 0.1, *x*_∞_ = 1, *z*_∞_ = 0 and *T*_*sw*_ = 1.

### 3.5 The response of passive cells to direct (sinusoidal) and indirect (periodic presynaptic) inputs shows different filtering properties

Here and in the next sections, using the insights and intuition gained in Sections 3.3 and 3.4, we first discuss the PSP summation HPFs in passive cells (controlled by *τ* and modulated by *τ*_*dec*_) in the absence of STP, and their link to the passive cells’ LPFs (captured by *Z*, also controlled by *τ*). We then discuss the PSP filtering properties of passive cells in response to periodic presynaptic inputs in the presence of either depression (LPF) or facilitation (HPF).

The PSP filtering properties, captured by the *V*_*peak*_, Γ_*V*_ and Φ_*V*_ profiles, depend on the filtering properties of the participating building blocks and are controlled by the time constants operating at each level (Fig. 1): *τ*_*dep*_ and *τ*_*fac*_ (STP), *τ*_*dec*_ (synaptic) and *τ* (postsynaptic). From our results in Section 3.2 (see also Section 2.2.4), the steady-state *S* peak profiles *S*_*peak*_ (or 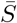) depend at most mildly on *τ*_*rse*_ for the type of fast synaptic rise times we consider here.

#### 3.5.1 PSP *V*_*peak*_ (LPFs; *Z*), Γ_*V*_ (LPFs; *Z*) and Φ_*Z*_ (delay) profiles in response to direct activation of oscillatory inputs (revisited)

Passive cells are LPFs in response to direct activation of sinusoidal input currents and always exhibit a delayed response (see Section 2.3.2). The *Z* (amplitude) profile (39) is a decreasing function of the input frequency *f* (Fig. 9-A1, green) and the Φ_*Z*_ profile (40) is a positive increasing function of *f*, converging to *π/*2 (Fig. 9-A2, green). These profiles are affected by the membrane time constant *τ* = *C/G*_*L*_. Increasing values of *τ* cause (i) an increase in *Z*_*max*_ = *Z*(0) (compare Figs. 9-A1 and B1, green), (ii) a sharper decrease of the *Z* profile (compare Figs. 9-A1 and B1, green), and (iii) a sharper increase of the Φ_*Z*_ profile (compare Figs. 9-A2 and B2, green).

#### 3.5.2 PSP *V*_*peak*_ (HPFs), Γ_*V*_ (LPFs) and Φ_*V*_ (delay) profiles in response to periodic presynaptic inputs

In response to periodic presynaptic inputs (with no STP), passive cells are Γ_*V*_ LPFs (Fig. 9, left column, light blue), but *V*_*peak*_ HPFs (Fig. 9, left column, blue), while always exhibit a delayed response (Fig. 9, right column, blue) similarly to Φ_*Z*_. The *V*_*peak*_ HPFs result from the summation phenomenon and are different in nature from the HPFs resulting from synaptic facilitation.

These profiles are modulated by both *τ* (membrane time constant) and *τ*_*dec*_ (synaptic decay time). Increasing values of *τ* cause (i) an amplification of the *V*_*peak*_ profiles (compare Figs. 9-A1 and B1, blue), (ii) a sharper increase of the *V*_*peak*_ profile with *f*_*spk*_ (compare Figs. 9-A1 and B1, blue), (iii) an amplification of the Γ_*V*_ profiles, which is more pronounced for the lower values of *f*_*spk*_ (compare Figs. 9-A1 and B1, light blue), (iv) a sharper decrease in the Γ_*V*_ profile with increasing values of *f*_*spk*_ (compare Figs. 9-A1 and B1, green), and (v) a sharper increase in the Φ_*V*_ profile with increasing values of *f*_*spk*_ (compare Figs. 9-A2 and B2, blue). Increasing values of *τ*_*dec*_ cause (i) an amplification of both the *V*_*peak*_ and Γ_*V*_ profiles (Fig. 9-C1, blue and light blue), (ii) a sharper decrease of the Γ_*V*_ profiles with increasing values of *f*_*spk*_ (Fig. 9-C1, light blue), and (iii) a sharper increase of the Φ_*V*_ profiles with increasing values of *f*_*spk*_ (Fig. 9-C2, blue).

### 3.6 PSP *V*_*peak*_ and Γ_*V*_ profiles: Inherited and cross-level mechanisms of generation of BPFs (PSP resonance)

The *V*_*peak*_, Γ_*V*_ and Φ_*V*_ profiles are shaped by the feedforward interaction of time scales and other constitutive properties across the participating building blocks (Fig. 1-B). Similarly to the BPFs generated by the simplified model used in Section 3.2, PSP BPFs can be inherited from one level (e.g, synaptic update) to another (PSP) or can be generated as the result of the interplay of LPFs and HPFs across levels.

For small enough values of *τ*_*dec*_ and *τ*, the *V*_*peak*_ profile is approximately proportional to the *S*_*peak*_ profile, the *V*_*trough*_ profile is almost zero, and therefore the Γ_*V*_ profile is approximately equal to the *V*_*peak*_ profile. In the limit of *τ* → 0 and *τ*_*dec*_ → 0 these relationships are strictly valid. In this sense, the PSP *V*_*peak*_ and Γ_*V*_ profiles (and the filtering properties) are inherited from the synaptic level *S*_*peak*_ profile.

If *τ*_*dec*_ or *τ* increase slightly, the *V*_*peak*_ and Γ_*V*_ profiles are modulated versions of the *S*_*peak*_ profiles. Specifically, if *τ*_*dec*_ increases (*τ ≪*1), then the *V*_*peak*_ profile remains almost proportional to *S*_*peak*_ profile, but the *V*_*trough*_ profile increases with *f*_*spk*_ and therefore the Γ_*V*_ profile is lower than the *V*_*peak*_ profile. If, on the other hand, *τ* increases (*τ*_*dec*_ *≪*1), the *V*_*peak*_ profile is no longer proportional to the *S*_*peak*_ profile and the *V*_*trough*_ profile is an increasing function of *f*_*spk*_, and therefore the Γ_*V*_ and *V*_*peak*_ profiles are different.

If *τ*_*dec*_ or *τ* increase further, then the *V*_*peak*_ and Γ_*V*_ profiles main remain modulated versions of the *S*_*peak*_ profiles or have a qualitatively different shape from the *S*_*peak*_ profiles. Similarly to the generation of the simplified STP-mediated 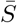 (PSP) BPFs discussed in Section 3.2, under certain balance conditions, synaptic and postsynaptic filters with opposite monotonic dependencies (with *f*_*spk*_) are expected to produce BPFs, while synaptic and postsynaptic filters with the same monotonic dependencies (with *f*_*spk*_) are expected to reinforce each other.

The interplay of synaptic depression (LPF) and *V*_*peak*_ summation (HPF) is able to generate *V*_*peak*_ BPFs (Fig. 10-A1, solid blue), but not Γ_*V*_ BPFs (Fig. 10-A, solid light blue) since both the *S*_*peak*_ and the Γ_*V*_ profiles are LPFs. In contrast, the interplay of synaptic facilitation (HPF) and Γ_*V*_ summation (LPF) is able to generate Γ_*V*_ BPFs (Fig. 10-B, solid light blue), but not *V*_*peak*_ BPFs since both the *S*_*peak*_ and *V*_*peak*_ summation profiles are HPFs (Fig. 10-B, solid blue). In the presence of both synaptic depression (LPF) and facilitation (HPF), the *S*_*peak*_ profiles are BPFs (Fig. 10-C, solid red). These are communicated to the PSP level where they are modulated by the postsynaptic membrane potential properties. In certain parameter regimes, these modulations produce PSP *V*_*peak*_ and Γ_*V*_ BPFs (Fig. 10-C, solid blue).

**Figure 10.**
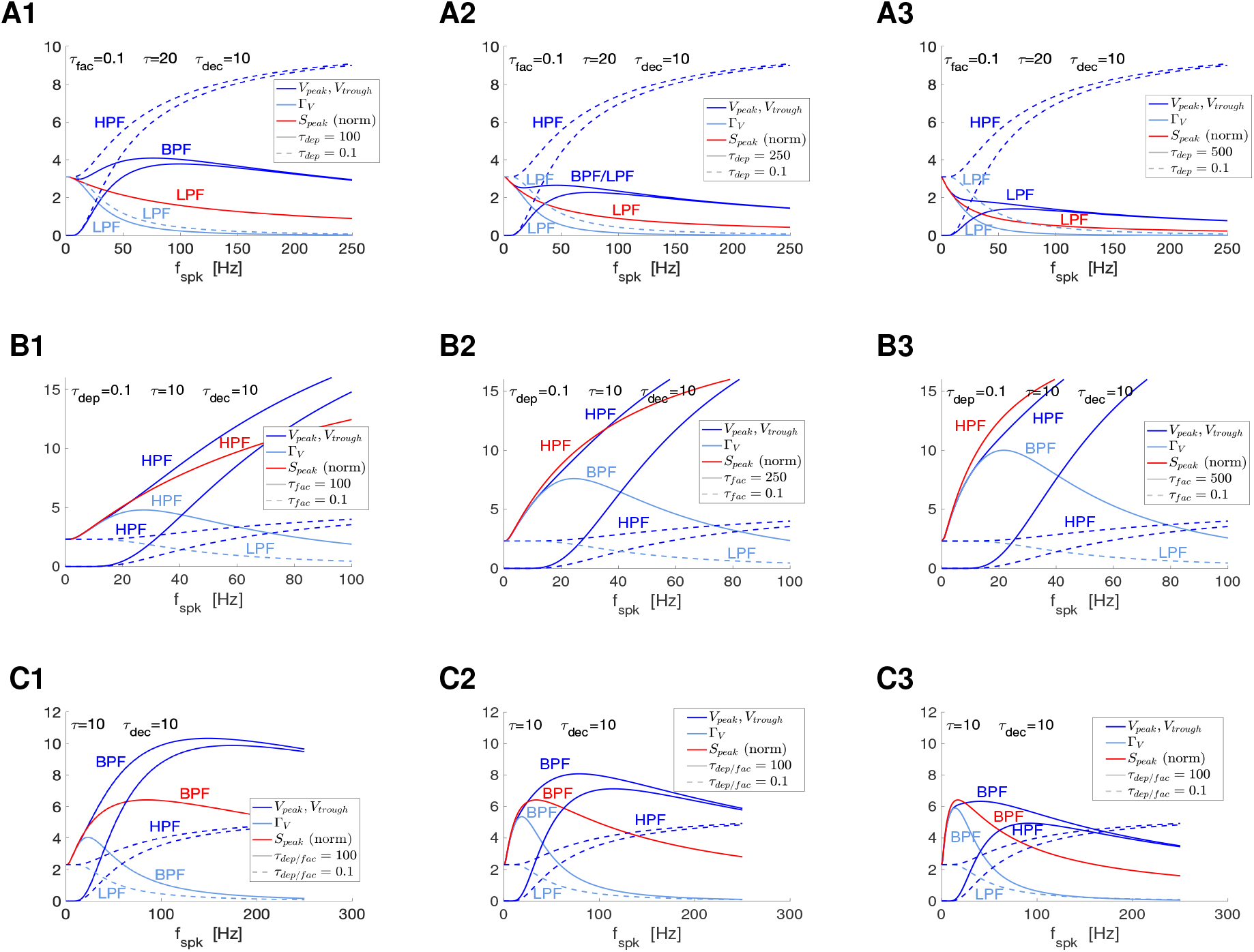
Postsynaptic BPFs filters in response to periodic presynaptic spike inputs emerging from the interplay of short-term depression (LPF), short-term facilitation (HPF) and postsynaptic summation (HPF). Each panel shows the superimposed *V*_*peak*_, *V*_*trough*_ and Γ_*V*_ profiles in response to presynaptic inputs in the presence (solid) and absence (dashed) of STP. The *S*_*peak*_ profiles are normalized to coincide with the *V*_*peak*_ profiles for the lowest value of *f*_*spk*_ (*f*_*spk*_ = 0.1 in the simulations). The summation HPFs are generated in response to presynaptic inputs in the absence of STP (*τ*_*dep*_ = 0.1, *τ*_*fac*_ = 0.1 or *τ*_*dep*_ = *τ*_*fac*_ = 0.1 in the simulations). The *S*_*peak*_ profiles in the absence of STP are horizontal lines (not shown). **A**. *V*_*peak*_ BFPs (solid blue) generated by the interplay of *S*_*peak*_ LPFs (red) and *V*_*peak*_ summation HPFs (dashed blue) in the presence of synaptic depression only. The interplay of the Γ_*V*_ summation LPF (dashed light blue) and the *S*_*peak*_ LPF (red) produces a Γ_*V*_ LPF (solid light blue). **B**. Γ_*V*_ BFPs (light blue) generated by the interplay of *S*_*peak*_ HPFs (red) and Γ_*V*_ summation LPFs (dashed blue) in the presence of synaptic facilitation only. The interplay of of the *V*_*peak*_ summation HPF (dashed blue) and the *S*_*peak*_ HPF (red) produces a *V*_*peak*_ HPF (solid blue). **C**. *V*_*peak*_ BPFs (solid blue) and Γ_*V*_ BPFs (solid light blue) generated by the interplay of the inherited *S*_*peak*_ BFPs (red) and modulated by the *V*_*peak*_ summation HPFs (dashed blue) and the *Gamma*_*V*_ summation LPFs (dashed light blue). We used eq. (65) for the PSP *V* with *I*_*syn*_ described by eqs. (2)-(4) appropriately adapted to account for the translation of *V* to the equilibrium point, and STP described by eqs. (12) and (13) (DA model). The impedance amplitude (*Z*) and phase (Φ_*Z*_) were computed using eqs. (39) and (40). The analytical approximations for the PSP peak sequence response of passive cells to presynaptic inputs are described in Section2.2.4 (see also Appendix A). The approximation of *V*_*peak,n*_ and *V*_*trough,n*_ were computed as described in Section 2.2.4. The PSP amplitude Γ_*V*_ was computed by using eq. (41) and the PSP phase Φ_*V*_ was computed using eq. (42). The synaptic (*S*) peak (*S*_*peak*_) and phase (Φ_*S*_) profiles were computed similarly to these for *V*. We used the following additional parameter values: *C* = 1, *E*_*L*_ = −60, *I*_*app*_ = 0, *G*_*syn*_ = 0.1, *E*_*syn*_ = 0, *a*_*d*_ = 0.1, *a*_*f*_ = 0.1, *x*_∞_ = 1, *z*_∞_ = 0 and *T*_*sw*_ = 1.

We analyze the various possible scenarios and mechanisms in the next sections.

### 3.7 Interplay of STD-mediated LPFs and PSP SUM-mediated HPFs: *V*_*peak*_ BPFs and Γ_*V*_ LPFs

The presence of synaptic STD generates *S*_*peak*_ LPFs that become sharper and more attenuated as *τ*_*dep*_ increases (Fig. 11-A3) and are independent of *τ, G*_*syn*_ and *τ*_*dec*_ (Fig. 11-B3 to -D3)

**Figure 11.**
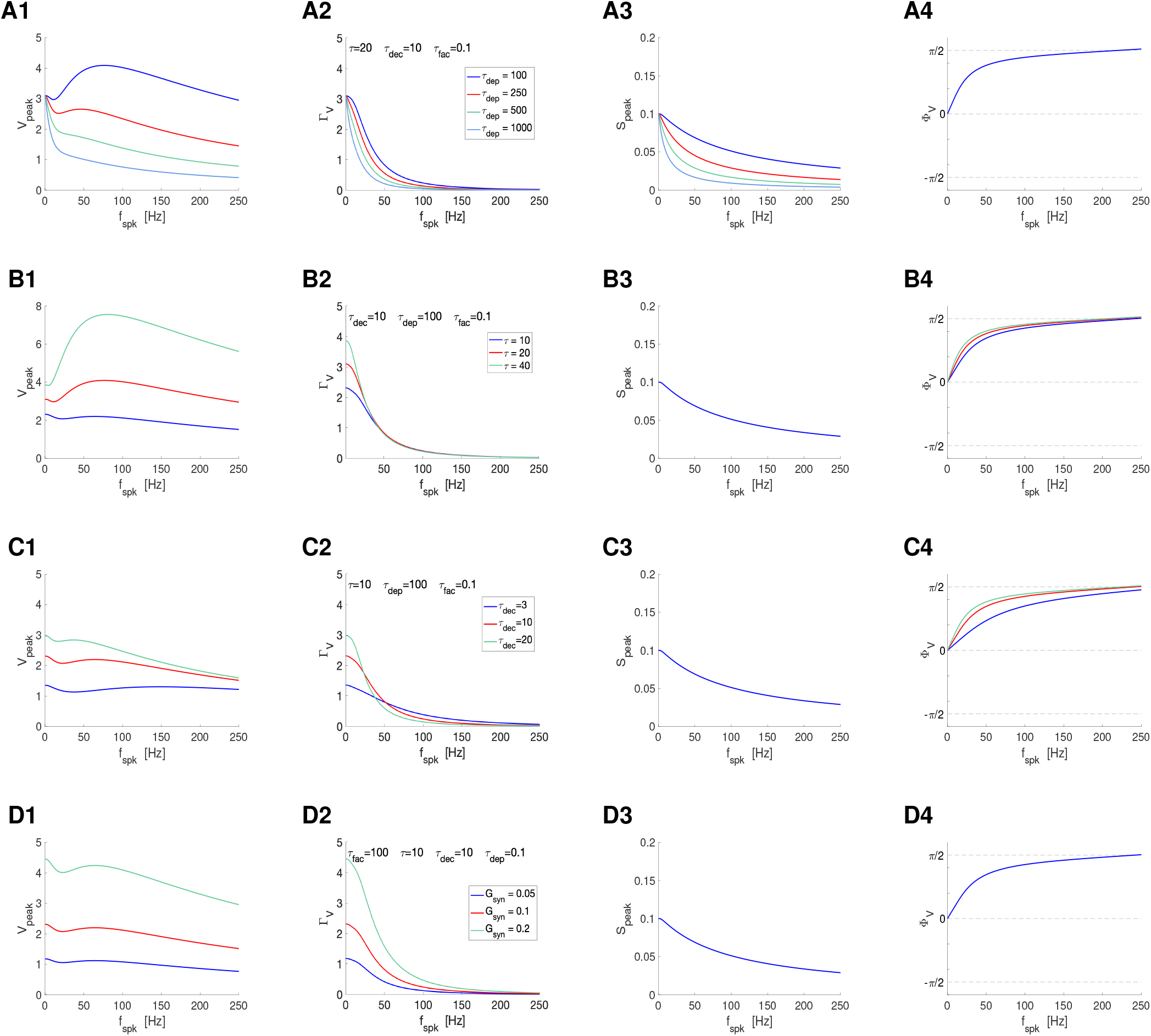
Postsynaptic filters in response to periodic presynaptic spike inputs emerging from the interplay of short-term depression and postsynaptic summation. **A**. Superimposed filters for various values of the short-term depression time constant *τ*_*dep*_ and representative parameter values: *τ* = 10, *τ*_*dec*_ = 10 and *τ*_*fac*_ = 0.1. Fig. S5 extends these results for additional values of *τ*. The Φ_*V*_ profiles are independent of *τ*_*dep*_. **B**. Superimposed filters for various values of the postsynaptic membrane time constant *τ* and representative parameter values: *τ*_*dec*_ = 10, *τ*_*dep*_ = 100 and *τ*_*fac*_ = 0.1. Fig. S6 extends these results for additional values of *τ*_*dep*_. The *S*_*peak*_ profiles are independent of *τ*_*dep*_. **C**. Superimposed filters for various values of the synaptic decay time *τ*_*dec*_ and representative parameter values: *τ* = 10, *τ*_*dep*_ = 100 and *τ*_*fac*_ = 0.1. The *S*_*peak*_ profiles are independent of *τ*_*dep*_. Fig. S7 extends these results for additional values of *τ*_*dep*_. **D**. Superimposed filters for representative values of the synaptic decay time constant *G*_*syn*_ and representative parameer values: *τ*_*fac*_ = 100, *τ* = 10, *τ*_*dec*_ = 10 and *τ*_*dep*_ = 0.1. Fig. S10 extends these results for additional values of *τ*_*fac*_. The *S*_*peak*_ and Φ_*V*_ profiles are independent of *G*_*syn*_. **Left column**. *V* peak profiles. **Middle-left column**. *V* peak-to-trough amplitude profiles. **Middle-right column**. *S* peak profiles. **Right column**. *V* phase profiles. We used eq. (65) for the PSP *V* with *I*_*syn*_ described by eqs. (2)-(4) appropriately adapted to account for the translation of *V* to the equilibrium point, and STP described by eqs. (12) and (13) (DA model). The impedance amplitude (*Z*) and phase (Φ_*Z*_) were computed using eqs. (39) and (40). The analytical approximations for the PSP peak sequence response of passive cells to presynaptic inputs are described in Section2.2.4 (see also Appendix A). The approximation of *V*_*peak,n*_, *V*_*trough,n*_ and *t*_*V,peak*_ were computed as described in Section 2.2.4. The PSP amplitude Γ_*V*_ was computed by using eq. (41) and the PSP phase Φ_*V*_ was computed using eq. (42). The synaptic (*S*) peak (*S*_*peak*_) and phase (Φ_*S*_) profiles were computed similarly to these for *V*. We used the following additional parameter values: *C* = 1, *E*_*L*_ = −60, *I*_*app*_ = 0, *G*_*syn*_ = 0.1, *E*_*syn*_ = 0, *a*_*d*_ = 0.1, *a*_*f*_ = 0.1, *x*_∞_ = 1, *z*_∞_ = 0 and *T*_*sw*_ = 1.

#### 3.7.1 Emergence of *V*_*peak*_ resonance (BPFs): interplay of STD-mediated LPF and a PSP summation-mediated HPF

The interaction between these STD-mediated LPFs and the PSP summation-mediated HPF (Fig. 9-A1, blue) produces *V*_*peak*_ BPFs for values of *τ*_*dep*_ within some range (Fig. 11-A1, blue and red). We refer to this preferred frequency PSP peak response to periodic presynaptic inputs as (PSP) *V*_*peak*_ resonance. This maximal amplification of the *V*_*peak*_ response is often preceded by a relatively small trough. *V*_*peak*_ resonance reflects balances between the two participating processes. As *τ*_*dep*_ increases (sharper decrease of the *S*_*peak*_ profile), the *V*_*peak*_ profiles are dominated by depression and therefore they are attenuated as they transition to LPFs for larger values of *τ*_*dep*_ (Fig. 11-A1, blue to light blue). This is accompanied by a decrease in the *V*_*peak*_ resonant frequency. As *τ* increases, the *V*_*peak*_ profiles are dominated by summation and therefore they are amplified (Fig. 11-A2). This is accompanied by an increase in the *V*_*peak*_ resonant frequency. Changes in *τ* and *G*_*syn*_ do not affect the *S*_*peak*_ profiles, but they affect the *V* response to presynaptic spikes in a frequency-dependent manner (Fig. 8). Consistently with that, increasing values of *τ* and *G*_*syn*_ amplify the *V*_*peak*_ response with lesser effects on their shapes than changes in *τ*_*dep*_ and *τ* (Fig. 11-C1 and -D1). This is more prominent for *G*_*syn*_, which has almost a multiplicative effect on the *V*_*peak*_ profiles, than for *τ*, consistently with the different ways in which they control the synaptic currents.

#### 3.7.2 Modulation of Γ_*V*_ LPFs: interplay of STD and PSP amplitude LPFs

In the absence of STP, the Γ_*V*_ LPF is controlled by the membrane time constant *τ* (Fig. 9-B1) and the synaptic decay time constant *τ*_*dec*_ (Fig. 9-C1). Here we focus on the effects of synaptic depression (*τ*_*dep*_) and the interplay between the three time constants in the modulation of Γ_*V*_. Increasing values of *τ*_*dep*_ sharpen Γ_*V*_ without affecting Γ_*V*_ (0) (Fig. 11-A2). The magnitude of the modulation depend on the other parameter values (Fig. S5, column 2). Increasing values of *τ* and *τ*_*dec*_ sharpen Γ_*V*_ and increase Γ_*V*_ (0) (Fig. 11-B2 and -C2). This is at most mildly affected by *τ*_*dep*_ (Fig. S7, column 2) and *τ*_*dec*_ (Fig. S8, column 2). Increasing values of *G*_*syn*_ have a multiplicative effect on Γ_*V*_ (Fig. 11-D2).

#### 3.7.3 Modulation of the Φ_*V*_

In the absence of STP, the Φ_*V*_ profiles are controlled by *τ* and *τ*_*dec*_ (Fig. 9). Changes in *τ*_*dep*_ do not affect Φ_*V*_ (Fig. 11-A4 and S5). Changes in *τ* and *τ*_*dec*_ affect Φ_*V*_ in the same direction as in the absence of STP (Figs. 11-B4, -A4, S6 and S7). Consistently with the previous findings, changes in *G*_*syn*_ do not affect Φ_*V*_.

Figs. S5-S7 extend our results for additional combinations of parameter values. Figs. S5-S7 (left column), in particular, illustrate how the balances between the depression LPFs and peak summation HPFs shape the *V*_*peak*_ profiles for additional parameter regimes.

@@@

### 3.8 Interplay of STF-mediated HPFs and PSP SUM-mediated filters: *V*_*peak*_ LPFs and Γ_*V*_ BPFs

The presence of STF generates *S*_*peak*_ HPFs that become sharper and more amplified as *τ*_*fac*_ increases (Fig. 12-A3) and are independent of *τ* and *G*_*syn*_ (Fig. 12-B3 to -D3).

**Figure 12.**
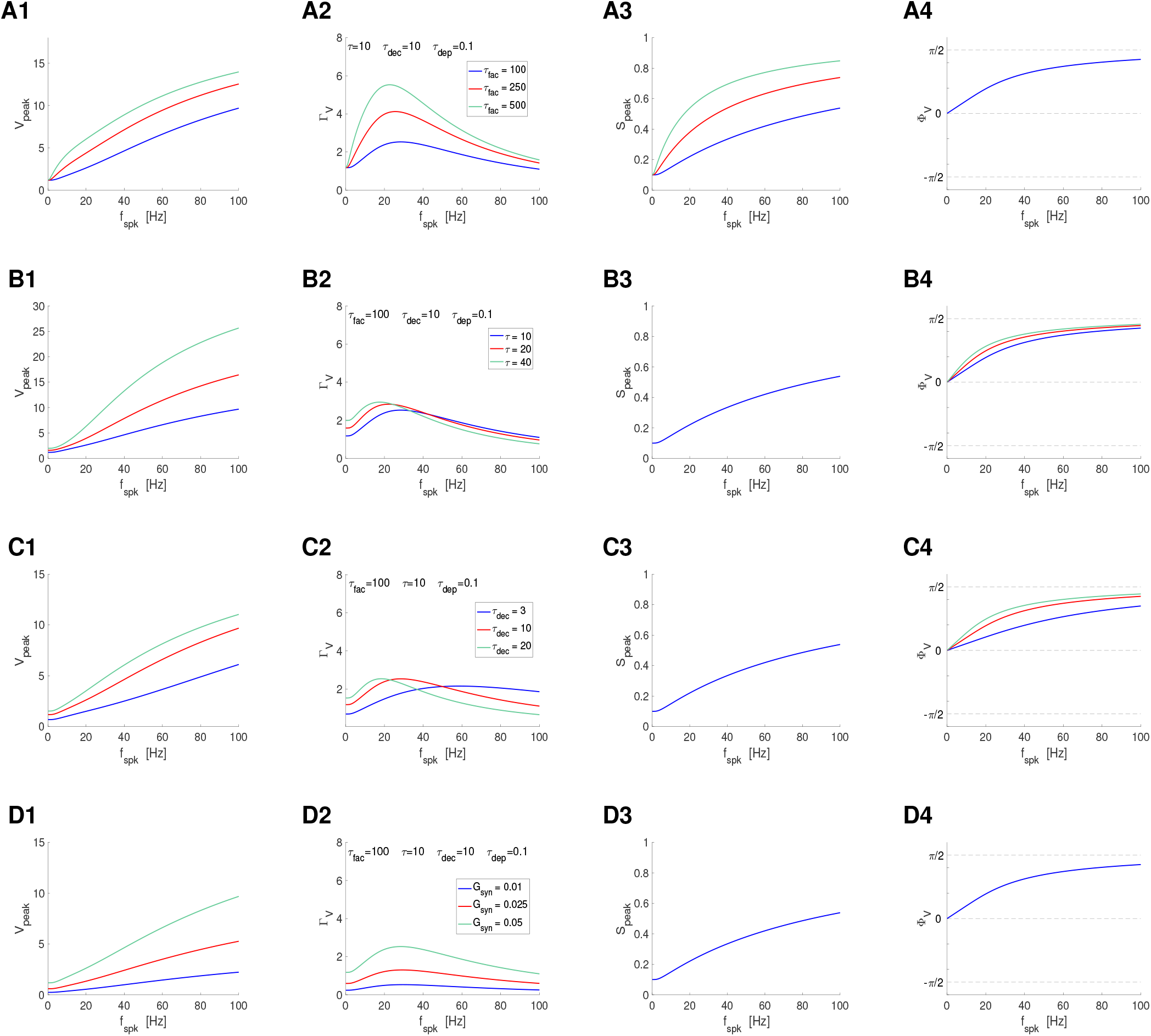
Postsynaptic filters in response to periodic presynaptic spike inputs emerging from the interplay of short-term facilitation and postsynaptic summation. **A**. Superimposed filters for various values of the short-term depression time constant *τ*_*fac*_ and representative parameter values: *τ* = 10, *τ*_*dec*_ = 10 and *τ*_*dep*_ = 0.1. Fig. S8 extends these results for additional values of *τ*. The Φ_*V*_ profiles are independent of *τ*_*fac*_. **B**. Superimposed filters for various values of the membrane time constant *τ* and representative parameter values: *τ*_*fac*_ = 100, *τ*_*dec*_ = 10 and *τ*_*dep*_ = 0.1. Fig. S9 extends these results for additional values of *τ*_*fac*_. The *S*_*peak*_ profiles are independent of *τ*. **C**. Superimposed filters for representative values of the synaptic decay time constant *τ*_*dec*_ and representative parameer values: *τ*_*fac*_ = 100, *τ* = 10 and *τ*_*dep*_ = 0.1. Fig. S10 extends these results for additional values of *τ*_*fac*_. The *S*_*peak*_ profiles are independent of *τ*_*dec*_. **D**. Superimposed filters for representative values of the synaptic decay time constant *G*_*syn*_ and representative parameer values: *τ*_*fac*_ = 100, *τ* = 10, *τ*_*dec*_ = 10 and *τ*_*dep*_ = 0.1. Fig. S10 extends these results for additional values of *τ*_*fac*_. The *S*_*peak*_ and Φ_*V*_ profiles are independent of *G*_*syn*_. **Left column**. *V* peak profiles. **Middle-left column**. *V* peak-to-trough amplitude profiles. **Middle-right column**. *S* peak profiles. **Right column**. *V* phase profiles. We used eq. (65) for the PSP *V* with *I*_*syn*_ described by eqs. (2)-(4) appropriately adapted to account for the translation of *V* to the equilibrium point, and STP described by eqs. (12) and (13) (DA model). The impedance amplitude (*Z*) and phase (Φ_*Z*_) were computed using eqs. (39) and (40). The analytical approximations for the PSP peak sequence response of passive cells to presynaptic inputs are described in Section2.2.4 (see also Appendix A). The approximation of *V*_*peak,n*_, *V*_*trough,n*_ and *t*_*V,peak*_ were computed as described in Section 2.2.4. The PSP amplitude Γ_*V*_ was computed by using eq. (41) and the PSP phase Φ_*V*_ was computed using eq. (42). The synaptic (*S*) peak (*S*_*peak*_) and phase (Φ_*S*_) profiles were computed similarly to these for *V*. We used the following additional parameter values: *C* = 1, *E*_*L*_ = −60, *I*_*app*_ = 0, *G*_*syn*_ = 0.05, *E*_*syn*_ = 0, *a*_*d*_ = 0.1, *a*_*f*_ = 0.1, *x*_∞_ = 1, *z*_∞_ = 0 and *T*_*sw*_ = 1.

#### 3.8.1 Modulation of *V*_*peak*_ HPFs: interplay of STF- and PSP SUM-mediated HPFs

In the absence of STP, the *V*_*peak*_ HPF is controlled by the membrane time constant *τ* (Fig. 9-B1) and the synaptic decay time constant *τ*_*dec*_ (Fig. 9-C1). Here we focus on the effects of synaptic facilitation (*τ*_*fac*_) and the interplay between the three time constants in the modulation of *V*_*peak*_. Increasing values of *τ*_*fac*_ cause an amplification of the *S*_*peak*_ profiles (Fig. 12-A3) and therefore an amplification of the *V*_*peak*_ profiles (Fig. 12-A1). Increasing values of *τ*_*dec*_ and *G*_*syn*_ increase *I*_*syn*_ in a frequency-dependent manner, and therefore they also amplify the *V*_*peak*_ profiles (Fig. 12-C1 and -D1). Finally, consistently with our findings in Fig. 8-A, increasing values of *τ* also amplify the *V*_*peak*_ profiles.

#### 3.8.2 Emergence of Γ_*V*_ resonance (BPFs): interplay of a STF-mediated HPF and a PSP SUM-mediated amplitude LPF

The presence of synaptic STF generates *S*_*peak*_ HPFs that become sharper and more amplified as *τ*_*fac*_ increases (Fig. 12-A3). The interaction between these filters and the PSP amplitude LPFs (Fig. 9-A1, light-blue) produces BPFs (Fig. 11-A2). We refer to this preferred frequency PSP amplitude response to periodic presynaptic inputs as (PSP) Γ_*V*_ resonance. Similarly to other resonances, Γ_*V*_ resonance reflects balances between the two participating process. As *τ*_*fac*_ increases, the Γ_*V*_ profiles are more dominated by facilitation and therefore the Γ_*V*_ profiles are amplified as the Γ_*V*_ resonant frequency decreases (Fig. 12-A2).

As *τ* increases, the STP-independent Γ_*V*_ LPFs are amplified and sharpened (Fig. 9-B1). Therefore, increasing values of *τ* amplify the Γ_*V*_ BPFs and shift the resonant frequency to lower values (Fig. 11-B2). Similarly, increasing values of *τ*_*dec*_, which sharpen the STP-independent Γ_*V*_ LPFs (Fig. 9-C1), shift the facilitation-induced Γ_*V*_ resonant frequency to lower values and amplifies the Γ_*V*_ BPFs within certain range of values of *τ* (Fig. 12-C2). Increasing values of *G*_*syn*_ in contrast amplify the Γ_*V*_ profile with a much lesser effect on the Γ_*V*_ resonant frequency (Fig. 12-D2)

#### 3.8.3 Modulation of the Φ_*V*_

In the absence of STP, the Φ_*V*_ profile is controlled by *τ* and *τ*_*dec*_ (Fig. 9). Similarly to our findings in the previous section, changes in *τ*_*fac*_ do not affect Φ_*V*_ (Fig. 12-A4 and S8). Changes in *τ* and *τ*_*dec*_ affect Φ_*V*_ in the same direction as in the absence of STP (Figs. 12-B4, -A4, S9 and S10). Consistently with the previous findings, changes in *G*_*syn*_ do not affect Φ_*V*_ (Fig. 12-D4).

Figs. S8-S10 extend our results for additional parameter combinations. Figs. S8-S10 (middle-left column), in particular, illustrate how the balances between the facilitation HPFs and amplitude LPFs shape the Γ_*V*_ profiles for additional parameter regimes.

### 3.9 Interplay of STD-mediated LPFs, STF-mediated HPFs and PSP SUM-mediated filters HPFs: *V*_*peak*_ and Γ_*V*_ BPFs

The presence of synaptic STD and STF generates *S*_*peak*_ BPFs (synaptic resonance) that become sharper as *τ*_*dep*_ and *τ*_*fac*_ increase (Fig. 13-A3). This is accompanied by a decrease in the synaptic resonance frequency. The *S*_*peak*_ BPFs are independent of *τ, τ*_*dec*_ and *G*_*syn*_ (Fig. 13-B3 to -D3). As discussed above, for small enough values of *τ*_*dec*_ and *τ*, the *S*_*peak*_ profiles are inherited to the postsynaptic level and therefore the PSP *V*_*peak*_ and Γ_*V*_ profiles are almost identical, and are almost proportional to the *S*_*peak*_ profiles (not shown). For larger values of *τ*_*dec*_ and *τ*, the PSP *V*_*peak*_ and Γ_*V*_ profiles are modulated by the PSP membrane properties and the SUM-mediated HPF.

**Figure 13.**
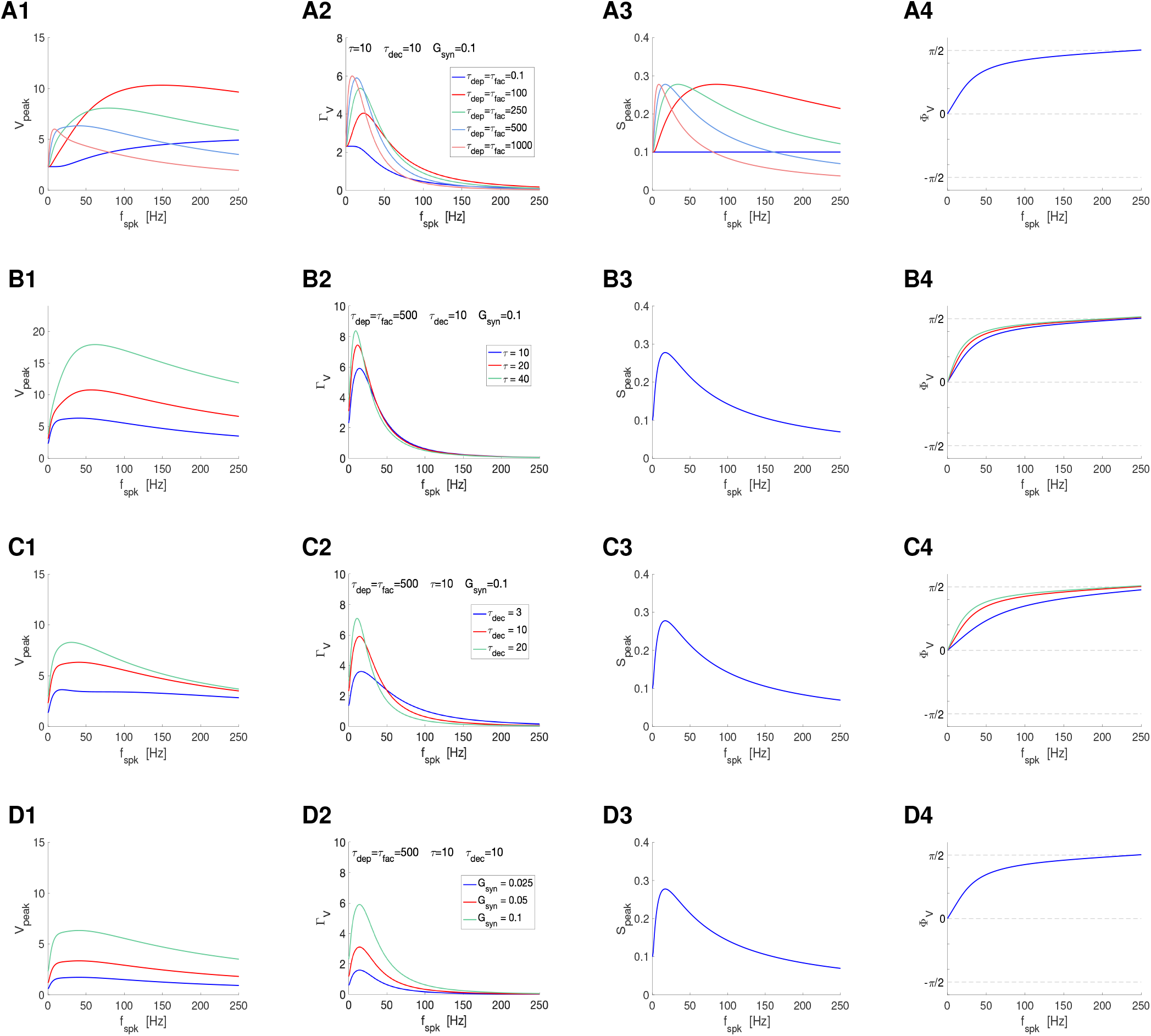
Postsynaptic filters in response to periodic presynaptic spike inputs emerging from the interplay of short-term depression, facilitation and postsynaptic summation. **A**. Superimposed filters for representative values of the depression and facilitation time constants *τ*_*dep*_ and *τ*_*fac*_, respectively, and representative parameter values: *τ* = 10, *τ*_*dec*_ = 10 and *G*_*syn*_ = 0.1. Fig. S11 extends these results for additional values of *τ*. The Φ_*V*_ profiles are independent of *τ*_*dep*_ and *τ*_*fac*_. **B**. Superimposed filters for representative values of the depression membrane time constant *τ* and representative parameter values: *τ*_*dep*_ = 500, *τ*_*fac*_ = 500, *τ*_*dec*_ = 10 and *G*_*syn*_ = 0.1. Fig. S12 extends these results for additional values of *τ*. The *S*_*peak*_ profiles are independent of *τ*. **C**. Superimposed filters for representative values of the depression synaptic decay time *τ*_*dec*_ and representative parameter values: *τ*_*dep*_ = 500, *τ*_*fac*_ = 500, *τ* = 10 and *G*_*syn*_ = 0.1. Fig. S13 extends these results for additional values of *τ*_fac_. The *S*_*peak*_ profiles are independent of *τ*_dec_. **D**. Superimposed filters for representative values of the synaptic decay time constant *G*_*syn*_ and representative parameer values: *τ*_*dep*_ = *τ*_*fac*_ = 500, *τ* = 10, *τ*_*dec*_ = 10. Fig. S10 extends these results fo **Left column**. *V* peak profiles. **Middle-left column**. *V* peak-to-trough amplitude profiles. **Middle-right column**. *S* peak profiles. **Right column**. *V* phase profiles. We used eq. (65) for the PSP *V* with *I*_*syn*_ described by eqs. (2)-(4) appropriately adapted to account for the translation of *V* to the equilibrium point, and STP described by eqs. (12) and (13) (DA model). The impedance amplitude (*Z*) and phase (Φ_*Z*_) were computed using eqs. (39) and (40). The analytical approximations for the PSP peak sequence response of passive cells to presynaptic inputs are described in Section 2.2.4 (see also Appendix A). The approximation of *V*_*peak,n*_, *V*_*trough,n*_ and *t*_*V,peak*_ were computed as described in Section 2.2.4. The PSP amplitude Γ_*V*_ was computed by using eq. (41) and the PSP phase Φ_*V*_ was computed using eq. (42). The synaptic (*S*) peak (*S*_*peak*_) andphase (Φ_*S*_) profiles were computed similarly to these for *V*. We used the following additional parameter values: *C* = 1, *E*_*L*_ = −60, *I*_*app*_ = 0, *E*_*syn*_ = 0, *a*_*d*_ = 0.1, *a*_*f*_ = 0.1, *x*_∞_ = 1, *z*_∞_ = 0 and *T*_*sw*_ = 1.

#### 3.9.1 Modulation of *V*_*peak*_ BPFs

Earlier studies modeled the PSP response of cells to periodic presynaptic spike inputs (*V*_*peak*_ profiles) in the presence of STP to be proportional to the *S*_*peak*_ profiles [30, 48, 93]. For larger, more realistic values of *τ* and *τ*_*dec*_ the *V*_*peak*_ profiles are wider than the *S*_*peak*_ profiles (compare Figs. 13-A1 and -A3) and the *V*_*peak*_ resonant frequency is larger than the *S*_*peak*_ resonant frequency. The *V*_*peak*_ profiles are amplified by decreasing values of *τ*_*dep*_ and *τ*_*fac*_. The amplification is stronger as *τ* increases (Fig. S11, column 1). The *V*_*peak*_ amplification is accompanied by an increase in the *V*_*peak*_ resonant frequency as *τ*_*dep*_ and *τ*_*fac*_ decrease. The *V*_*peak*_ profiles are also amplified by increasing values of *τ, τ*_*dec*_ and *G*_*syn*_ (Figs. 13, column 1). Increasing values of *τ* and *τ*_*dec*_ cause an increase in the *V*_*peak*_ resonant frequency (Figs. 13-B1 and -C1, see also Figs. S12 and S13, column 1), but the *V*_*peak*_ resonant frequency is at most slightly affected by increasing values of *G*_*syn*_ (Figs. 13-D1).

#### 3.9.2 Modulation of Γ_*V*_ BPFs

The Γ_*V*_ profiles are amplified and sharpened by increasing values of *τ*_*dep*_ and *τ*_*fac*_ (Fig. 13-A2). This is more pronounced as *τ* increases (Fig. S11, column 2). This is accompanied by a decrease in the Γ_*V*_ resonant frequency. The Γ_*V*_ profiles are also amplified by increasing values of *τ, τ*_*dec*_ and *G*_*syn*_. In the former two cases they are also sharpened.

#### 3.9.3 Modulation of the Φ_*V*_

In the absence of STP, the Φ_*V*_ profile is controlled by *τ* and *τ*_*dec*_ (Fig. 9). Consistent with our findings in the previous two sections, changes in *τ*_*dep*_ and *τ*_*fac*_ do not affect Φ_*V*_ (Fig. 13-A4 and S11). Changes in *τ* and *τ*_*dec*_ affect Φ_*V*_ in the same direction as in the absence of STP (Figs. 13-B4, -A4, S11 and S12). Consistent with the previous findings, changes in *G*_*syn*_ do not affect Φ_*V*_ (Figs. 13-D4)

Figs. S11-S13 extend our results for additional parameter combinations.

### 3.10 Persistence and modulation of STP-mediated PSP *V*_*peak*_ and Γ_*V*_ BPFs in response to randomly-distributed spike trains

In the previous sections we used periodic presynaptic inputs over a range of spiking frequencies to describe a number of mechanisms of generation of STP-mediated PSP *V*_*peak*_ and Γ_*V*_ BPFs both inherited from the *S* level of organization (*S*_*peak*_ BPFs) and generated by filters of opposing types across levels of organization.

The question arises whether PSP *V*_*peak*_ and Γ_*V*_ filters emerge in more general scenarios, in response to more realistic presynaptic spike trains having some frequency content, what are the properties of these filters, how they are affected by the input variability, and how they are related to the classical (deterministic) filters in response to periodic inputs. We address these questions by using two types of presynaptic spike inputs: jittered-periodic spike trains [46] and Poisson-distributed spike trains [86, 99].

#### 3.10.1 PSP *V*_*peak*_ and Γ_*V*_ BPFs persist in response to randomly perturbed periodic spike trains

Following [46], we consider perturbations of periodic presynaptic spiking patterns with ISIs of the form (6) for *n* = 1, …, *N*_*spk*_, where Δ_*spk*_ is constant (*n*-independent) and 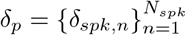 is a random variable with zero mean and variance equal to *δ* Δ_*spk*_ for a non-negative real number *δ*.

Fig. 14-A illustrates that PSP *V*_*peak*_ and Γ_*V*_ filters discussed above persist in response to the jittered-periodic presynaptic spike trains. The solid curves correspond to the mean value for each attribute (*V*_*peak*_, *V*_*trough*_, *S*_*peak*_ and Γ_*V*_) and the dashed-gray curves correspond to the classical, unperturbed filters. Fig. 14-A illustrates that these two quantities almost coincide. This also occurs for the *X*_*peak*_, *Z*_*peak*_ and Δ*S*_*peak*_ filters (Fig. S16). The variability of the filters is larger for *V*_*peak*_ and *V*_*trough*_ than for the other filters, and is frequency-dependent (compare the shadow regions for each filter across frequencies) and STP-dependent (compare the shadow regions for each input frequency across values of *τ*_*dep*_ and *τ*_*fac*_).

**Figure 14.**
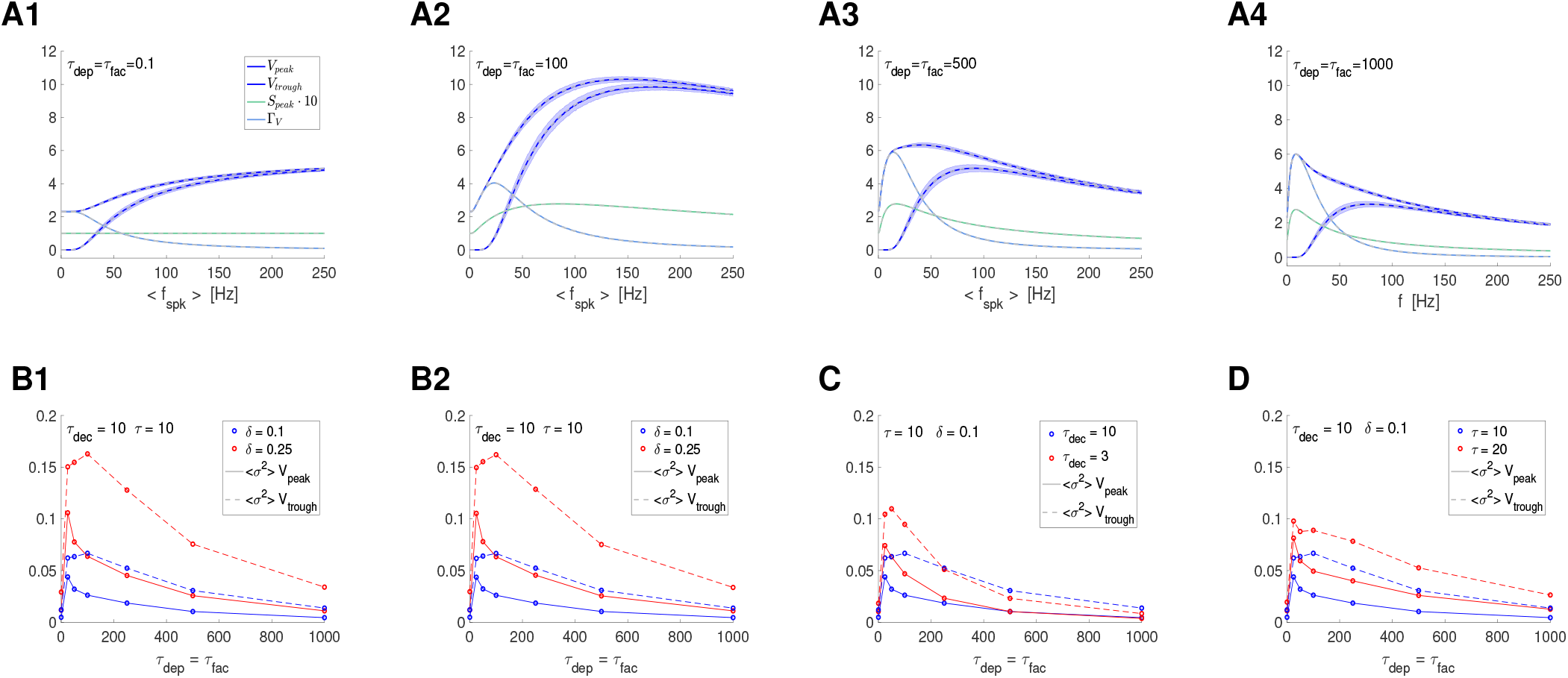
Postsynaptic filters in response to jittered (randomly perturbed) periodic presynaptic inputs in the presence of STP: frequency- and STP-dependent variability. For each value of the mean presynaptic input frequency *< f*_*spk*_ *>*, the ISI sequence *{*Δ_*spk,n*_*}* (*n* = 1, …, *N*_*spk*_) has the form Δ_*spk,n*_ = Δ_*spk*_ + *δ*_*spk,n*_ where Δ_*spk*_ is the ISI corresponding to *f*_*spk*_ (*f*_*spk*_ = 1000*/*Δ_*spk*_) and the sequence *{δ*_*spk,n*_*}* are drawn from a normal distribution with zero mean and variance equal to *δ* Δ_*spk*_. **A**. Superimposed *V*_*peak*_, *V*_*trough*_, *S*_*peak*_ and Γ_*V*_ profiles for representative parameter values. We used *τ*_*dec*_ = 10 and *τ* = 10 in all panels. Solid curves correspond to the mean values for each attribute (*V*_*peak*_, *V*_*trough*_, *S*_*peak*_ and Γ_*V*_). The shadow regions (*V*_*peak*_, *V*_*trough*_ and *S*_*peak*_ correspond to one standard deviation from the mean. The dashed gray curves, almost coinciding with the solid curves, represent the corresponding deterministic profiles (response to periodic spike train inputs with frequency *f*_*spk*_). **A1**. *τ*_*dep*_ = *τ*_*fac*_ = 0.1. **A2**. *τ*_*dep*_ = *τ*_*fac*_ = 100. **A3**. *τ*_*dep*_ = *τ*_*fac*_ = 500. **A4**. *τ*_*dep*_ = *τ*_*fac*_ = 1000. **B**. Averaged variances for the *V*_*peak*_ (solid) and *V*_*trough*_ (dashed) profiles as a function of *τ*_*dep*_ = *τ*_*fac*_ for representative parameter values. The average variance for each profile was computed by averaging the corresponding response variances across all values of *< f*_*spk*_ *>*. **B**. Increasing the input variability (*δ*) increases the response variability. Panels B1 and B2 show two realizations for the same parameter values. We used *τ* = 10 and *τ*_*dec*_ = 10. **C**. Decreasing the synaptic decay time *τ*_*dec*_ causes an increase in the response variability. We used *τ* = 10 and *δ* = 0.1. **D**. Increasing the membrane potential time constant causes an increase in the response variability. We used *τ*_*dec*_ = 10 and *δ* = 0.1. We used the following additional parameter values: *C* = 1, *E*_*L*_ = −60, *I*_*app*_ = 0, *E*_*syn*_ = 0, *a*_*d*_ = 0.1, *a*_*f*_ = 0.1, *x*_∞_ = 1, *z*_∞_ = 0 and *T*_*sw*_ = 1.

#### 3.10.2 PSP *V*_*peak*_ and Γ_*V*_ BPFs persist in response to Poisson-distributed spike trains and are modulated by these inputs

We use Poisson-distributed spike trains with stationary mean firing rates *r*_*spk*_ (*< f*_*spk*_ *>*) within the same range as the spiking input frequencies used above. For comparison with the previously discussed cases, we identify the mean firing rate with the spiking frequency *f*_*spk*_ from which it originates.

Fig. 15-A illustrates that PSP *V*_*peak*_ and Γ_*V*_ filters persists in response to Poisson-distributed presynaptic inputs, but the perturbations from the classical filters (in response to periodic inputs) are more prominent than in the case discussed above (the solid and dashed curves do not coincide and they are significantly further apart). As expected, the response variability to Poisson inputs is larger than for jittered-periodic inputs (compare the shadow regions in Figs. 14-A and 15-A and Figs. S16-A and S17-A). Similarly to the filters discussed above, the variability is larger for the *V*_*peak*_ and *V*_*trough*_ filters than for the other filters, is frequency-dependent (compare the shadow regions for each filter across frequencies), and STP-dependent (compare the shadow regions for each input frequency across values of *τ*_*dep*_ and *τ*_*fac*_.

**Figure 15.**
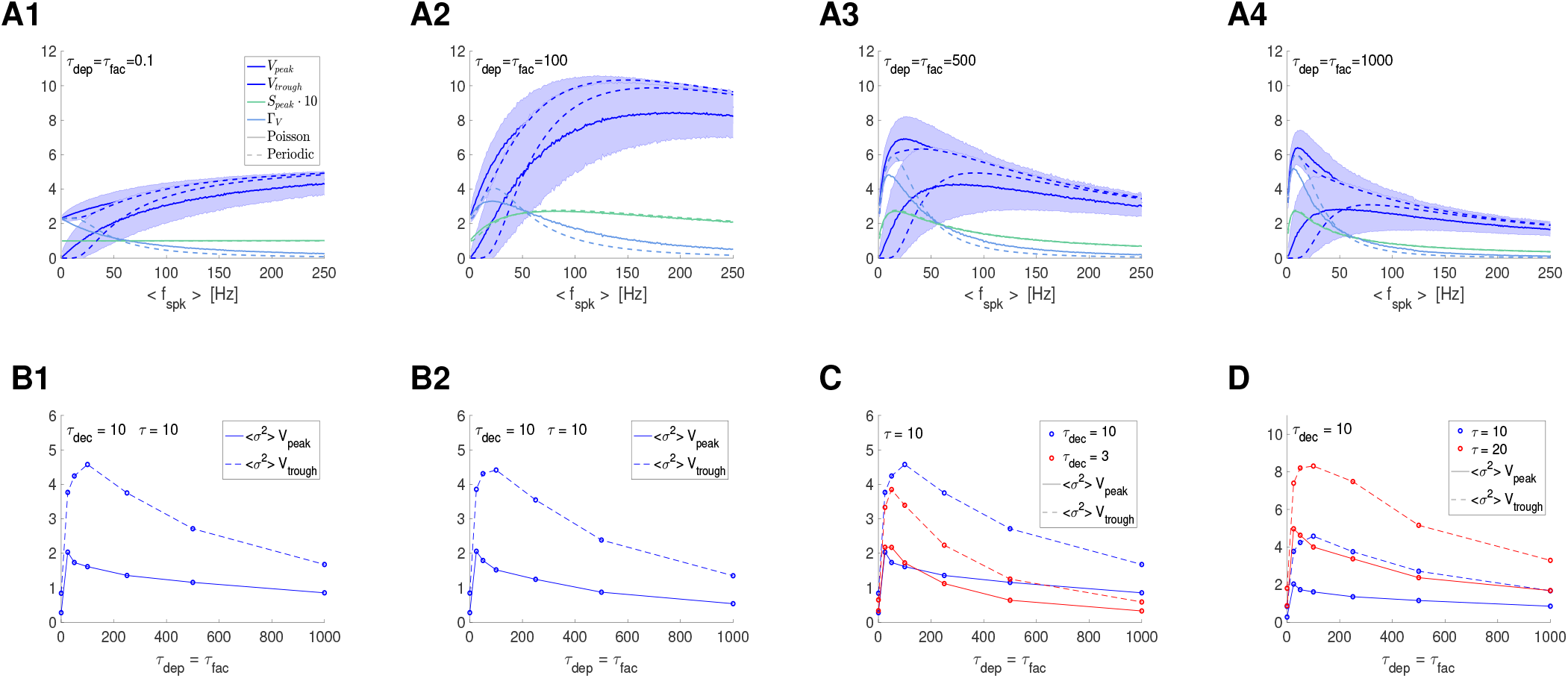
Postsynaptic filters in response to Poisson-distributed presynaptic inputs in the presence of STP: frequency- and STP-dependent variability. The mean rate of the Poisson distributed spike trains corresponds to *< f*_*spk*_ *>*. **A**. Superimposed *V*_*peak*_, *V*_*trough*_, *S*_*peak*_ and Γ_*V*_ profiles for representative parameter values. We used *τ*_*dec*_ = 10 and *τ* = 10 in all panels. Solid curves correspond to the mean values for each attribute (*V*_*peak*_, *V*_*trough*_, *S*_*peak*_ and Γ_*V*_). The shadow regions (*V*_*peak*_, *V*_*trough*_ and *S*_*peak*_ correspond to one standard deviation from the mean. The dashed curves represent the corresponding deterministic profiles (response to periodic spike train inputs with frequency *f*_*spk*_). **A1**. *τ*_*dep*_ = *τ*_*fac*_ = 0.1. **A2**. *τ*_*dep*_ = *τ*_*fac*_ = 100. **A3**. *τ*_*dep*_ = *τ*_*fac*_ = 500. **A4**. *τ*_*dep*_ = *τ*_*fac*_ = 1000. **B**. Averaged variances for the *V*_*peak*_ (solid) and *V*_*trough*_ (dashed) profiles as a function of *τ*_*dep*_ = *τ*_*fac*_ for representative parameter values. The average variance for each profile was computed by averaging the corresponding response variances across all values of *< f*_*spk*_ *>*. **B**. Two realizations for the same parameter values. We used *τ* = 10 and *τ*_*dec*_ = 10. **C**. Decreasing the synaptic decay time *τ*_*dec*_ causes a decrease in the response variability for large values of *< f*_*spk*_ *>* and an increase in the response variability for (very) small values of *f*_*spk*_. We used *τ* = 10 and *δ* = 0.1. **D**. Increasing the membrane potential time constant causes an increase in the response variability. We used *τ*_*dec*_ = 10 and *δ* = 0.1. We used the following additional parameter values: *C* = 1, *E*_*L*_ = −60, *I*_*app*_ = 0, *E*_*syn*_ = 0, *a*_*d*_ = 0.1, *a*_*f*_ = 0.1, *x*_∞_ = 1, *z*_∞_ = 0 and *T*_*sw*_ = 1.

### 3.11 STP controls the variability of PSP *V*_*peak*_ and Γ_*V*_ BPFs in response to jittered-periodic and Poisson distributed spike trains

A salient feature of the frequency filters presented in Figs. 14-A and 15-A is the dependence of the response variability (shadow regions), not only with the mean input frequency (or rate), but also with the time constants controlling the short-term depression and facilitation processes (for the same input frequencies).

We quantified these dependencies in Figs. 14- and 15-B, -C and D for values of *τ*_*dep*_ = *τ*_*fac*_ within a representative range for the *V*_*peak*_ and *V*_*trough*_ profiles. For each value of *τ*_*dep*_ = *τ*_*fac*_ we computed the average variance for each profile (across input frequencies). We obtained similar results by using the maximal variance for each profile.

In all cases, the variability first increases as *τ*_*dep*_ = *τ*_*fac*_ increase for relative small values of these parameters, and then it decreases as *τ*_*dep*_ = *τ*_*fac*_ continues to increase. For each set of time constants (*τ*_*dec*_ and *τ*), there is a value of *τ*_*dep*_ = *τ*_*fac*_ for which the variability is maximal. As expected, for the jittered periodic inputs the profiles’ variability increases as the input variability increases (Figs. 14-B). The profiles’ variability also increases as *τ*_*dec*_ decreases (Figs. 14-C) and *τ* increases (Figs. 14-D). For the Poisson-distributed inputs, in contrast, the profiles’ variability decreases with decreasing values of *τ*_*dec*_ (Fig. 15-C), while it increases for increasing values of *τ* (Fig. 15-D).

Together, these results and the results of the previous section show that STP plays important roles not only in determining the patterns exhibit by networks, but also their robustness and accuracy of the information transmission.

## 4 Discussion

Neuronal filters play important roles in neuronal communication, information selection and processing, neuronal oscillations, neuronal resonance and brain computations [1–18]. Neuronal filters emerge as the result of a variety of mechanisms that involve a multiplicity of time scales associated to the neuronal electric circuit and biophysical properties and history-dependent processes. In spite of it ubiquitousness, the mechanisms of generation of neuronal filters are poorly understood beyond the single cell level. In previous work [4], we showed that band-pass filters (BPFs) can be inherited across levels of neuronal organization or can be created independently at various levels of organization by the interplay of low-pass filters (LPFs) and high-pass filters (HPFs) belonging to the same or different levels (e.g., schematic diagrams in Fig. 2). However, the mechanism by which these BPFs, LPFs and HPFs are generated, interact and are modulated as they transition across levels of organization are not clear.

We set out to address these issues in an elementary network motif (Fig. 1). We investigated the biophysical and dynamic mechanisms of generation of postsynaptic (PSP) filters in response to presynaptic spike inputs in the presence of synaptic short-term plasticity (STP). We were particularly interested in understanding how PSP filters are shaped by the time scales associated with the participating building blocks: the synaptic rise and decay dynamics (*τ*_*rse*_ and *τ*_*dec*_), the synaptic STP time constants (*τ*_*dep*_ and *τ*_*fac*_), and the intrinsic time constant (*τ*) of the postsynaptic cells (Fig. 1). To this end, we conducted a systematic study of the steady-state PSP responses to (i) periodic presynaptic inputs over a range of frequencies (*f*_*spk*_ = 1000*/*Δ_*spk*_; Fig. 1-A, top), (ii) jittered-periodic inputs over a range of mean frequencies (*f*_*spk*_ = 1000*/*Δ_*spk*_), and (iii) Poisson-distributed presynaptic inputs over a range of mean rates (*r*_*spk*_ = 1000*/ <* Δ_*spk*_ *>*; Fig. 1-A, bottom).

The use of periodic presynaptic spike trains allowed us to systematically understand the basic mechanistic aspects governing the generation of frequency filters in the elementary network motif: (i) how frequency-filters at the different levels of organization are shaped by the time scales of the participating building blocks, (ii) how the filtering properties are communicated across levels of neuronal organization, and (iii) how filters interact across levels of neuronal organization. The use of Poisson-distributed presynaptic spikes trains allowed us to extend our investigation and findings to more realistic scenarios and to explore how the variability of the filtering properties is controlled by the biophysical properties and time scales of the participating building blocks, primarily STP. The jittered-periodic inputs were used as an intermediate step to link between the purely deterministic and stochastic spike-train inputs.

We used a combination of mathematical modeling, analytical calculations and numerical simulations. As part of our study, we developed reduced models and analytical approximations of the membrane potential response of passive cells to presynaptic spike trains in the presence of STP. This allowed us to compute the peak (*V*_*peak*_) and peak-to-trough amplitude (Γ_*V*_) filters in terms of the participating time constants and other model parameters. While we could have conducted this study by using solely numerical simulations, obtaining analytical expressions provided us with a better understanding of the dependence of the PSP filters on the model parameters and time scales. This was achieved at the expense of some simplifying assumptions that transform the multiplicative synaptic input to the passive membrane equation into an sequentially-corrected additive input where the additive synaptic input was corrected at the arrival of each presynaptic spike to account for the membrane potential changes during the presynaptic interspike interval. The resulting expressions make the contribution of the time constants and other parameters to the PSP filters’s shape apparent. One could argue that the sequentially-corrected reduced model was not necessary since the passive membrane equation is linear, and the response to multiplicative inputs is analytically solvable (e.g., by using Laplace transforms). However, the complex formula describing the exact analytical solution fails to provide the level of clarity and intuition desired to understand how the model parameters shape the PSP filters. While we consider here the sequentially-corrected reduced model, eqs. (25)-(27), as an approximation to the more detailed biophysical model, we note that it could stand as a model of their own [100]. The approach we used in this paper can be extended to more complex scenarios such as networks where the presynaptic or postsynaptic cells are described by more complex linearized models involving additional ionic currents and weakly nonlinear models [101].

The role of STP on information filtering and related phenomena has been investigated before by many authors [3, 9–11, 17, 24, 26, 28–45, 48–53]. However, previous studies have not focused on the mechanisms of generation of PSP frequency-filters and how they are shaped by the participating time scales. Previous work has also ignored the mechanisms governing the variability of the response to realistic presynaptic spike-train inputs. Because the synaptic response *S* is not directly measurable, a common simplifying assumption has been made by some authors [30, 48, 49, 51]: that the voltage response of the postsynaptic cell is a scaled version of the synaptic response. While this assumption might be justified in many cases, our results indicate that it is by no mean universally expected, and there could be significant qualitative differences between the synaptic and postsynaptic frequency-filters. This was also highlighted in previous work on temporal filters in the presence of STP [46].

We divided our study in three steps. We investigated the response profiles (frequency-filters) of (i) the synaptic update Δ*S* to the presynaptic spike trains (the target of the synaptic variable *S*), (ii) the synaptic variable *S* through the synaptic update Δ*S*, and (iii) the postsynaptic membrane potential *V* to the synaptic variable *S*. We characterized the frequency-filters by using the (stationary) peak profiles (for 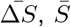 and 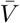) and the (stationary) peak-to-trough amplitude profiles (Γ_*S*_ and Γ_*V*_). By design, the effects of STP are present at the Δ*S* level giving rise to the synaptic update sequences Δ*S*_*n*_ = *X*_*n*_*Z*_*n*_, which depend on the time constants *τ*_*dep*_ and *τ*_*fac*_ and on *f*_*spk*_ (the 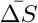 filter is steady-state of these sequences). These are the target of the synaptic variables *S* during the rise phase immediately after the arrival of each presynaptic spike. The *S* frequency-filters are shaped by Δ*S*_*n*_ = *X*_*n*_*Z*_*n*_ and the synaptic time constantes *τ*_*rse*_ and *τ*_*dec*_. In turn, the synaptic variable *S* is the input to the current-balance equation describing the dynamics of the passive cell. The PSP frequency-filters result from the interaction between *S* and the biophysical properties of the postsynaptic passive cell, particularly the time constant *τ* = *C/G*_*L*_.

Consistently with previous work, [3, 23, 30, 48, 49, 51], 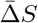 BPFs are generated by the interplay of low-pass (depression) and high-pass (facilitation) filters for the appropriate balances between the two processes (e.g., Fig. 2-A). They are amplified by increasing values of *τ*_*fac*_ and the 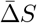 resonant frequency decreases as *τ*_*fac*_ increases. The 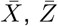 and 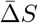 profiles develop in response to multiple events, each controlled by the time constants *τ*_*dep*_ and *τ*_*fac*_. We described the (global in time) filter properties in terms of the characteristic frequencies *σ*_*dep*_ 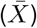, *σ*_*fac*_ 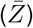, and the characteristic frequency difference Δ_*kappa*_ = *κ*_*rse*_ and *κ*_*dec*_ 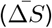. These depend on *τ*_*dep*_ and *τ*_*fac*_ in a nonlinear manner. Increasing values of *τ*_*fac*_ and *τ*_*dep*_ cause *σ*_*fac*_ and *σ*_*dep*_, respectively, to decrease (sharpen the 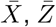 filters, respectively). Increasing values of *τ*_*fac*_ cause the 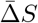 resonant frequency to decrease, the 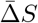 peak to increase and the 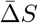 BPF to become sharper. Increasing values of *τ*_*dep*_ cause the 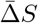 resonant frequency to decrease and the 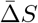 BPF to become sharper, but cause the the 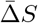 peak to decrease.

For the to-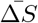 update model with instantaneous *S* rise, the peak envelope profiles 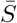 are identical to the 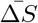 profiles. The Γ_*S*_ BPFs can be inherited from 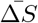 ones or can be created by the interplay of a 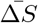 HPF and *Q*_*A*_ (a LPF). In all cases, they become sharper and less peakier as *τ*_*dec*_ increases. For the to-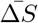 update model with non instantaneous *S* rise, the 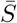 profiles are attenuated as *τ*_*rse*_ increases and the Γ_*S*_ profiles are also attenuated as *τ*_*rse*_ increases.

For the by-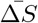 update model with instantaneous *S* rise, in contrast to the to-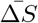 model, the Γ_*S*_ profiles are inherited from the 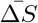 profiles. The 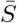 profiles transition from LPFs or BPFs to HPFs as *τ*_*dec*_ increases, which may be bounded or unbounded (if they are HPFs, they remain so). These models lack a biophysical mechanism that balances the summation effects and created realistically saturated profiles. For the by-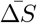 update model with non-instantaneous *S* rise, the 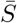 profiles are attenuated as *τ*_*rse*_ increases and the Γ_*S*_ profiles are also attenuated as *τ*_*rse*_ increases.

For linear systems in response to sinusoidal inputs, amplitude and membrane potential peak filters coincide. This is no longer true for nonlinear systems or neuronal systems receiving presynaptic inputs. Therefore, we investigated the two types of filters (*V*_*peak*_ and Γ_*V*_). Passive cells exhibit LPFs in response to sinusoidal input currents. In contrast, they exhibit *V*_*peak*_ HPFs in response to periodic presynaptic inputs in the absence of STP, while still exhibiting Γ_*V*_ LPFs. These filters are modulated by the synaptic time constant *τ*_*dec*_ and the postsynaptic time constant *τ*. In the presence of STP, *V*_*peak*_ and Γ_*V*_ BPFs are possible under certain conditions. We found two qualitatively different mechanisms. The *V*_*peak*_ and Γ_*V*_ BPFs can be either inherited from the synaptic level, subject to the appropriate modulations, or generated across levels. Specifically, a *V*_*peak*_ BPF can be either result of an *S*_*peak*_ BPF generated as the result of synaptic depression and facilitation or the result of a *S*_*peak*_ LPF generated by the interplay of synaptic depression and PSP summation. Similarly, a Γ_*V*_ BPF can be the result of the interplay of synaptic facilitation and PSP summation.

These types of BPFs persist in response to jitter periodic spike trains and Poisson-distributed spike trains, exhibiting a frequency-dependent variability. Importantly, the variability properties of these BPFs are controlled by STP in a frequency-dependent manner. This highlights a role of STP as regulating the variability of PSP filters and creating variability filters. To our knowledge, this has not been described before and requires a systematic study.

The results of this study complement previous work [46] where we thoroughly investigated the temporal (transient) responses and the associated temporal filters in the same feedforward network motif (Fig. 1). One important result of this study was the description of the link between the STP time constants governing the dynamics of the single events (in response to each presynaptic spikes; *τ*_*dep*_ and *τ*_*fac*_) and the corresponding global, emergent time scales describing the long-term dynamics of the (low-, high- and band-pass) temporal filters. A second important result was the discovery of a third global time scale for temporal band-pass filters involving a combination of both *τ*_*dep*_ and *τ*_*fac*_, highlighting the complexity of the non-trivial interaction between depression and facilitation. A third important result was the finding that the postsynaptic temporal filters are not proportional to the synaptic temporal filters as assumed in some simplified models, [48], thus demonstrating that the synaptic temporal filters are not directly communicated to the postsynaptic cell level, but rather modified by the postsynaptic intrinsic properties. However, temporal filters are not necessarily predictive of frequency-filters and both types of filters are generated by different mechanisms.

The feedforward network motif we studied (Fig. 1) is arguably the basic network processing unit that can produce STP-dependent PSP BPFs. This allow us to systematically investigate the qualitatively different types of mechanisms of generation of PSP BPFs according to whether they are inherited from the synaptic level or generated across levels of organization. These mechanisms are expected to operate in larger and more complex networks of which the basic network motif we studied here becomes a building block. Testing these ideas requires extending our work to include presynaptic BPFs and presynaptic patterns generated by oscillatory input currents to the presynaptic cell. It would be natural to hypothesize the existence of inherited PSP BPFs in the absence of STP and PSP BPFs generated by cross-level mechanisms including the presynaptic level. Further research is needed to understand these scenarios and to understand more complex network motifs including additional connections (e.g., feedback connections both excitatory and inhibitory) in the presence of STP in one or all of them and combinations of the network motif studied here. Additional research should focus on the modulation of STP-dependent PSP filters by astrocytes, which are known to regulate synaptic depression and facilitation [102]. Additional research is also needed to understand the mechanisms of regulation of PSP filters’ variability by STP. Finally, future work should focus on more complex scenarios and more complex biophysically realistic models of STP where STP is determined by the combined effect of both pre- and post-synaptic factors [103] (see also [47, 48, 104]). Complex scenarios should include the effects of background noise, neuronal noise and spiking irregularities [15, 105–108].

In this context, our study is a first step in the systematic understanding of the mechanisms of generation of STP-mediated neuronal filters in networks and contributes to the systematic understanding of information processing via neuronal filters in neuronal systems.

## Acknowledgments

HGR acknowledges support from the National Science Foundation grants DMS-1608077 and IOS-2002863. YM acknowledges support from the National Science Foundation Graduate Research Fellowship. The authors are grateful to Allen Tannenbaum for useful comments and support, and to Farzan Nadim, Dirk Bucher, Nelly Daur, Antoni Guillamon, Gema Huguet and Tomas Lázaro for useful comments and discussions. HGR is a Corresponding Investigator in CONICET, Argentina and a Graduate Faculty Member in the Behavioral Neurosciences Program at Rutgers University. HGR is also Visiting Researcher/Academic in the Neurosciences Institute and the Courant Institute of Mathematical Sciences at New York University.

## A Approximate analytical solution to the passive postsynaptic cell receiving presynaptic spike inputs

Here we compute an analytical approximate solution to the model (1)-(4), describing the dynamics of a postsynaptic passive cell receiving presynaptic spike inputs at time *t*_1_, *t*_2_, ….

### A.1 Approximate solution to the synaptic variable

We first compute the approximate solution to the synaptic variable *S* whose dynamics are described by eq. (3) with *S*(0) = 0, under the assumption of instantaneous rise to a value Δ*S*_*n*_ (*n* = 1, 2 …) at the arrival of each presynaptic spike. The assumption *S*(0) = 0 implies that *S*(*t*) = 0 for 0 ≤ *t* ≤ *t*_1_. For the duration of a presynaptic spike (*t*_*n*_ *< t < t*_*n*_ + *T*_*sw*_), we approximate *S*(*t*) by the synaptic update value Δ*S*_*n*_ (constant). For the remainder of the presynaptic interspike interval (*t*_*n*_ + *T*_*sw*_ ≤ *t < t*_*n*+1_), *S*(*t*) decreases according to the second term in (3). The approximate solution is given by

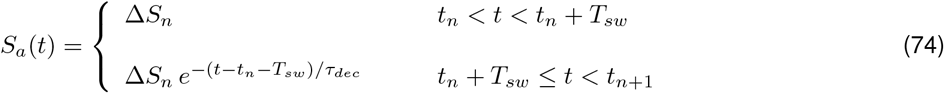

for *n* = 1, 2, ….

From eq. (74) for *S*_*a*_(*t*), the larger the duration *T*_*sw*_ of the presynaptic spike (*T*_*sw*_ *<* Δ_*spk*_) or the larger the synaptic decay time *τ*_*dec*_, the larger *S*_*a*_(*t*) during the presynaptic ISI. In that sense, increasing *τ*_*dec*_ or *T*_*sw*_ has a similar overall effect on the PSP response as increasing *G*_*syn*_. However, changes in the former two have stronger dynamic effects (e.g., cause changes in the voltage response peak) than changes in *G*_*syn*_ (Fig. 8).

### A.2 Approximate solution to the postsynaptic membrane potential: Postsynaptic passive cell

We first approximate eq. (1) as follows

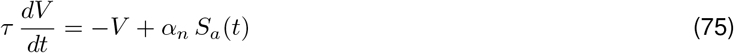

for *n* = 1, …, *N* where

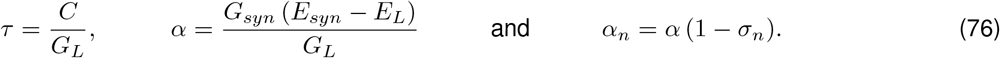

The variable *V* in eq. (75) represents *V* − *E*_*L*_ − *I*_*app*_*/G*_*L*_ in eq. (1). The third term in eq. (75) is a current input approximation to the conductance input in the synaptic current (1). To account for this, we introduce a correction factor 1 − *σ*_*n*_ where *σ*_*n*_ is updated at the beginning of each ISI and *σ*_1_ = 0.

Prior to the arrival of the first presynaptic spike (*t* ≤ *t*_1_), *V* (*t*) = 0. For the duration of a presynaptic spike (*t*_*n*_ *< t < t*_*n*_ + *T*_*sw*_),

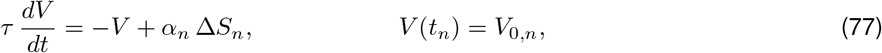

where *V*_0,*n*_ is the value of *V* at the end of the previous cycle. The solution to (77) is given by

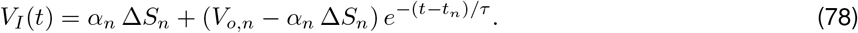

We define

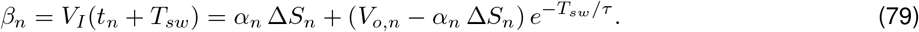

For the remainder of the presynaptic interspike interval (*t*_*n*_ + *T*_*sw*_ *< t < t*_*n*+1_), *dt*

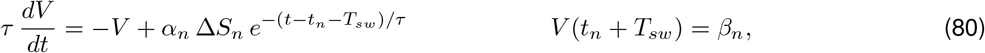

For *τ*_*dec*_ */*= *τ*, the solution to (80) is given by

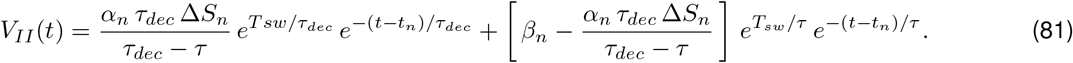

We define

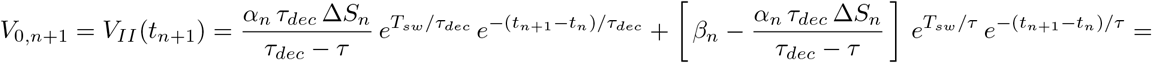

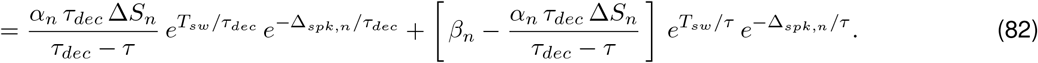

For the duration of the presynaptic spike, *V* (*t*) = *V*_*I*_ (*t*) is an increasing function. *V* (*t*) = *V*_*II*_ (*t*) continues to be an increasing function after the presynaptic spike is off until the two exponential terms balance out and *V*_*II*_ (*t*) decreases. Therefore, within some frequency range, *V* (*t*) peaks after the presynaptic spike is off. The peak time for *V*_*II*_ (*t*) in eq. (81) is given

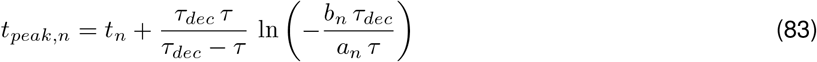

where

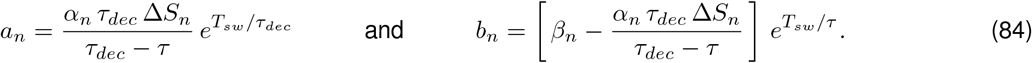

The peak value of *V* (*t*) is given by

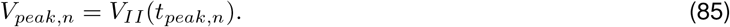

We define *σ*_*n*+1_ as

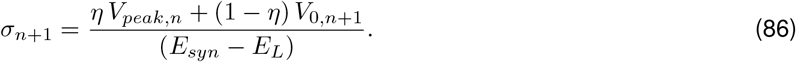

Or

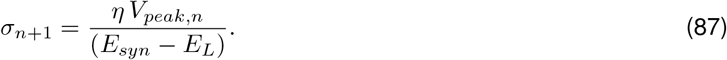

For *τ*_*dec*_ = *τ*, the solution to (80) is given by

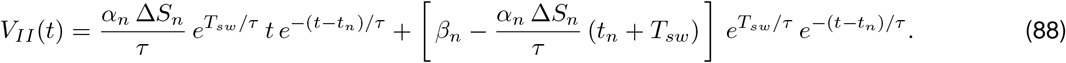

The peak time for *V*_*II*_ (*t*) in eq. (88) is given

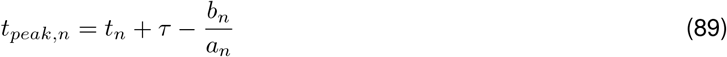

Where

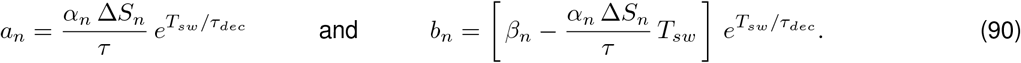

The peak values of *V* (*t*) is given by (85) with *β*_*n*_ given by (79) and *V*_0,*n*+1_ = *V*_*II*_ (*t*_*n*+1_)

## Supplementary Material

**Figure S1:**
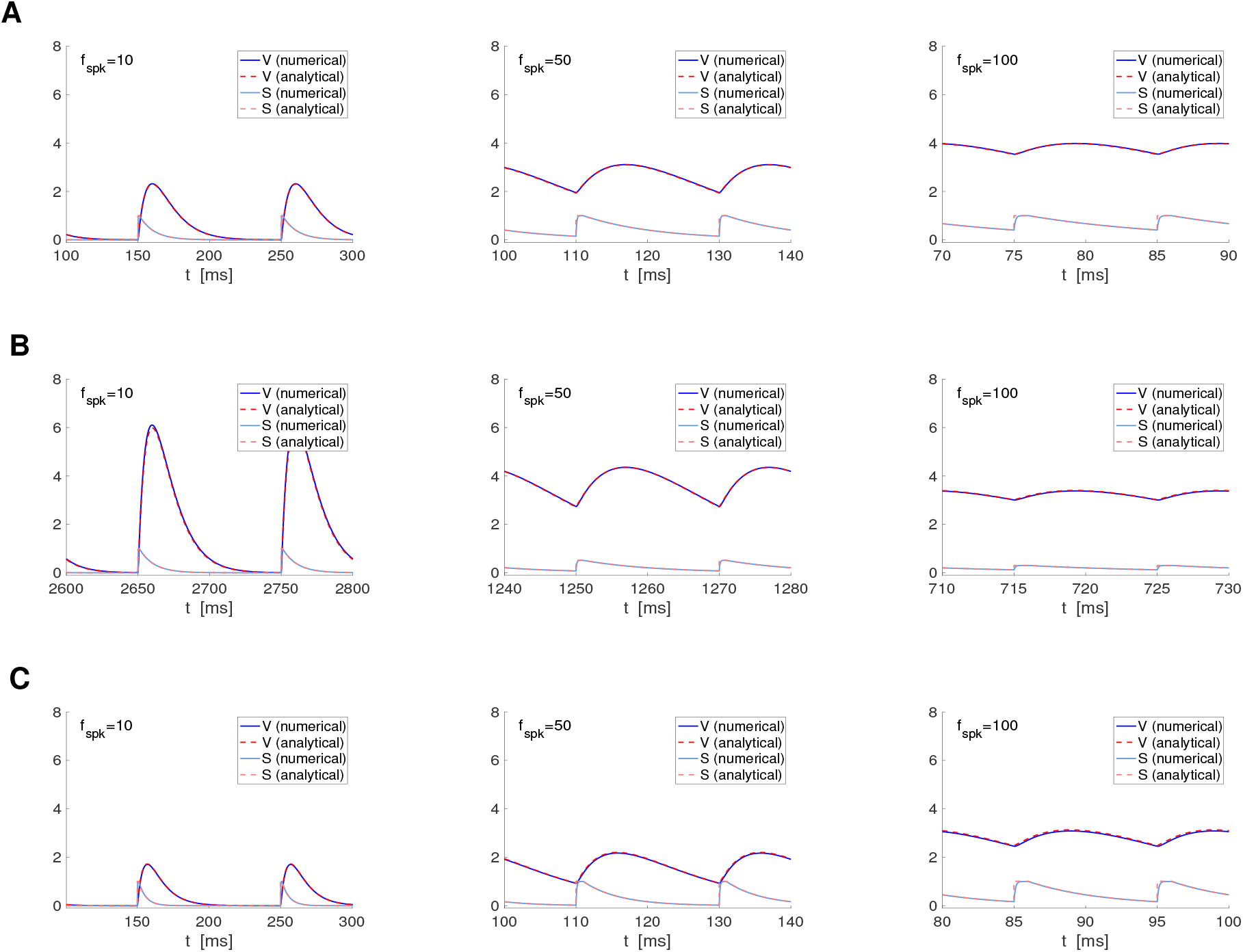
Analytical approximation of the membrane potential response of passive cells to presynaptic spikes: Representative examples I. For the numerical approximations we used the model for a passive cell receiving presynaptic spike-train input (1)-(4). For STP we use the DA model (7)-(9). For the analytical approximations we used eqs. (25)-(27) together with eqs. (78) and (81) in the Appendix A. **A**. *G*_*L*_ = 0.1 (*τ* = 10), *τ*_*dec*_ = 10, *τ*_*dep*_ = *τ*_*fac*_ = 0.1. **B**. *G*_*L*_ = 0.1 (*τ* = 10), *τ*_*dec*_ = 10, *τ*_*dep*_ = *τ*_*fac*_ = 1000. **C**. *G*_*L*_ = 0.1 (*τ* = 10), *τ*_*dec*_ = 5, *τ*_*dep*_ = *τ*_*fac*_ = 0.1. We used the following additional parameter values: *a*_*d*_ = 0.1, *a*_*f*_ = 0.1, *x*_∞_ = 1, *z*_∞_ = 0, *τ*_*rse*_ = 0.1, *C* = 1, *E*_*L*_ = −60, *I*_*app*_ = 0, *G*_*syn*_ = 0.1, *E*_*syn*_ = −60.

**Figure S2:**
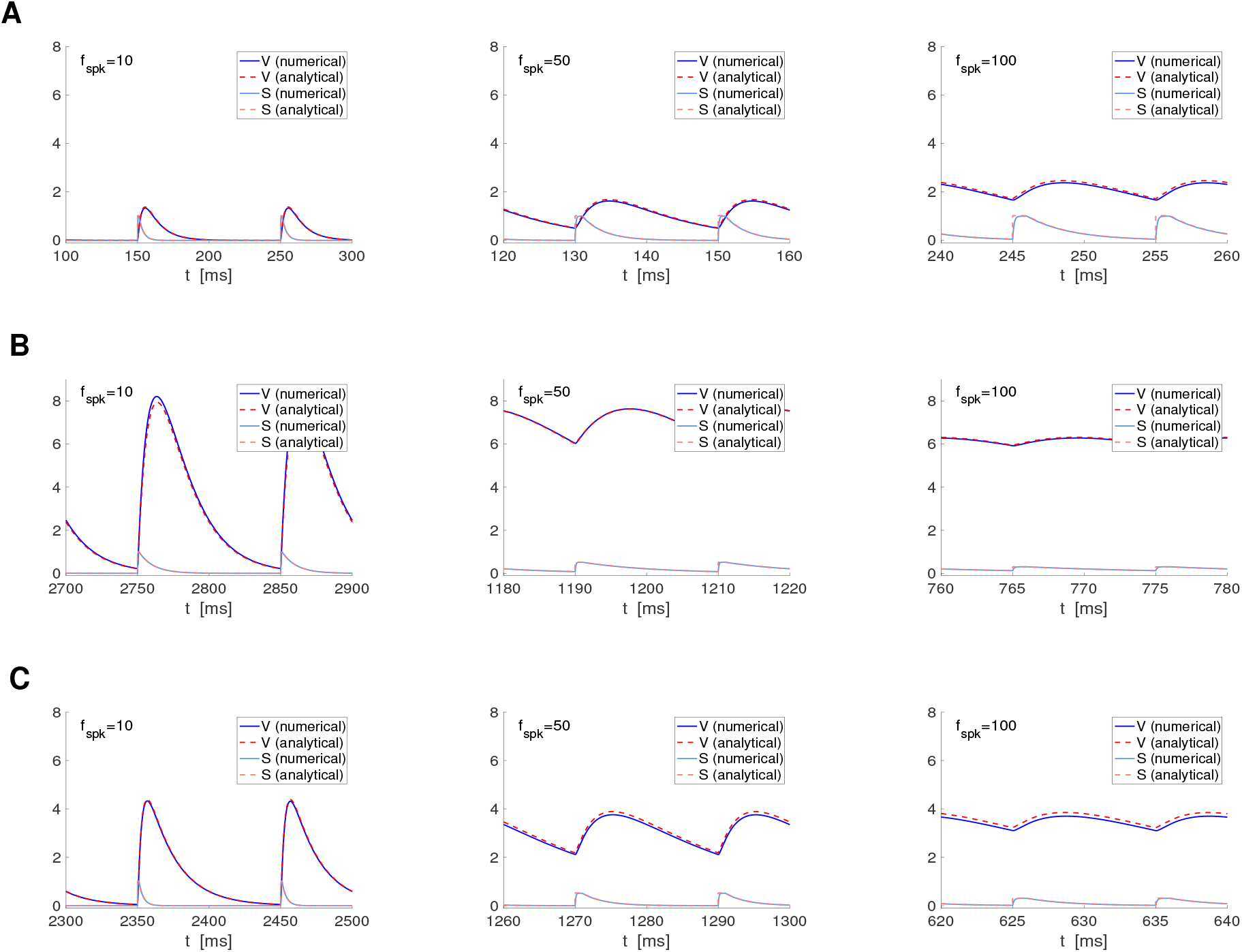
Analytical approximation of the membrane potential response of passive cells to presynaptic spikes: Representative examples II. For the numerical approximations we used the model for a passive cell receiving presynaptic spike-train input (1)-(4). For STP we used the DA model (7)-(9). For the analytical approximations we used eqs. (25)-(27) together with eqs. (78) and (81) in the Appendix A. **A**. *G*_*L*_ = 0.1 (*τ* = 10), *τ*_*dec*_ = 3, *τ*_*dep*_ = *τ*_*fac*_ = 0.1. **B**. *G*_*L*_ = 0.05 (*τ* = 20), *τ*_*dec*_ = 10, *τ*_*dep*_ = *τ*_*fac*_ = 1000. **C**. *G*_*L*_ = 0.05 (*τ* = 20), *τ*_*dec*_ = 5, *τ*_*dep*_ = *τ*_*fac*_ = 1000. We used the following additional parameter values: *a*_*d*_ = 0.1, *a*_*f*_ = 0.1, *x*_∞_ = 1, *z*_∞_ = 0, *τ*_*rse*_ = 0.1, *C* = 1, *E*_*L*_ = −60, *I*_*app*_ = 0, *G*_*syn*_ = 0.1, *E*_*syn*_ = −60.

**Figure S3:**
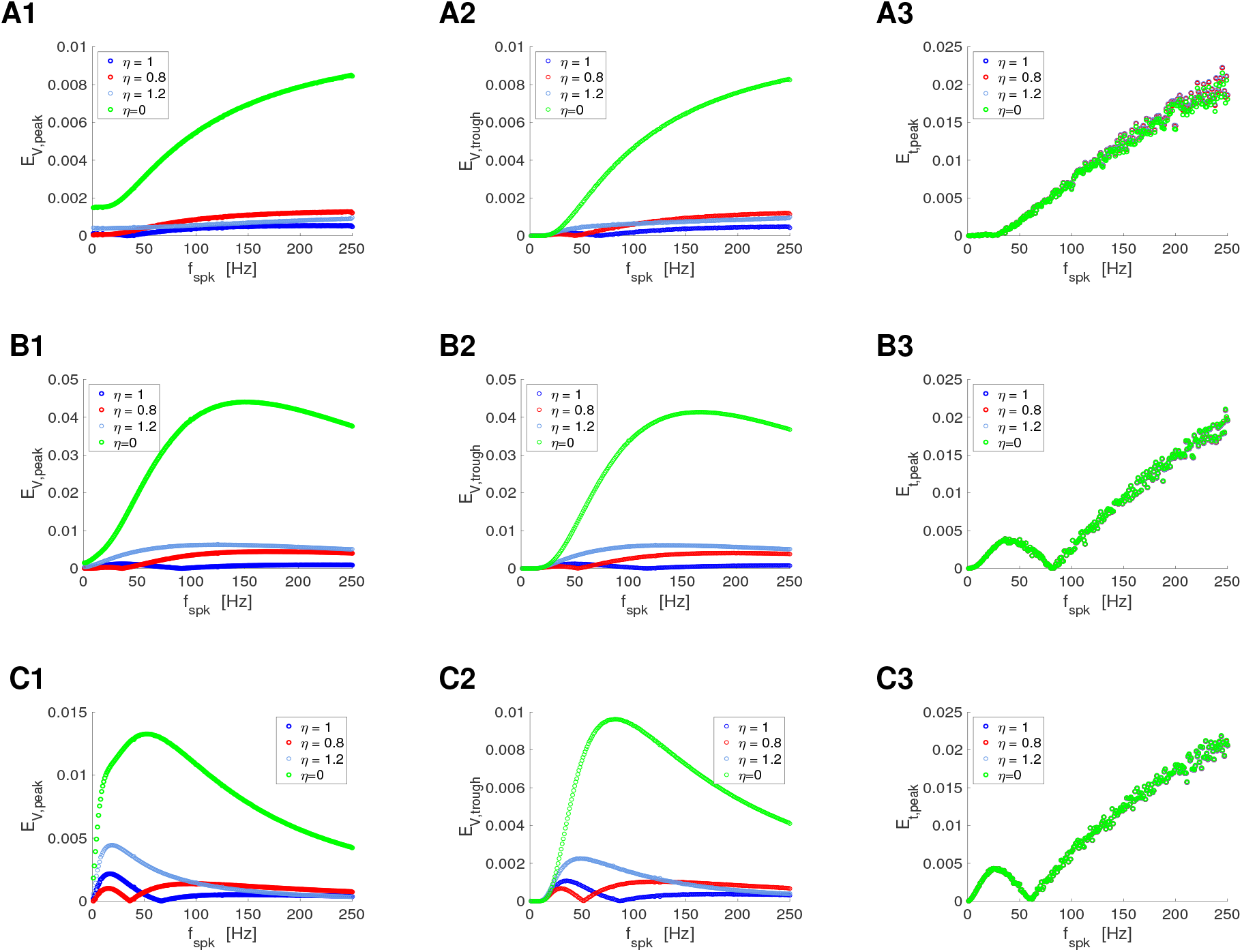
Error between the numerical and analytical approximations for the stationary peaks (V_*peak*_), troughs (V_*trough*_), and peak times (t_*peak*_) of the membrane potential responses of passive cells to presynaptic spikes: Representative examples I. For the numerical (num) approximations we used the model for a passive cell receiving presynaptic spike-train input (1)-(4). For STP we used the DA model (7)-(9). For the analytical approximations we used eqs. (25)-(27) together with eqs. (78) and (81) in the Appendix A. For the computations of the analytical (anl) approximations to *V*_*peak*_, *V*_*trough*_ and *t*_*peak*_ we used eqs. (28)-(35). Simulations were carried out until the difference between two consecutive numerical peaks were below a tolerance value equal to 0.001. The last values of *V*_*peak*_, *V*_*trough*_ and *t*_*peak*_ in the resulting sequences were taken as an approximation to the corresponding stationary values. **Left column**. Relative error for *V*_*peak*_ defined as | *V*_*peak,num*_ − 60 − *V*_*peak,anl*_ |*/*| *V*_*peak,num*_ |. **Middle column**. Relative error for *V*_*trough*_ defined as | *V*_*trough,num*_ −60−*V*_*trough,anl*_ |*/*| *V*_*trough,num*_ |. **Right column**. Relative error for *t*_*peak*_ defined as | *t*_*peak,num*_ −*t*_*peak,anl*_ |*/*| Δ_*spk*_ |. **A**. *G*_*L*_ = 0.1 (*τ* = 10), *τ*_*dec*_ = 10, *τ*_*dep*_ = *τ*_*fac*_ = 0.01. **B**. *G*_*L*_ = 0.1 (*τ* = 10), *τ*_*dec*_ = 5, *τ*_*dep*_ = *τ*_*fac*_ = 100. **C**. *G*_*L*_ = 0.1 (*τ* = 10), *τ*_*dec*_ = 10, *τ*_*dep*_ = *τ*_*fac*_ = 500. We used the following additional parameter values: *a*_*d*_ = 0.1, *a*_*f*_ = 0.1, *x*_∞_ = 1, *z*_∞_ = 0, *τ*_*rse*_ = 0.1, *C* = 1, *E*_*L*_ = −60, *I*_*app*_ = 0, *G*_*syn*_ = 0.1, *E*_*syn*_ = −60, Δ*t* = 0.01.

**Figure S4:**
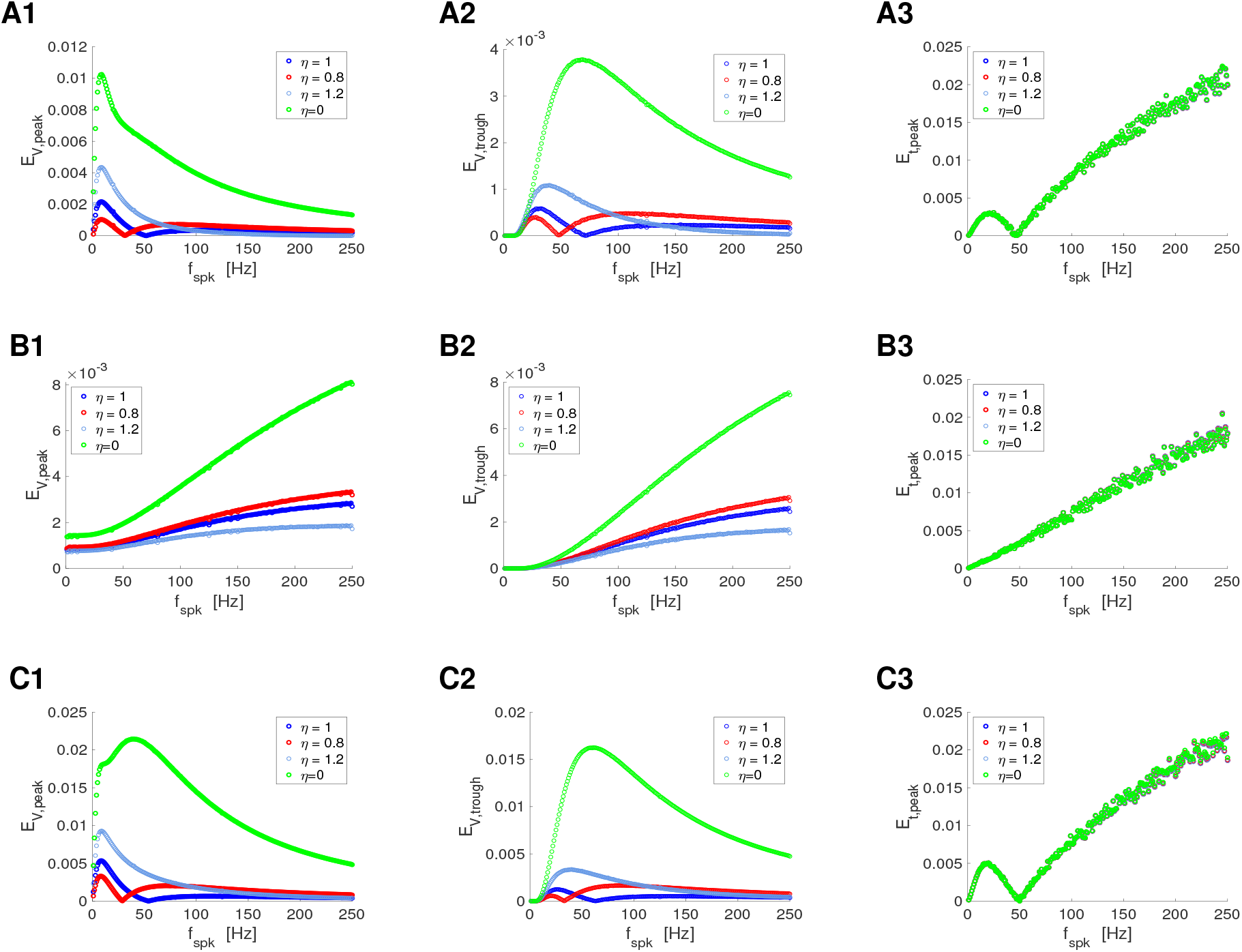
Error between the numerical and analytical approximations for the stationary peaks (V_*peak*_), troughs (V_*trough*_), and peak times (t_*peak*_) of the membrane potential responses of passive cells to presynaptic spikes: Representative examples II. For the numerical (num) approximations we used the model for a passive cell receiving presynaptic spike-train input (1)-(4). For STP we used the DA model (7)-(9). For the analytical approximations we used eqs. (25)-(27) together with eqs. (78) and (81) in the Appendix A. For the computations of the analytical (anl) approximations to *V*_*peak*_, *V*_*trough*_ and *t*_*peak*_ we used eqs. (28)-(35). Simulations were carried out until the difference between two consecutive numerical peaks were below a tolerance value equal to 0.001. The last values of *V*_*peak*_, *V*_*trough*_ and *t*_*peak*_ in the resulting sequences were taken as an approximation to the corresponding stationary values. **Left column**. Relative error for *V*_*peak*_ defined as | *V*_*peak,num*_ − 60 − *V*_*peak,anl*_ |*/*| *V*_*peak,num*_ |. **Middle column**. Relative error for *V*_*trough*_ defined as | *V*_*trough,num*_ −60−*V*_*trough,anl*_ |*/*| *V*_*trough,num*_ |. **Right column**. Relative error for *t*_*peak*_ defined as | *t*_*peak,num*_ − *t*_*peak,anl*_ |*/*| Δ_*spk*_ |. **A**. *G*_*L*_ = 0.1 (*τ* = 10), *τ*_*dec*_ = 10, *τ*_*dep*_ = *τ*_*fac*_ = 1000. **B**. *G*_*L*_ = 0.1 (*τ* = 10), *τ*_*dec*_ = 3, *τ*_*dep*_ = *τ*_*fac*_ = 0.01. **C**. *G*_*L*_ = 0.05 (*τ* = 20), *τ*_*dec*_ = 10, *τ*_*dep*_ = *τ*_*fac*_ = 1000. We used the following additional parameter values: *a*_*d*_ = 0.1, *a*_*f*_ = 0.1, *x*_∞_ = 1, *z*_∞_ = 0, *τ*_*rse*_ = 0.1, *C* = 1, *E*_*L*_ = −60, *I*_*app*_ = 0, *G*_*syn*_ = 0.1, *E*_*syn*_ = −60, Δ*t* = 0.01.

**Figure S5:**
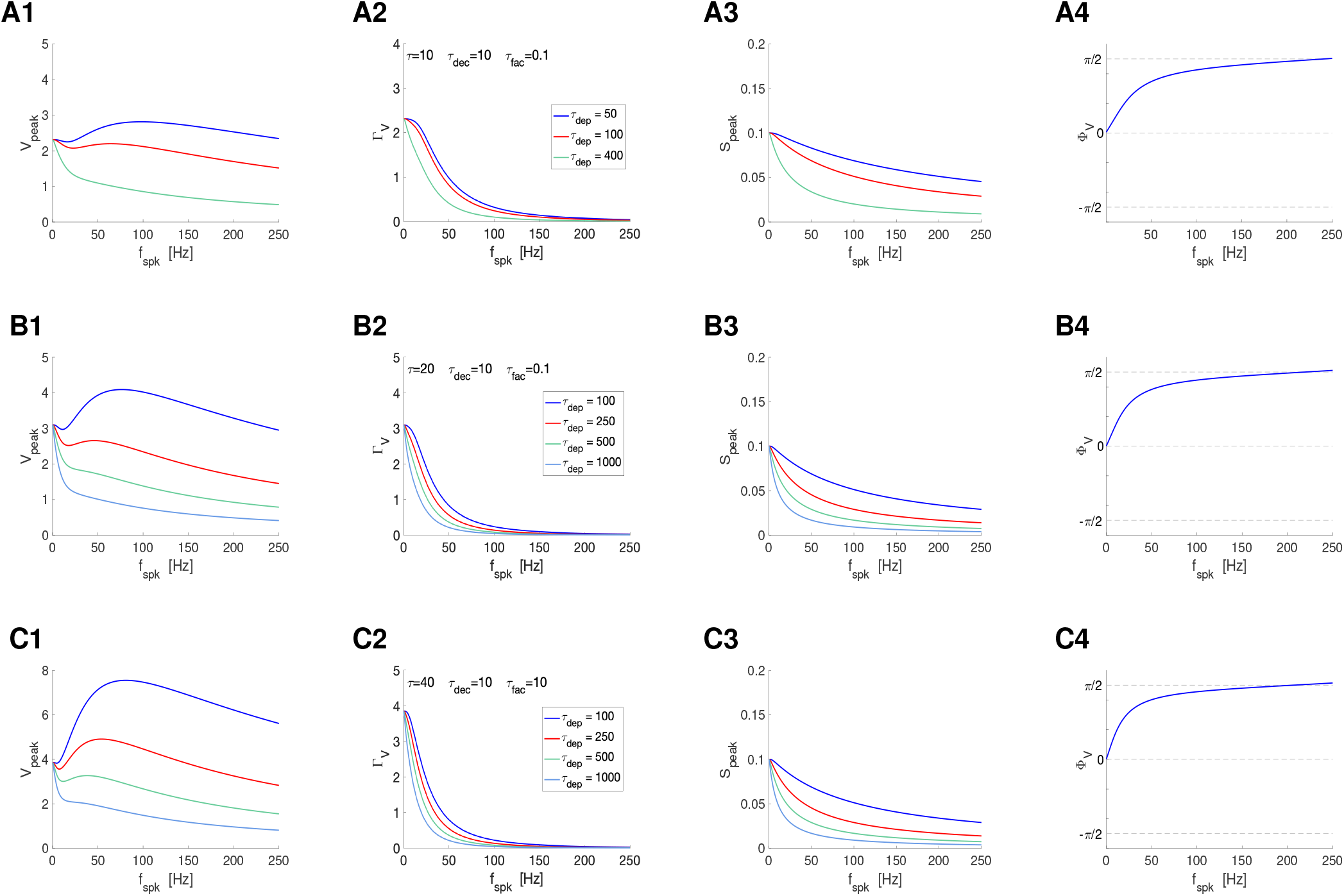
Postsynaptic filters in response to periodic presynaptic spike inputs emerging from the interplay of short-term depression and postsynaptic summation. Superimposed filters for representative values of the short-term depression time constant *τ*_*dep*_. **A**. *τ* = 10. **B**. *τ* = 20. **C**. *τ* = 40. **A, B, C**. *τ*_*dec*_ = 10 and *τ*_*fac*_ = 0.1. **Left column**. *V* peak profiles. **Middle-left column**. *V* peak-to-trough amplitude profiles. **Middle-right column**. *S* peak profiles. **Right column**. *V* phase profiles. We used eq. (65) for the PSP *V* with *I*_*syn*_ described by eqs. (2)-(4) appropriately adapted to account for the translation of *V* to the equilibrium point, and STP described by eqs. (12) and (13) (DA model). The impedance amplitude (*Z*) and phase (Φ_*Z*_) were computed using eqs. (39) and (40). The analytical approximations for the PSP peak sequence response of passive cells to presynaptic inputs are described in Section2.2.4 (see also Appendix A). The approximation of *V*_*peak,n*_, *V*_*trough,n*_ and *t*_*V,peak*_ were computed as described in Section 2.2.4. The PSP amplitude Γ_*V*_ was computed by using eq. (41) and the PSP phase Φ_*V*_ was computed using eq. (42). The synaptic (*S*) peak (*S*_*peak*_) and phase (Φ_*S*_) profiles were computed similarly to these for *V*. We used the following additional parameter values: *C* = 1, *E*_*L*_ = −60, *I*_*app*_ = 0, *G*_*syn*_ = 0.1, *E*_*syn*_ = 0, *a*_*d*_ = 0.1, *a*_*f*_ = 0.1, *x*_∞_ = 1, *z*_∞_ = 0 and *T*_*sw*_ = 1.

**Figure S6:**
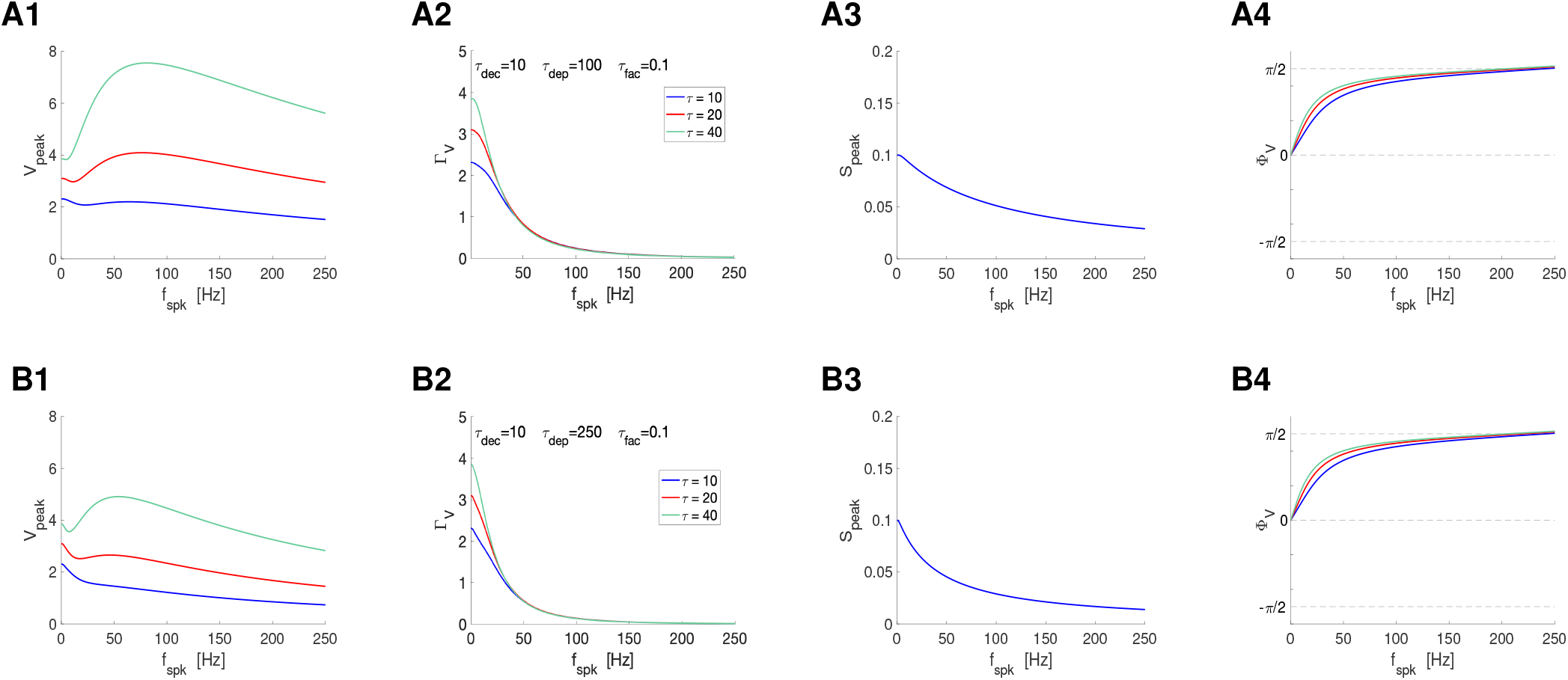
Postsynaptic filters in response to periodic presynaptic spike inputs emerging from the interplay of short-term depression and postsynaptic summation. Superimposed filters for representative values of the membrane time constant *τ*. **A**. *τ*_*dep*_ = 100. **B**. *τ*_*dep*_ = 250. **A, B** *τ*_*dec*_ = 10 and *τ*_*fac*_ = 0.1. **Left column**. *V* peak profiles. **Middle-left column**. *V* peak-to-trough amplitude profiles. **Middle-right column**. *S* peak profiles. They are independent of *τ*. **Right column**. *V* phase profiles. We used eq. (65) for the PSP *V* with *I*_*syn*_ described by eqs. (2)-(4) appropriately adapted to account for the translation of *V* to the equilibrium point, and STP described by eqs. (12) and (13) (DA model). The impedance amplitude (*Z*) and phase (Φ_*Z*_) were computed using eqs. (39) and (40). The analytical approximations for the PSP peak sequence response of passive cells to presynaptic inputs are described in Section2.2.4 (see also Appendix A). The approximation of *V*_*peak,n*_, *V*_*trough,n*_ and *t*_*V,peak*_ were computed as described in Section 2.2.4. The PSP amplitude Γ_*V*_ was computed by using eq. (41) and the PSP phase Φ_*V*_ was computed using eq. (42). The synaptic (*S*) peak (*S*_*peak*_) and phase (Φ_*S*_) profiles were computed similarly to these for *V*. We used the following additional parameter values: *C* = 1, *E*_*L*_ = −60, *I*_*app*_ = 0, *G*_*syn*_ = 0.1, *E*_*syn*_ = 0, *a*_*d*_ = 0.1, *a*_*f*_ = 0.1, *x*_∞_ = 1, *z*_∞_ = 0 and *T*_*sw*_ = 1.

**Figure S7:**
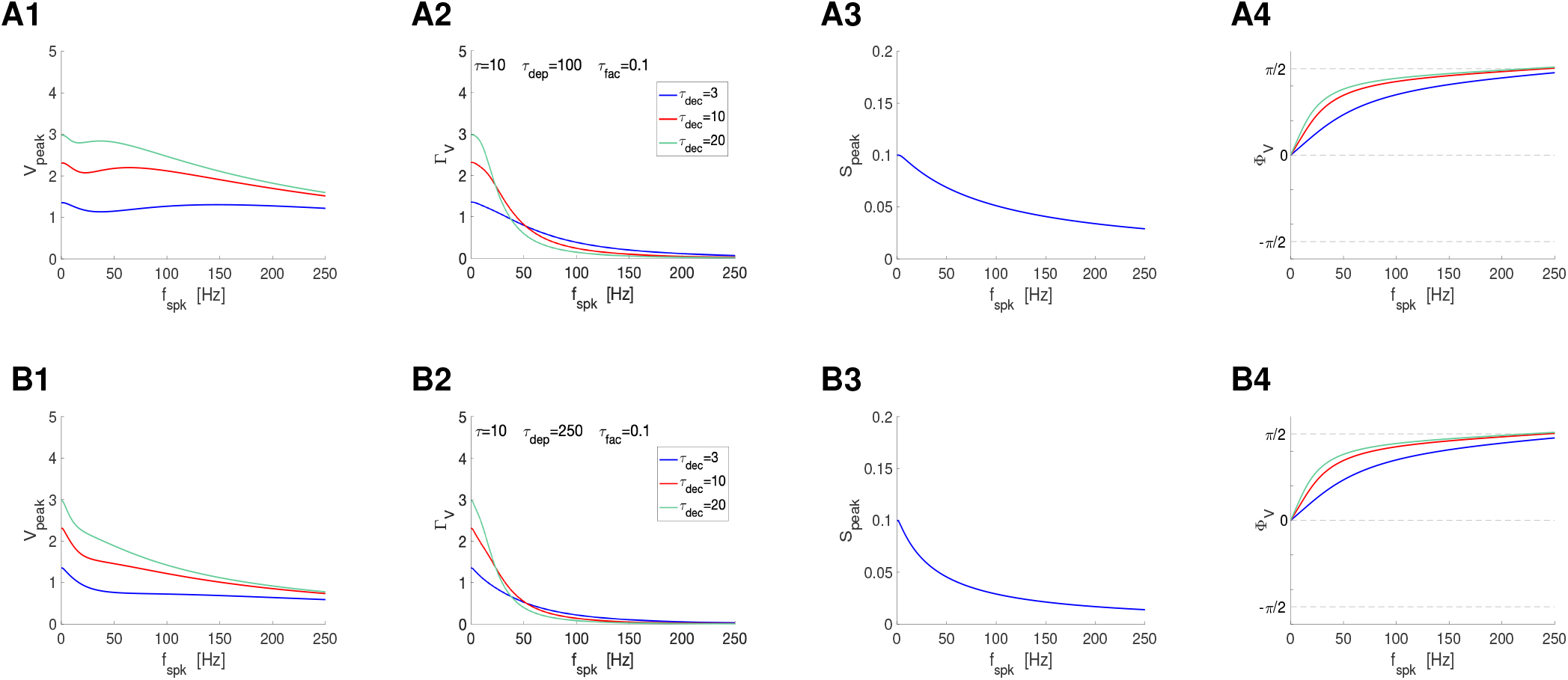
Postsynaptic filters in response to periodic presynaptic spike inputs emerging from the interplay of short-term depression and postsynaptic summation. Superimposed filters for representative values of the synaptic decay time constant *τ*_*dec*_. **A**. *τ*_*dep*_ = 100. **B**. *τ*_*dep*_ = 150. **A, B**. *τ* = 10 and *τ*_*fac*_ = 0.1. **Left column**. *V* peak profiles. **Middle-left column**. *V* peak-to-trough amplitude profiles. **Middle-right column**. *S* peak profiles. They are independent of *τ*_*dec*_. **Right column**. *V* phase profiles. We used eq. (65) for the PSP *V* with *I*_*syn*_ described by eqs. (2)-(4) appropriately adapted to account for the translation of *V* to the equilibrium point, and STP described by eqs. (12) and (13) (DA model). The impedance amplitude (*Z*) and phase (Φ_*Z*_) were computed using eqs. (39) and (40). The analytical approximations for the PSP peak sequence response of passive cells to presynaptic inputs are described in Section2.2.4 (see also Appendix A). The approximation of *V*_*peak,n*_, *V*_*trough,n*_ and *t*_*V,peak*_ were computed as described in Section 2.2.4. The PSP amplitude Γ_*V*_ was computed by using eq. (41) and the PSP phase Φ_*V*_ was computed using eq. (42). The synaptic (*S*) peak (*S*_*peak*_) and phase (Φ_*S*_) profiles were computed similarly to these for *V*. We used the following additional parameter values: *C* = 1, *E*_*L*_ = −60, *I*_*app*_ = 0, *G*_*syn*_ = 0.1, *E*_*syn*_ = 0, *a*_*d*_ = 0.1, *a*_*f*_ = 0.1, *x*_∞_ = 1, *z*_∞_ = 0 and *T*_*sw*_ = 1.

**Figure S8:**
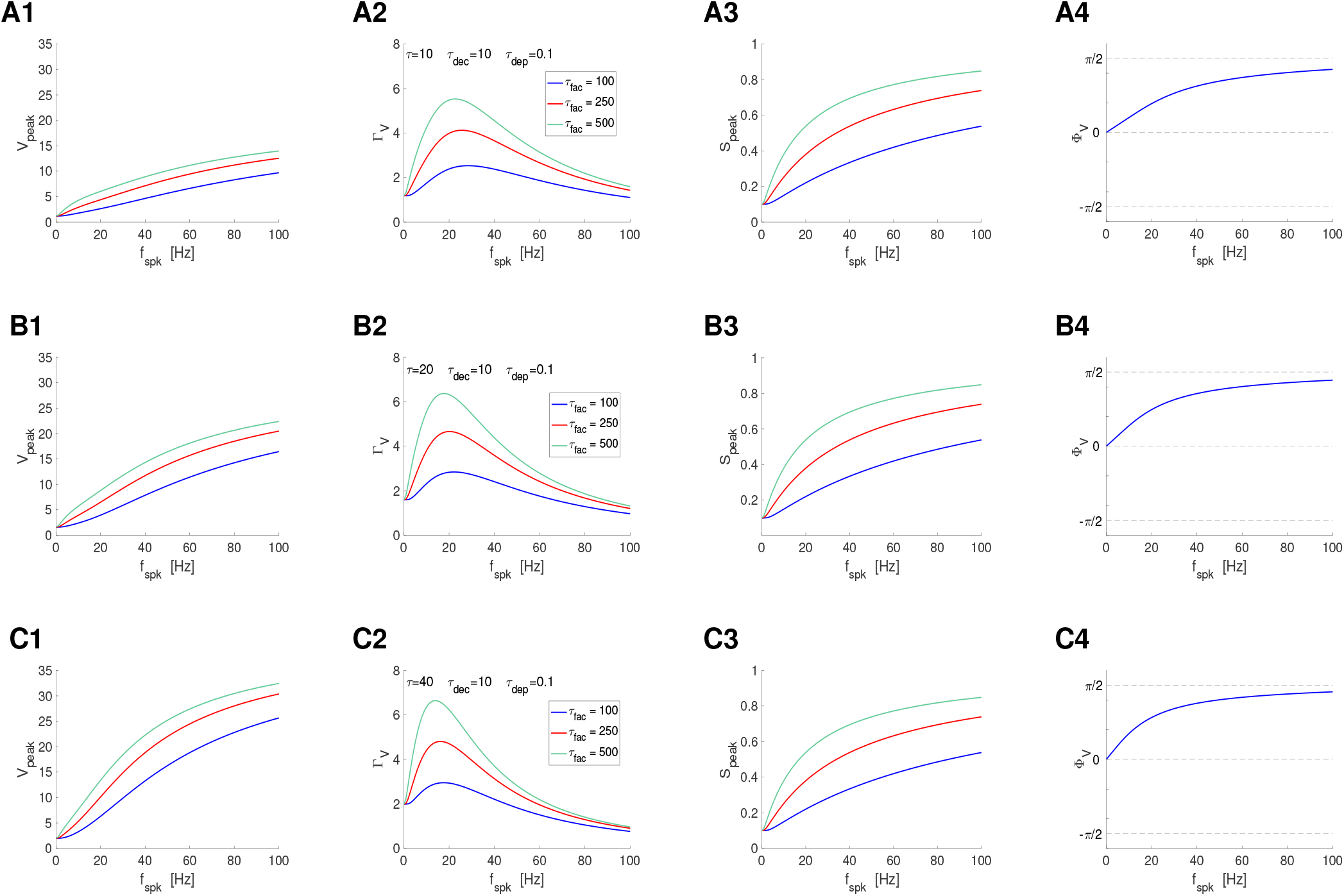
Postsynaptic filters in response to periodic presynaptic spike inputs emerging from the interplay of short-term facilitation and postsynaptic summation. Superimposed filters for representative values of the short-term facilitation time constant *τ*_*fac*_. **A**. *τ* = 10. **B**. *τ* = 20. **C**. *τ* = 40. **A, B, C**. *τ*_*dec*_ = 10 and *τ*_*dep*_ = 0.1. **Left column**. *V* peak profiles. **Middle-left column**. *V* peak-to-trough amplitude profiles. **Middle-right column**. *S* peak profiles. **Right column**. *V* phase profiles. We used eq. (65) for the PSP *V* with *I*_*syn*_ described by eqs. (2)-(4) appropriately adapted to account for the translation of *V* to the equilibrium point, and STP described by eqs. (12) and (13) (DA model). The impedance amplitude (*Z*) and phase (Φ_*Z*_) were computed using eqs. (39) and (40). The analytical approximations for the PSP peak sequence response of passive cells to presynaptic inputs are described in Section2.2.4 (see also Appendix A). The approximation of *V*_*peak,n*_, *V*_*trough,n*_ and *t*_*V,peak*_ were computed as described in Section 2.2.4. The PSP amplitude Γ_*V*_ was computed by using eq. (41) and the PSP phase Φ_*V*_ was computed using eq. (42). The synaptic (*S*) peak (*S*_*peak*_) and phase (Φ_*S*_) profiles were computed similarly to these for *V*. We used the following additional parameter values: *C* = 1, *E*_*L*_ = −60, *I*_*app*_ = 0, *G*_*syn*_ = 0.05, *E*_*syn*_ = 0, *a*_*d*_ = 0.1, *a*_*f*_ = 0.1, *x*_∞_ = 1, *z*_∞_ = 0 and *T*_*sw*_ = 1.

**Figure S9:**
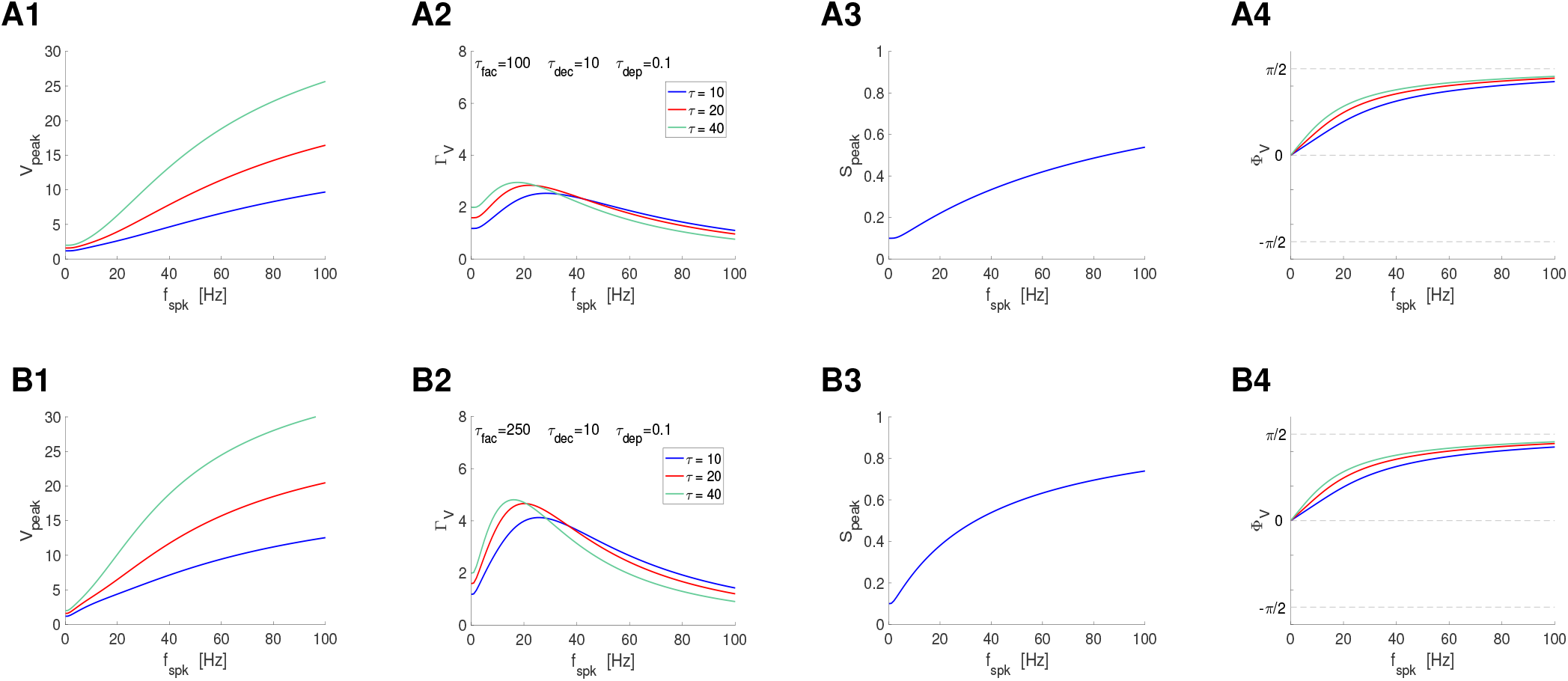
Postsynaptic filters in response to periodic presynaptic spike inputs emerging from the interplay of short-term facilitation and postsynaptic summation. Superimposed filters for various values of the membrane time constant *τ*. **A**. *τ*_*fac*_ = 100. **B**. *τ*_*fac*_ = 250. **A, B, C**. *τ*_*dec*_ = 10 and *τ*_*dep*_ = 0.1. **Middle-left column**. *V* peak-to-trough amplitude profiles. **Middle-right column**. *S* peak profiles. **Right column**. *V* phase profiles. We used eq. (65) for the PSP *V* with *I*_*syn*_ described by eqs. (2)-(4) appropriately adapted to account for the translation of *V* to the equilibrium point, and STP described by eqs. (12) and (13) (DA model). The impedance amplitude (*Z*) and phase (Φ_*Z*_) were computed using eqs. (39) and (40). The analytical approximations for the PSP peak sequence response of passive cells to presynaptic inputs are described in Section2.2.4 (see also Appendix A). The approximation of *V*_*peak,n*_, *V*_*trough,n*_ and *t*_*V,peak*_ were computed as described in Section 2.2.4. The PSP amplitude Γ_*V*_ was computed by using eq. (41) and the PSP phase Φ_*V*_ was computed using eq. (42). The synaptic (*S*) peak (*S*_*peak*_) and phase (Φ_*S*_) profiles were computed similarly to these for *V*. We used the following additional parameter values: *C* = 1, *E*_*L*_ = −60, *I*_*app*_ = 0, *G*_*syn*_ = 0.1, *E*_*syn*_ = 0, *a*_*d*_ = 0.1, *a*_*f*_ = 0.1, *x*_∞_ = 1, *z*_∞_ = 0 and *T*_*sw*_ = 1.

**Figure S10:**
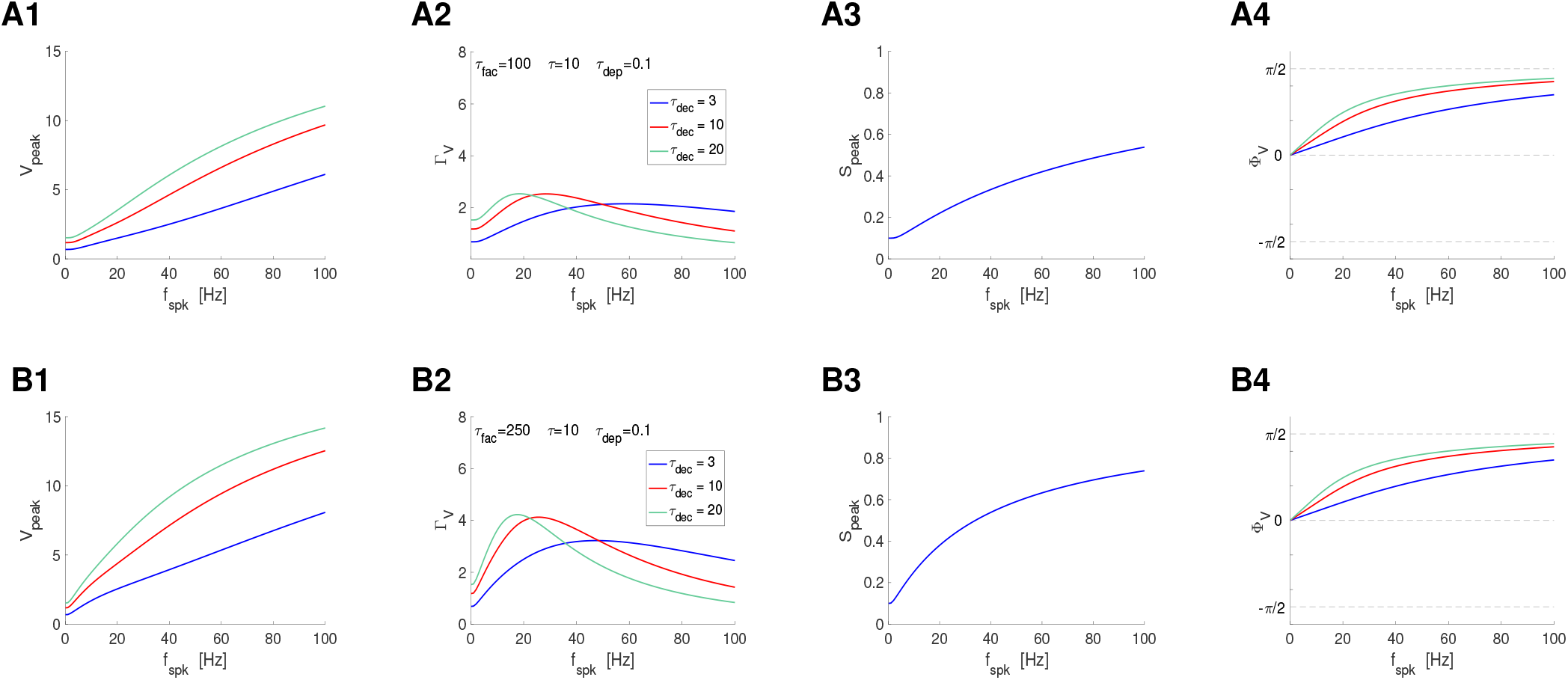
Postsynaptic filters in response to periodic presynaptic spike inputs emerging from the interplay of short-term depression and postsynaptic summation. Superimposed filters for representative values of the synaptic decay time constant *τ*_*dec*_. **A**. *τ*_*dep*_ = 100. **B**. *τ*_*dep*_ = 150. **A, B**. *τ* = 10 and *τ*_*fac*_ = 0.1. **Left column**. *V* peak profiles. **Middle-left column**. *V* peak-to-trough amplitude profiles. **Middle-right column**. *S* peak profiles. They are independent of *τ*_*dec*_. **Right column**. *V* phase profiles. We used eq. (65) for the PSP *V* with *I*_*syn*_ described by eqs. (2)-(4) appropriately adapted to account for the translation of *V* to the equilibrium point, and STP described by eqs. (12) and (13) (DA model). The impedance amplitude (*Z*) and phase (Φ_*Z*_) were computed using eqs. (39) and (40). The analytical approximations for the PSP peak sequence response of passive cells to presynaptic inputs are described in Section2.2.4 (see also Appendix A). The approximation of *V*_*peak,n*_, *V*_*trough,n*_ and *t*_*V,peak*_ were computed as described in Section 2.2.4. The PSP amplitude Γ_*V*_ was computed by using eq. (41) and the PSP phase Φ_*V*_ was computed using eq. (42). The synaptic (*S*) peak (*S*_*peak*_) and phase (Φ_*S*_) profiles were computed similarly to these for *V*. We used the following additional parameter values: *C* = 1, *E*_*L*_ = −60, *I*_*app*_ = 0, *G*_*syn*_ = 0.1, *E*_*syn*_ = 0, *a*_*d*_ = 0.1, *a*_*f*_ = 0.1, *x*_∞_ = 1, *z*_∞_ = 0 and *T*_*sw*_ = 1.

**Figure S11:**
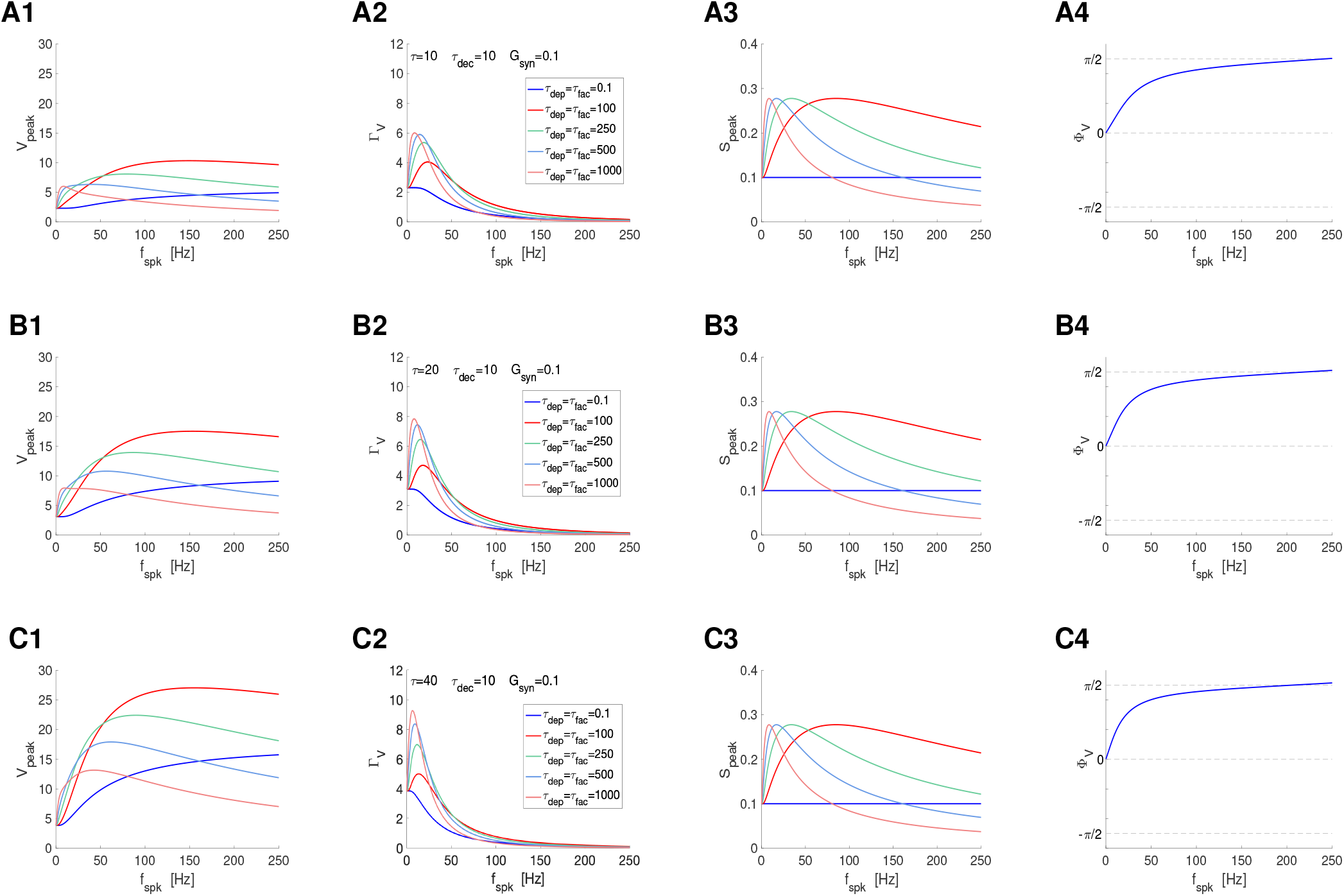
Postsynaptic filters in response to periodic presynaptic spike inputs emerging from the interplay of short-term depression, facilitation and postsynaptic summation. Superimposed filters for representative values of the depression and facilitation time constants *τ*_*dep*_ and *τ*_*fac*_, respectively. **A**. *τ* = 10. **B**. *τ* = 20. **C**. *τ* = 40. **A, B, C**. *τ*_*dec*_ = 10 and *G*_*syn*_ = 0.1. **Left column**. *V* peak profiles. **Middle-left column**. *V* peak-to-trough amplitude profiles. **Middle-right column**. *S* peak profiles. **Right column**. *V* phase profiles. We used eq. (65) for the PSP *V* with *I*_*syn*_ described by eqs. (2)-(4) appropriately adapted to account for the translation of *V* to the equilibrium point, and STP described by eqs. (12) and (13) (DA model). The impedance amplitude (*Z*) and phase (Φ_*Z*_) were computed using eqs. (39) and (40). The analytical approximations for the PSP peak sequence response of passive cells to presynaptic inputs are described in Section2.2.4 (see also Appendix A). The approximation of *V*_*peak,n*_, *V*_*trough,n*_ and *t*_*V,peak*_ were computed as described in Section 2.2.4. The PSP amplitude Γ_*V*_ was computed by using eq. (41) and the PSP phase Φ_*V*_ was computed using eq. (42). The synaptic (*S*) peak (*S*_*peak*_) and phase (Φ_*S*_) profiles were computed similarly to these for *V*. We used the following additional parameter values: *C* = 1, *E*_*L*_ = −60, *I*_*app*_ = 0, *G*_*syn*_ = 0.05, *E*_*syn*_ = 0, *a*_*d*_ = 0.1, *a*_*f*_ = 0.1, *x*_∞_ = 1, *z*_∞_ = 0 and *T*_*sw*_ = 1.

**Figure S12:**
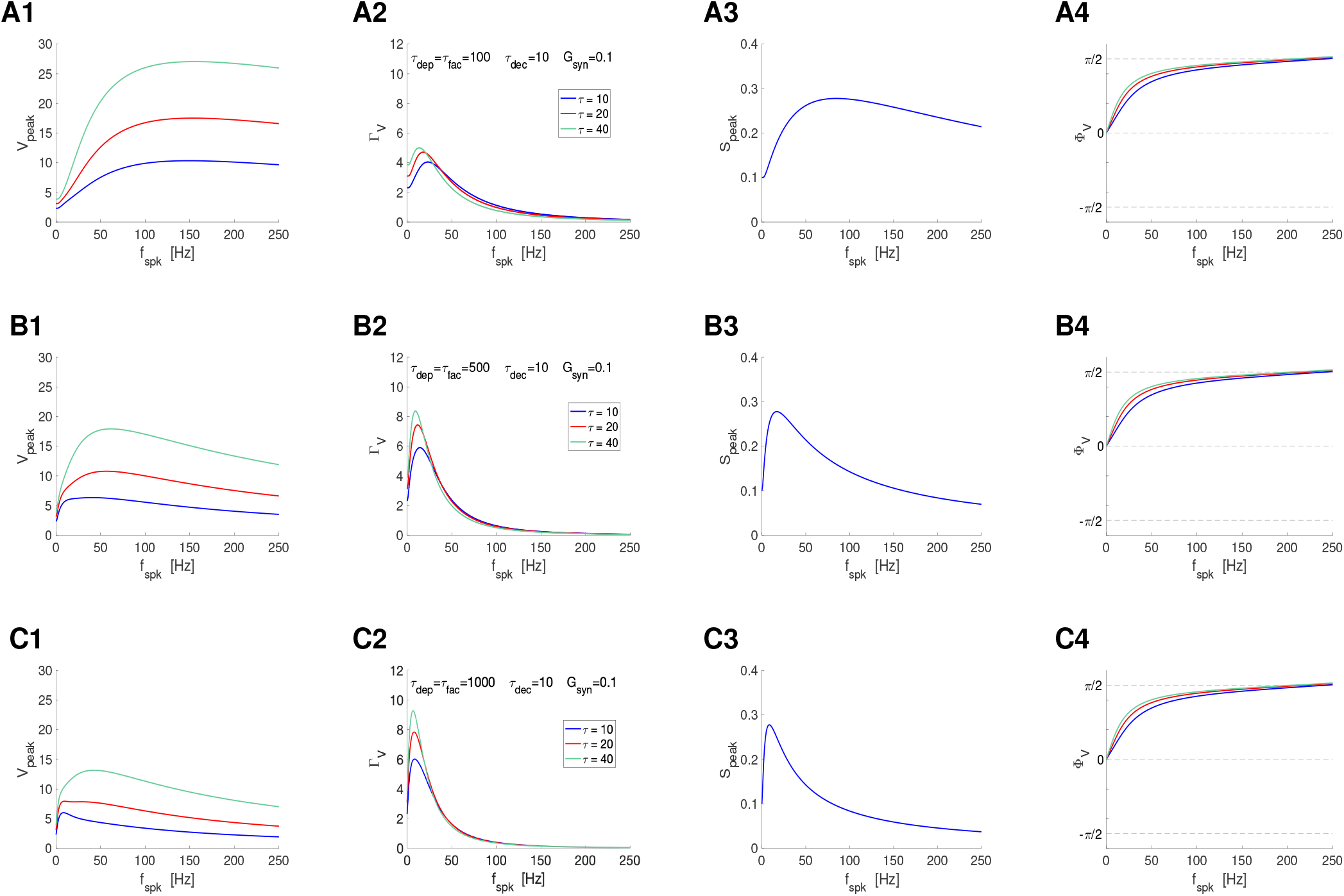
Postsynaptic filters in response to periodic presynaptic spike inputs emerging from the interplay of short-term depression, facilitation and postsynaptic summation. Superimposed filters for representative values of the membrane time constant *τ*. **A**. *τ*_*dep*_ = *τ*_*fac*_ = 100. **B**. *τ*_*dep*_ = *τ*_*fac*_ = 500. **C**. *τ*_*dep*_ = *τ*_*fac*_ = 1000. **A, B, C**. *τ*_*dec*_ = 10 and *G*_*syn*_ = 0.1. **Left column**. *V* peak profiles. **Middle-left column**. *V* peak-to-trough amplitude profiles. **Middle-right column**. *S* peak profiles. **Right column**. *V* phase profiles. We used eq. (65) for the PSP *V* with *I*_*syn*_ described by eqs. (2)-(4) appropriately adapted to account for the translation of *V* to the equilibrium point, and STP described by eqs. (12) and (13) (DA model). The impedance amplitude (*Z*) and phase (Φ_*Z*_) were computed using eqs. (39) and (40). The analytical approximations for the PSP peak sequence response of passive cells to presynaptic inputs are described in Section2.2.4 (see also Appendix A). The approximation of *V*_*peak,n*_, *V*_*trough,n*_ and *t*_*V,peak*_ were computed as described in Section 2.2.4. The PSP amplitude Γ_*V*_ was computed by using eq. (41) and the PSP phase Φ_*V*_ was computed using eq. (42). The synaptic (*S*) peak (*S*_*peak*_) and phase (Φ_*S*_) profiles were computed similarly to these for *V*. We used the following additional parameter values: *C* = 1, *E*_*L*_ = −60, *I*_*app*_ = 0, *G*_*syn*_ = 0.05, *E*_*syn*_ = 0, *a*_*d*_ = 0.1, *a*_*f*_ = 0.1, *x*_∞_ = 1, *z*_∞_ = 0 and *T*_*sw*_ = 1.

**Figure S13:**
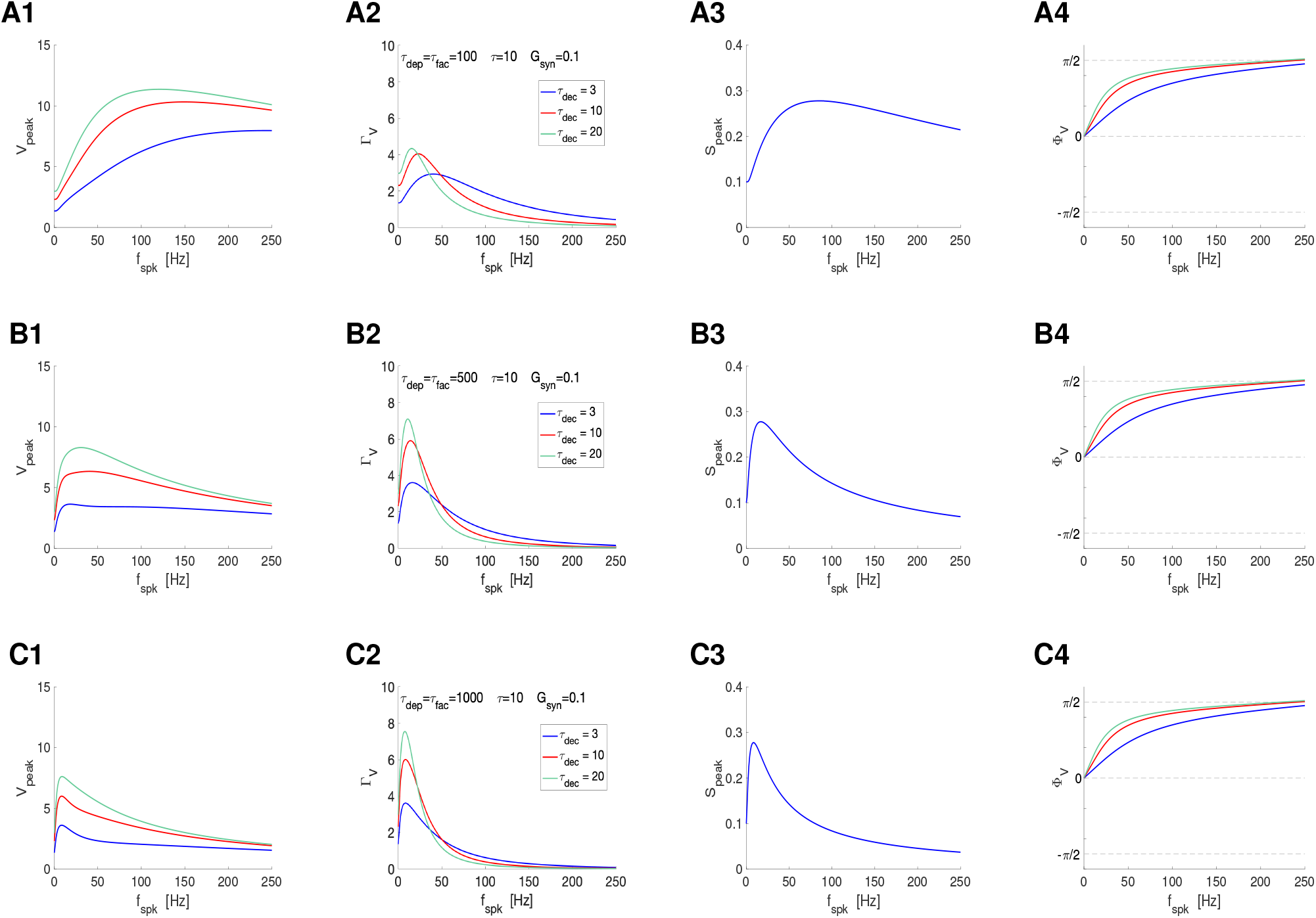
Postsynaptic filters in response to periodic presynaptic spike inputs emerging from the interplay of short-term depression, facilitation and postsynaptic summation. Superimposed filters for representative values of the depression synaptic decay time *τ*_*dec*_. **A**. *τ*_*dep*_ = *τ*_*fac*_ = 100. **B**. *τ*_*dep*_ = *τ*_*fac*_ = 500. **C**. *τ*_*dep*_ = *τ*_*fac*_ = 1000. **A, B, C**. *τ* = 10 and *G*_*syn*_ = 0.1. **Left column**. *V* peak profiles. **Middle-left column**. *V* peak-to-trough amplitude profiles. **Middle-right column**. *S* peak profiles. **Right column**. *V* phase profiles. We used eq. (65) for the PSP *V* with *I*_*syn*_ described by eqs. (2)-(4) appropriately adapted to account for the translation of *V* to the equilibrium point, and STP described by eqs. (12) and (13) (DA model). The impedance amplitude (*Z*) and phase (Φ_*Z*_) were computed using eqs. (39) and (40). The analytical approximations for the PSP peak sequence response of passive cells to presynaptic inputs are described in Section2.2.4 (see also Appendix A). The approximation of *V*_*peak,n*_, *V*_*trough,n*_ and *t*_*V,peak*_ were computed as described in Section 2.2.4. The PSP amplitude Γ_*V*_ was computed by using eq. (41) and the PSP phase Φ_*V*_ was computed using eq. (42). The synaptic (*S*) peak (*S*_*peak*_) and phase (Φ_*S*_) profiles were computed similarly to these for *V*. We used the following additional parameter values: *C* = 1, *E*_*L*_ = −60, *I*_*app*_ = 0, *G*_*syn*_ = 0.05, *E*_*syn*_ = 0, *a*_*d*_ = 0.1, *a*_*f*_ = 0.1, *x*_∞_ = 1, *z*_∞_ = 0 and *T*_*sw*_ = 1.

**Figure S14:**
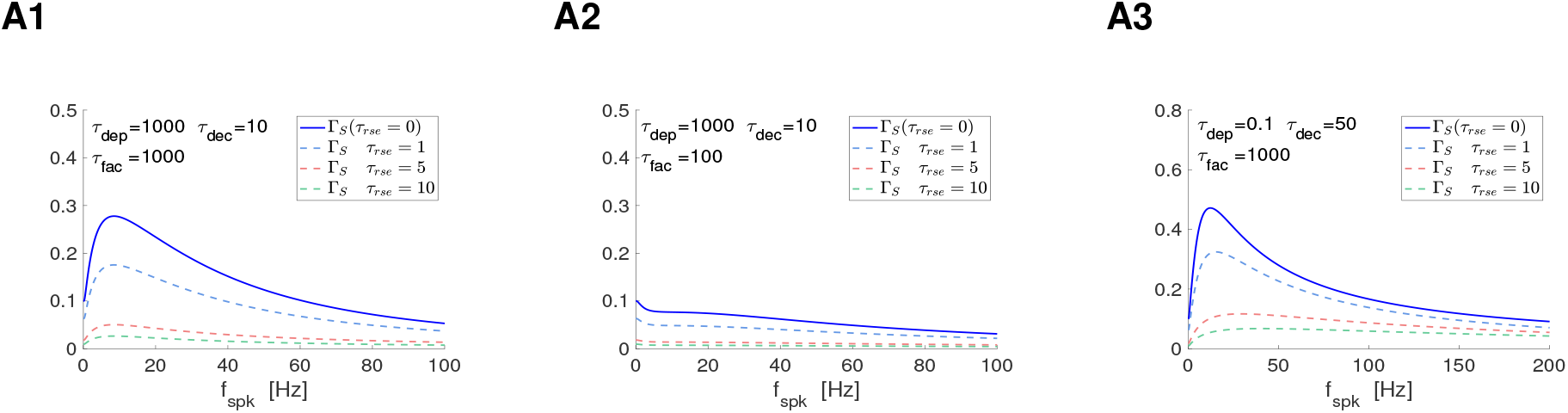
Γ_*S*_ filters in response to periodic presynaptic spike inputs (frequency *f*_*spk*_) for the to-Δ*S* update models with non-instantaneous update: representative examples. We used eqs. (12) and (13) (DA model) for 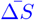 **A**. To-Δ*S* model (synaptic update to Δ*S*). Effects of *τ*_*rse*_. The *S* and Γ_*S*_ filters were computed using eqs. (53) and (54), respectively. **A1**. Γ_*S*_ band-pass filters attenuated by increasing values of *τ*_*rse*_. **A2**. Γ_*S*_ low-pass filters attenuated by increasing values of *τ*_*rse*_. **A3**. The Γ_*S*_ band-pass filter generated by the interplay of a Δ*S* high-pass filter and a *Q*_*A*_ (a low-pass filter) (Fig. 6-A6) is attenuated by increasing values of *τ*_*rse*_ and transitions to a high-pass filter. We used the following additional parameter values: *a*_*d*_ = 0.1, *a*_*f*_ = 0.1, *x*_∞_ = 1, *z*_∞_ = 0 and *T*_*sw*_ = 1.

**Figure S15:**
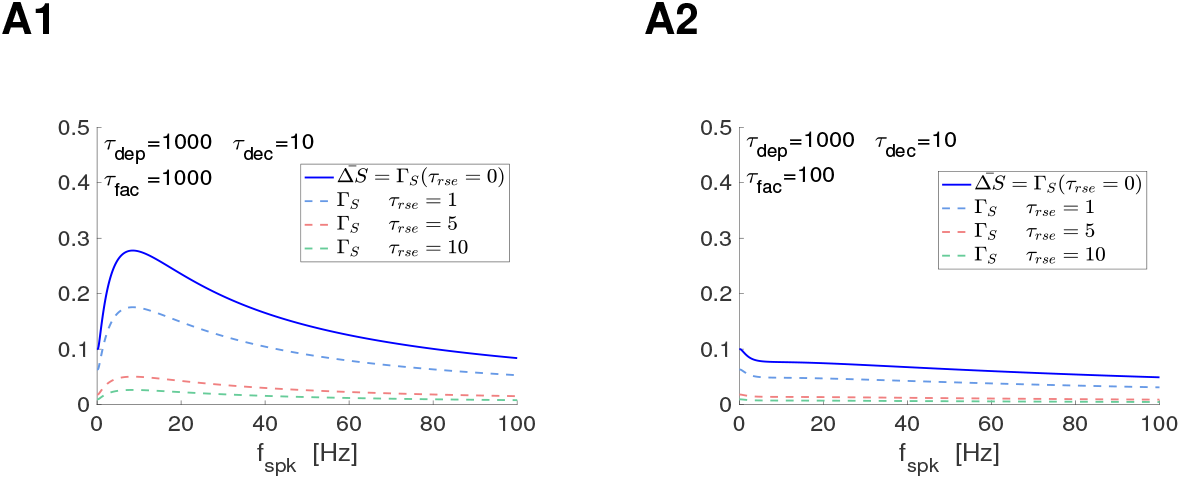
Γ_*S*_ filters in response to periodic presynaptic spike inputs (frequency *f*_*spk*_) for the by-Δ*S* update models with non-instantaneous update: representative examples. We used eqs. (12) and (13) (DA model) for 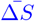 **A**. By-Δ*S* model (synaptic update to Δ*S*). Effects of *τ*_*rse*_. The *S* and Γ_*S*_ filters were computed using eqs. (62) and (63), respectively. **A1**. Γ_*S*_ band-pass filters attenuated by increasing values of *τ*_*rse*_. **A2**. Γ_*S*_ low-pass filters attenuated by increasing values of *τ*_*rse*_. We used the following additional parameter values: *a*_*d*_ = 0.1, *a*_*f*_ = 0.1, *x*_∞_ = 1, *z*_∞_ = 0 and *T*_*sw*_ = 1.

**Figure S16:**
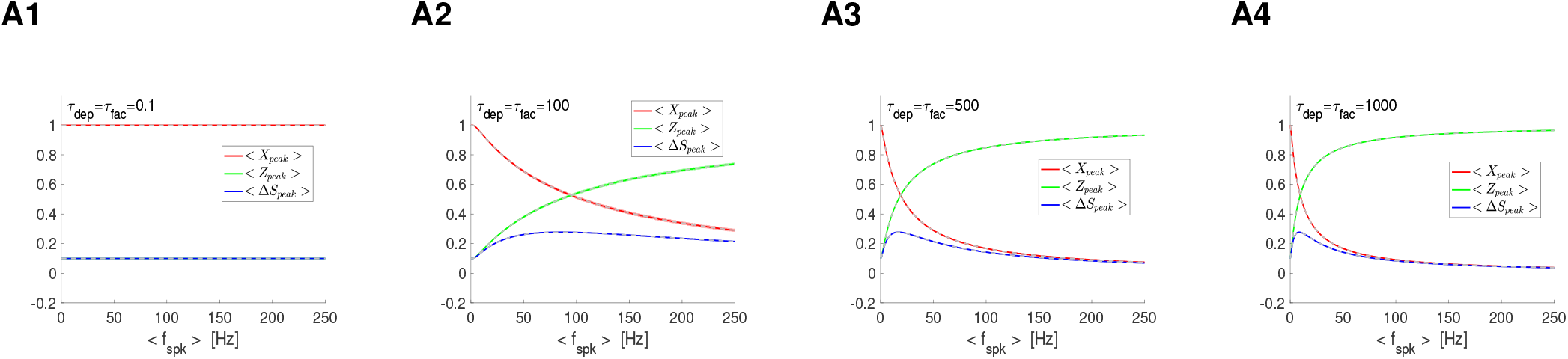
*X, Z* and Δ*S* filters in response to jittered (randomly perturbed) periodic presynaptic inputs in the presence of STP: frequency- and STP-dependent variability. For each value of the mean presynaptic input frequency *< f*_*spk*_ *>*, the ISI sequence *{*Δ_*spk,n*_*}* (*n* = 1, …, *N*_*spk*_) has the form Δ_*spk,n*_ = Δ_*spk*_ + *δ*_*spk,n*_ where Δ_*spk*_ is the ISI corresponding to *f*_*spk*_ (*f*_*spk*_ = 1000*/*Δ_*spk*_) and the sequence *{δ*_*spk,n*_*}* are drawn from a normal distribution with zero mean and variance equal to *δ* Δ_*spk*_. **A**. Superimposed *X*_*peak*_, *Z*_*peak*_ and Δ*S*_*peak*_ profiles for representative parameter values. We used *τ*_*dec*_ = 10 and *τ* = 10 in all panels. Solid curves correspond to the mean values for each attribute (*X*_*peak*_, *Z*_*peak*_ and Δ*S*_*peak*_). The shadow regions correspond to one standard deviation from the mean. The dashed gray curves, almost coinciding with the solid curves, represent the corresponding deterministic profiles (response to periodic spike train inputs with frequency *f*_*spk*_). **A1**. *τ*_*dep*_ = *τ*_*fac*_ = 0.1. **A2**. *τ*_*dep*_ = *τ*_*fac*_ = 100. **A3**. *τ*_*dep*_ = *τ*_*fac*_ = 500. **A4**. *τ*_*dep*_ = *τ*_*fac*_ = 1000. We used the following additional parameter values: *C* = 1, *E*_*L*_ = −60, *I*_*app*_ = 0, *E*_*syn*_ = 0, *a*_*d*_ = 0.1, *a*_*f*_ = 0.1, *x*_∞_ = 1, *z*_∞_ = 0 and *T*_*sw*_ = 1.

**Figure S17:**
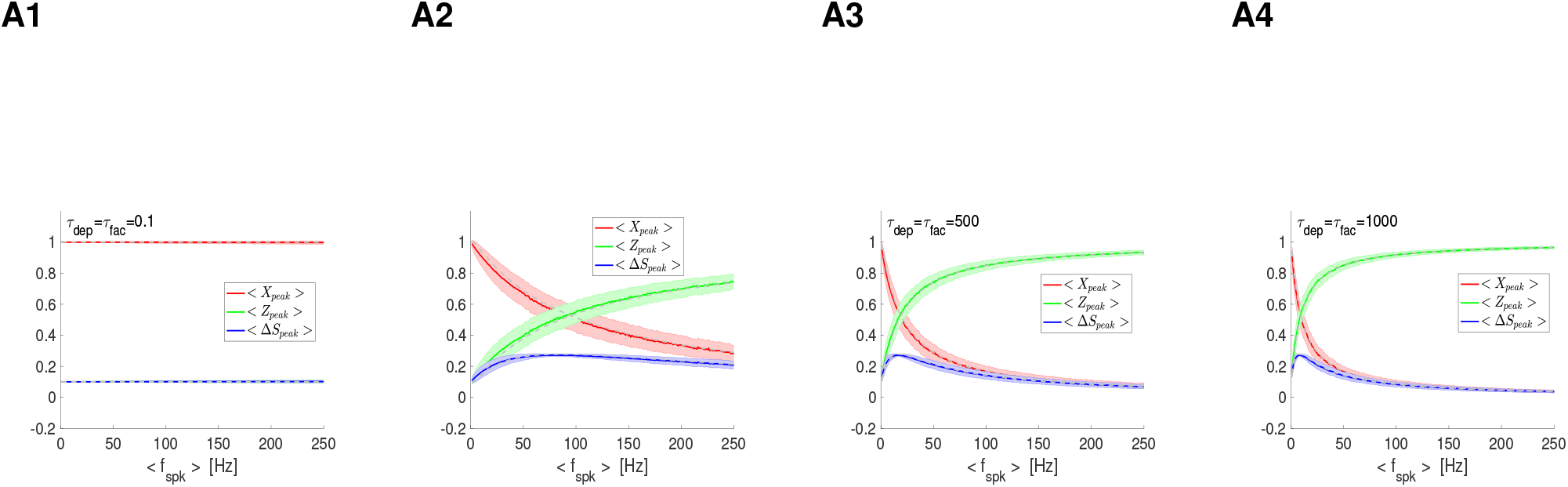
*X, Z* and Δ*S* filters in response to Poisson-distributed presynaptic inputs in the presence of STP: frequency- and STP-dependent variability. The mean rate of the Poisson distributed spike trains corresponds to *< f*_*spk*_ *>*. Superimposed *X*_*peak*_, *Z*_*peak*_ and Δ*S*_*peak*_ profiles for representative parameter values. We used *τ*_*dec*_ = 10 and *τ* = 10 in all panels. Solid curves correspond to the mean values for each attribute (*X*_*peak*_, *Z*_*peak*_ and Δ*S*_*peak*_). The shadow regions correspond to one standard deviation from the mean. The dashed curves represent the corresponding deterministic profiles (response to periodic spike train inputs with frequency *f*_*spk*_). **A1**. *τ*_*dep*_ = *τ*_*fac*_ = 0.1. **A2**. *τ*_*dep*_ = *τ*_*fac*_ = 100. **A3**. *τ*_*dep*_ = *τ*_*fac*_ = 500. **A4**. *τ*_*dep*_ = *τ*_*fac*_ = 1000. We used the following additional parameter values: *C* = 1, *E*_*L*_ = −60, *I*_*app*_ = 0, *E*_*syn*_ = 0, *a*_*d*_ = 0.1, *a*_*f*_ = 0.1, *x*_∞_ = 1, *z*_∞_ = 0 and *T*_*sw*_ = 1.

